# Identifying the Link Between Chemical Exposures and Breast Cancer in African American Women via ToxCast High Throughput Screening Data

**DOI:** 10.1101/2021.01.22.427848

**Authors:** Katelyn Polemi, Vy Nguyen, Julien Heidt, Adam Kahana, Olivier Jolliet, Justin A. Colacino

**Affiliations:** Department of Environmental Health Sciences, University of Michigan, Ann Arbor, MI, USA; Department of Computational Medicine and Bioinformatics, University of Michigan, Ann Arbor, MI, USA; Center for Computational Medicine and Bioinformatics, University of Michigan, Ann Arbor, MI, USA; Department of Nutritional Sciences, University of Michigan, Ann Arbor, MI, USA

## Abstract

Among women, breast cancer is the most prevalent form of cancer worldwide and has the second highest mortality rate of any cancer in the United States. The breast cancer related death rate is 40% higher in African American women compared to European American women. The incidence of triple negative breast cancer (TNBC), an aggressive subtype of breast cancer for which there is no targeted therapy, is approximately three times higher in non-Hispanic Black women (NHBW) compared to non-Hispanic White women (NHWW). The drivers of these differences in breast cancer incidence and mortality are still poorly understood, and likely lie in an interaction between genetic and environmental factors. Here, we aimed to identify chemical exposures which may play a role in breast cancer disparities. Using chemical biomonitoring data from the National Health and Nutrition Examination Survey (NHANES) and biological activity data from the EPA’s ToxCast program, we assessed the toxicological profiles of chemicals with higher biomarker concentrations in US NHBW. We conducted a literature search to identify a gene set of breast cancer targets included in ToxCast to analyze the response of prioritized chemicals in these assays. Forty-three chemical biomarkers are significantly higher in NHBW. Investigation of these chemicals in ToxCast resulted in a total of 32,683 assays for analysis, 5,172 of which contained nonzero values for the concentration at which the dose-response fitted model reaches the cutoff considered “active” and the scaled top value of dose response curve. Of these chemicals BPA, PFOS, and thiram are most comprehensively assayed. 2,5-dichlorophenol, 1,4-dichlorobenzene, and methyl and propyl parabens had higher biomarker concentrations in NHBW and moderate testing and activity in ToxCast. The distribution of active concentrations for these chemicals in ToxCast assays are comparable to biomarker concentrations in NHBW. Through this integrated analysis, we have identified that multiple chemicals, including thiram, propylparaben, and p,p’ DDE, with disproportionate exposures in NHBW, have breast cancer associated biological activity at human exposure relevant doses.

## 1. Introduction

Breast cancer accounts for more than 1 in 10 new cancer diagnoses each year and is the second leading cause of cancer-related mortality in women (Simon and Robb, 2014; Power *et al*., 2018; Siegel *et al*., 2019). Breast cancer outcomes, however, are significantly different across women of different races/ethnicities. African American women are 40% more likely to die from breast cancer than women of any other race (Williams *et al*., 2016). Furthermore, the incidence of triple negative breast cancer (TNBC), an aggressive subtype of breast cancer for which there is no targeted therapy, is approximately three times higher in non-Hispanic Black women compared to non-Hispanic White women (Carey *et al*., 2006; Stark *et al*., 2010). Relative to non-Hispanic White women, non-Hispanic Black women are also 2.4 times more likely to die of breast cancer after being diagnosed with the pre-invasive lesion, ductal carcinoma *in situ* (Narod *et al*., 2015).

The mechanisms driving these differences in breast cancer outcomes are likely due to interactions between genetic and environmental factors, although these interactions are complex and remain poorly understood. Genetic epidemiology cohort studies have identified a small number of genetic polymorphisms linked to breast cancer in African American women (Murphy *et al*., 2017; Feng *et al*., 2017). However, the contribution of genetic variations appear minor in explaining these cancer disparities (Braun, 2002; Diez Roux, 2012; Cooper *et al*., 2003; Jing *et al*., 2014; Rodgers *et al*., 2018). A number of additional non-genetic contributing factors such as disparities in income, barriers to screening, differences in treatment, and higher stage of disease at diagnosis have also been identified (Dietze *et al*., 2015). The role of chemical exposures has been less explored in the context of racial disparities in breast cancer outcomes; however, differences in chemical exposures have been hypothesized to be important etiologic factors in racial disparities for multiple diseases (Hoover *et al*., 2012; Juarez and Matthews-Juarez, 2018; Ruiz *et al*., 2018; Wang *et al*., 2016; Zota and Shamasunder, 2017). Mounting evidence from toxicology and epidemiology studies, and an increased understanding of the mechanisms linking toxicant exposures with breast cancer, suggests that many common chemical exposures may alter breast cancer risk (Gray *et al*., 2017).

We recently have examined racial disparities in chemical biomarker concentrations in US women. Using data from the National Health and Nutrition Examination Survey (NHANES), an ongoing population-based health study conducted by the US Centers for Disease Control and Prevention, we identified stark differences in chemical exposure biomarker concentrations across women of various racial/ethnic groups (Nguyen *et al*., 2020). We identified a set of chemicals with concentrations significantly higher, on average, in African American women including pesticide metabolites (2,5-dichlorophenol, 1,4-dichlorobenzene), chemicals of personal care products (methyl paraben, propyl paraben, monoethyl phthalate), and heavy metals (mercury and lead). Here, our goal is to assess the biological activity of these chemicals identified as highly disparate in non-Hispanic Black women using data from the U.S Environmental Protection Agency’s (EPA) ToxCast program (now CompTox). Numerous studies have highlighted the utility of ToxCast data to identify toxicant effects on cell stress and cytotoxicity (Judson *et al*., 2016) and cancer (Iyer *et al*., 2019), as well as to inform adverse outcome pathway development (Fay *et al*., 2018a). In this study, we integrate human population chemical biomarker concentrations from NHANES with biological activity data from ToxCast. We identify a suite of chemicals with high biologic activity and significantly substantial exposure in African American women for further toxicologic and epidemiologic assessment for their relationship to breast cancer disparities. We hypothesize that chemicals found most disparate will show high biological activity in breast cancer related assays.

## 2. Methods

In the current study, we integrate chemical biomonitoring data from NHANES and biological activity data from the EPA’s ToxCast program to assess the toxicological relevance to breast cancer of chemicals with higher exposures in non-Hispanic Black women.

**Figure 1** shows a graphical workflow of the overall approach: Using NHANES biomarker data, we first identified chemicals with higher concentrations in Non-Hispanic Black Women. We extracted the high throughput toxicity data from the ToxCast database and combined with the NHANES exposure data. We compared exposed and bioactive concentrations and identify whether chemicals found at higher concentration in that subpopulation have the potential to activate known breast cancer genes.

**Figure 1:**
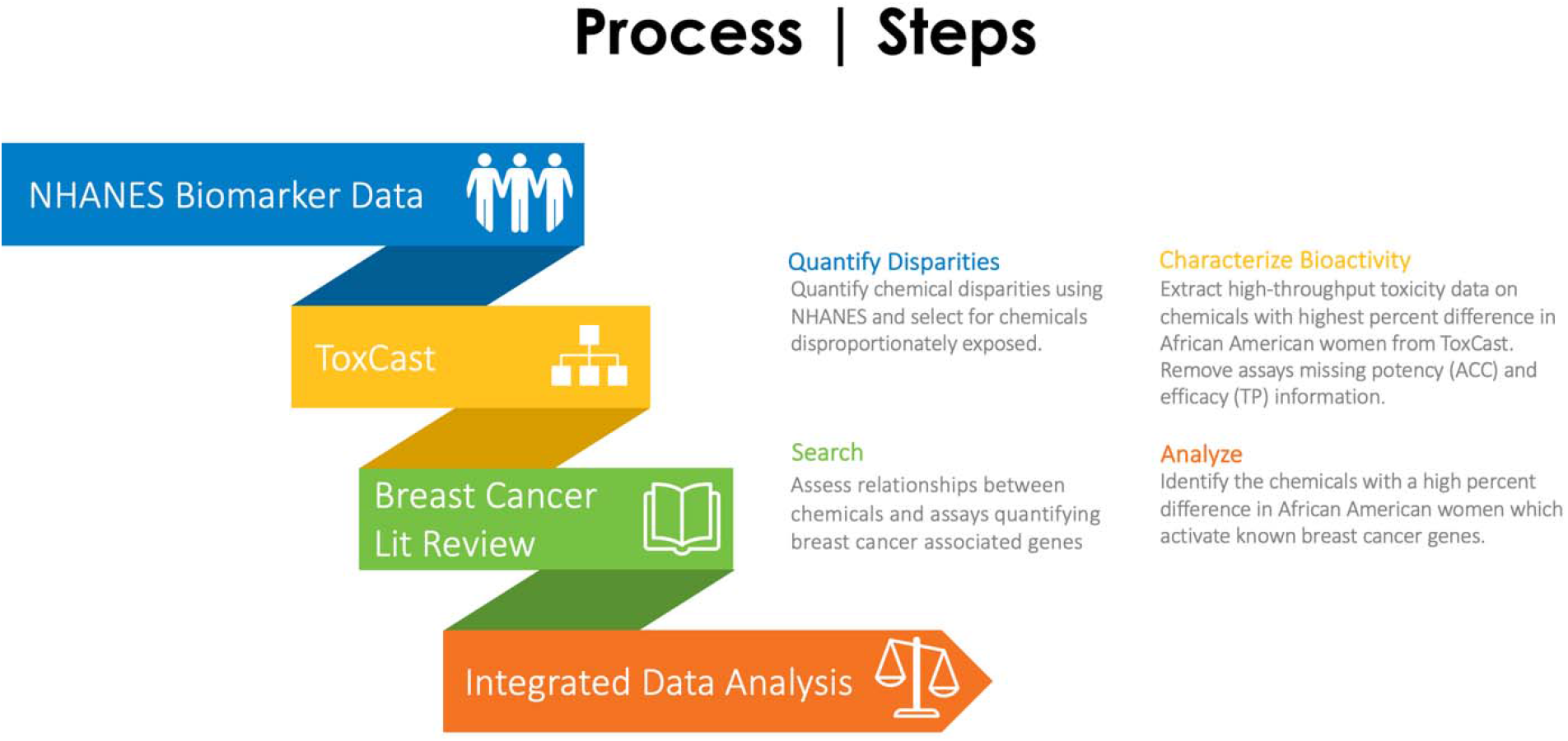
A Comprehensive Workflow for the Methodology of this Study. TP (modl_tp: scaled top of winning dose response curve; referred to as “TP”) and ACC (modl_acc: the concentration at which the dose-response fitted model reaches the cutoff considered “active”; referred to as “ACC”) are used as measures of efficacy and potency, respectively

### 2.1 Top 43 Chemicals Biomarkers Higher in Non-Hispanic Black Women

We previously identified chemical biomarkers disproportionately found in non-Hispanic Black women compared to non-Hispanic White women in NHANES (Nguyen *et al*., 2020). NHANES is a cross-sectional study designed to collect data on demographic, socioeconomic, dietary, and health-related characteristics in the non-institutionalized, civilian US population. The analysis used the continuous data on chemical biomarkers and demographics, collected from 1999-2014. The analysis focused on characterizing the chemical disparities in US women. Briefly, using a generalized linear regression-based approach, we quantified the relative difference in biomarker concentration between non-Hispanic Black and non-Hispanic White women across 143 chemicals, 43 of which are higher in non-Hispanic Black women. Some metabolites detected in NHANES had insufficient ToxCast data, therefore parent compounds are used in place. When measured, some chemicals, such as persistent organic pollutants, partition into the lipid component of an individual’s blood. Therefore, for these specific chemicals, the lipid-adjusted concentration is used to take this effect into account. For the chemicals found to be disproportionately higher in non-Hispanic Black women, we extracted high-throughput chemical assay data from the ToxCast database (version invitroDBv3.1) for further analysis in R. Bisphenol A (BPA), though not significantly higher in non-Hispanic Black women, is included in the analysis as a model chemical due to the large number of assays in which it has been tested and the association between BPA exposure and breast cancer (Wang *et al*., 2017; Zhang *et al*., 2015; Clément *et al*., 2017). Metals elevated in non-Hispanic Black women, such as lead or mercury, are considered in their biologically relevant forms. For example, assay data for Triphenyllead acetate and lead (II) acetate trihydrate are extracted from the ToxCast database rather than elemental lead. NHANES chemicals and their corresponding chemicals extracted from ToxCast can be found in **Supplemental Table 1**.

### 2.2 Collection of ToxCast Assays for Analysis

The publicly available ToxCast database offers high throughput in vitro toxicity information generated for over 8,000 different chemicals and over 1,000 biological endpoints (Brunner *et al*., 2019). Through screening a large chemical library, ToxCast aims to identify biological pathways relevant for response to toxic stressors, develop high throughput screening (HTS) assays for these pathways, generate comprehensive chemical dose-response data for these HTS assays, identify points of departure from the HTS data, and link HTS results to adverse effects. These HTS assays expose living cells, isolated proteins, and other biological molecules to chemicals to screen for biological activity and thus assess toxic effects of chemicals. Resulting endpoints from these assays indicate quantifiable effects linked to known biological processes, which are then used to determine the prioritization level of the chemical.

We extract publicly available ToxCast summary assay information for the 43 chemicals with available ToxCast data out of the top 50 chemical biomarkers that are higher in Non-Hispanic Black Women. Assay information includes intended target family, defined as the target family of the objective target for the assay such as ‘cytokine,’ and the intended target gene name/symbol. From the composite ToxCast assay results dataset, TP (modl_tp: scaled top of winning dose response curve; referred to as “TP”) and ACC (modl_acc: the concentration at which the dose-response fitted model reaches the cutoff considered “active”; referred to as “ACC”) are used as measures of efficacy and potency, respectively. Any assay-chemical pairs with missing TP or ACC values are included for the purpose of generating summary statistics but excluded from all subsequent analyses. The hitcall variable indicates whether the assay is active or inactive. Chemicals are first analyzed based on a ratio of active (defined as hitcall=1) to total assays. To understand how ToxCast considered a dose-response curve to have an active hitcall, we first need to understand the different models. The Hill or Gain-Loss models represent the dose-response relationship by minimizing a prediction performance measure, the Akaike Information Criterion (AIC). The Constant model is defined as the efficacy cutoff. Finally, a dose-response series must meet the following criteria to have an active hitcall: the Hill or Gain-Loss model must have a better prediction performance compared to the Constant model (i.e. a lower AIC), the top of the modeled curve must exceed the efficacy cutoff, and the median response value must also exceed the efficacy cutoff for at least one concentration (EPA, 2018). The Toxcast data processing pipeline incorporates the use of data quality flags which alerts the users to potential technical or statistical concerns such as false activity or noisy data (Ryan, N. and Becker, R., 2017). Because of limited data and varying degrees of flag severity, consistent with Judson et al. 2016, no assays were omitted due to flags.

### 2.3 Identifying breast cancer relevant assays in ToxCast

After a first analysis of all Toxcast assay, assay data is further filtered by selecting assays with Intended Target Family Genes considered to play a role in breast cancer. We conducted a review of breast cancer literature using Google Scholar to identify these target genes of interest. Keywords and phrases used for the literature review included, “molecular genetics of breast cancer,” “molecular characteristics of breast cancer,” “genes involved in TNBC pathogenesis,” “genes involved in breast cancer metastasis,” “genes involved in TNBC metastasis,” “TNBC and epithelial-mesenchymal transition,” “molecular mechanism of breast cancer,” “molecular pathways of breast cancer,” “molecular pathways of TNBC,” and “Molecular characteristics of TNBC”. Additionally, incorporated genes the BCScreen breast carcinogenesis gene panel from (Grashow *et al*., 2018) with assays in Toxcast. A list of 99 genes were produced from the literature search with the requirement that they also had to be tested in ToxCast. **Supplemental table 2** defines the function of each gene and provides references to support the relationship of this gene with breast cancer. The genes identified are involved in critical processes such as immune response, cell cycle regulation, epithelial/mesenchymal transition (EMT), metastasis, and DNA repair mechanisms. This list is used to understand the activity of ToxCast assays with breast cancer (BC) gene targets to assess the prioritization of our 43 chemicals and further analyze the mechanism of action of these chemicals in breast cancer. ToxCast gene symbols are all converted to human gene symbols. We performed unsupervised hierarchical clustering to visualize and group chemicals with similar activation patterns for gene specific assays. Additionally, using ACC and TP, assay activity and efficacy for all assays with breast cancer gene targets were plotted for each individual chemical.

### 2.4 Evaluating Relevant Chemical Concentrations

Chemical biomarker concentrations in non-Hispanic Black women in NHANES were converted to molarity units to compare to ToxCast assay ACC concentrations in order to understand the biological activity of human relevant exposure doses. We quantified the overlap in concentrations of our three studied datasets: NHANES, ToxCast, and ToxCast BC. We first defined all values between the minimum and maximum concentrations incremented by 0.01 uM for each dataset and then determined the intersecting values across the three different datasets. Finally, we calculated the range of the intersected values to obtain the overlap in concentrations.

## 3. Results

### Fraction of active assays in chemicals with disparities

Our previous analyses of chemical biomarker concentrations measured in NHANES identified a suite of chemical biomarkers significantly elevated in in non-Hispanic Black women (Nguyen *et al*., 2020). To characterize the biological activity of these chemical exposures and identify chemicals with both high bioactivity (fraction active > 0.1) and high difference between black and white women (difference>50%), we linked these exposures to ToxCast data by using the fraction of active assays, i.e., the ratio of the active assays to total ToxCast assays performed per chemical, a surrogate readout of biological activity. **Figure 2A** presents the biological activity of chemicals found disproportionately in African American women by displaying the relative percent difference of chemical biomarker concentrations detected in NHBW compared to NHWW in NHANES as a function of the fraction of active breast cancer related assays. On average, 2,5-Dichlorophenol is found more than 3-times higher in non-Hispanic Black women compared to non-Hispanic White women. 1,4-Dichlorobenzene is found slightly more than 2-times higher in non-Hispanic Black women, and methylparaben, propylparaben, and 2,4-Dichlorophenol are more than twice as high in non-Hispanic Black women. In addition, BPS, p,p’-DDE, o,p’-DDT, and o,p’-DDE are all found to be nearly 2-times higher. Of the chemicals that are approximately twice as high in non-Hispanic Black women, p,p’-DDE, o,p’-DDT, o,p’-DDE, propylparaben, and BPS have higher ratios of active assays to total assays compared to the other 37 chemicals in ToxCast. **Supplemental Table 3** shows the biological activity of chemicals disproportionately found in NHBW.

**Figure 2:**
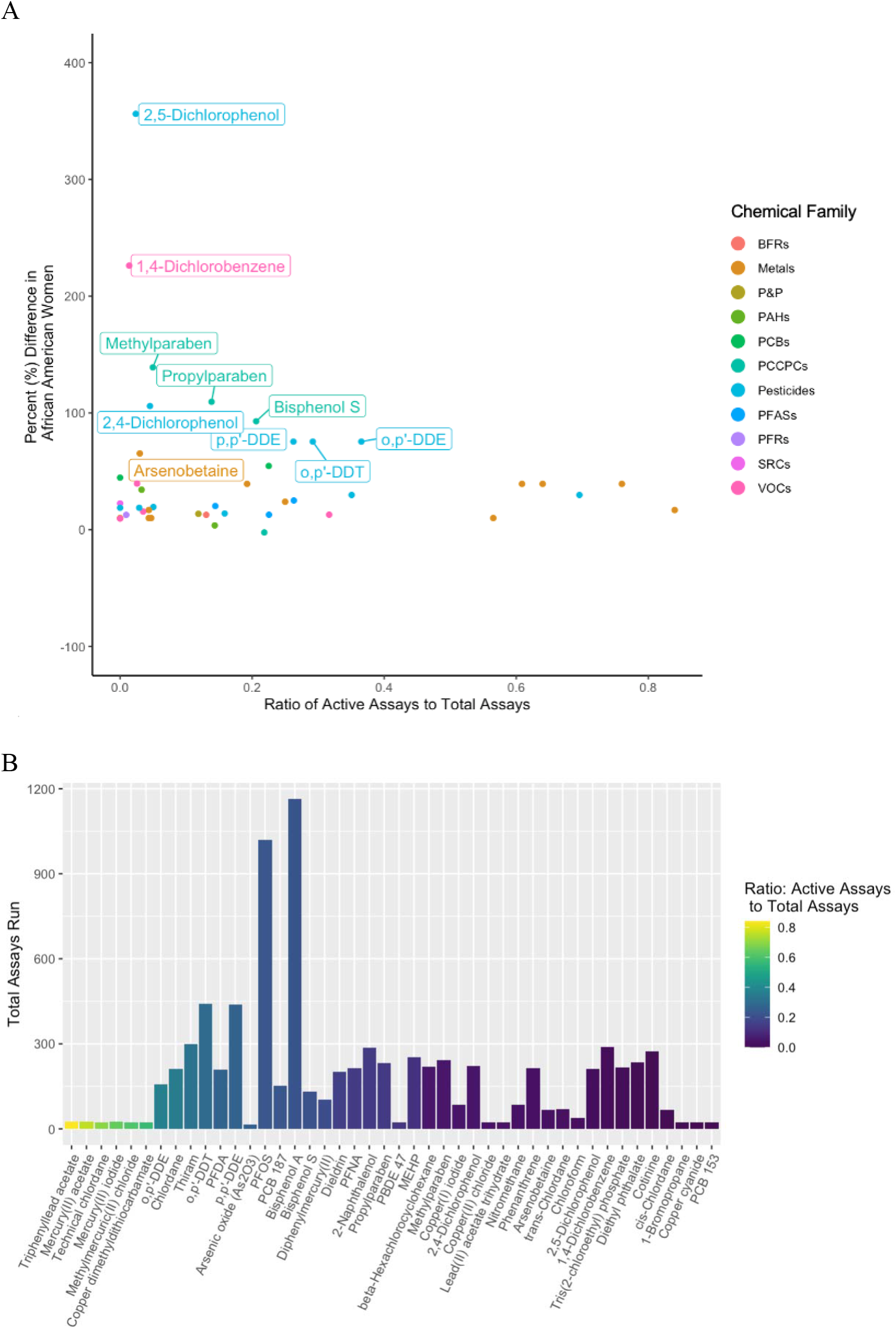
**A) Breast Cancer Related Biological Activity of Chemicals Disproportionately Found in African American Women.** Ratio of active (hitcall=1) to total breast cancer related assays compared with percent (%) difference between non-Hispanic Black women and non-Hispanic White women. **B: Active to Total Assays Testing Breast Cancer Genes.** Visualization of total number of assays run for each chemical colored by ratio of active to total assays.

Next, data exploration and visualization are performed across all breast cancer relevant assays obtained from ToxCast; no assays are excluded based on missing data or flags. **Figure 2B** shows that metals such as lead, mercury and copper in addition to the pesticide chlordane activate the most breast cancer related assays in ToxCast. Additionally, those chemicals between o,p’-DDE and MEHP show a range of activation of breast cancer related assays indicating their potential role in mechanisms underlying breast cancer. **Supplemental Figure 1** visually represents the number of assays run for each of the 43 chemicals in ToxCast. A total of 32,683 assays are available for analysis, 5,172 of which contained nonzero values for the concentration at which the dose-response fitted model reaches the cutoff considered “active” and the scaled top value of dose response curve. The color indicates the ratio of active to total assays run for each chemical, yellow representing a high ratio of activity and dark blue representing a lower ratio of activity.

### Visualization and grouping of chemicals with similar gene activation patterns

We next created a heatmap to visualize the unbiased hierarchal clustering of ToxCast assays for breast cancer (BC) genes based on ratio of active to total assays run per chemical (**Figure 3**) to group chemicals and genes by biological activity. Grey boxes indicate no assays are run for that chemical with that specific gene target. White indicates assays are run, but none are active. The ratio of activity ranges from 0.0-1.0. Genes most frequently tested with highest biological activity across chemicals of interest include *TP53*, *PPARG*, *ESR1*, *AR*, and *HIF1A*. Chemicals grouped in the same node as our model chemical, BPA, include PFOS, chlordane, thiram, p,p’-DDE and 2-naphthalenol. The metals all cluster near each other, demonstrating similar activation profiles. Interestingly, o,p’-DDE clusters far away from p,p’-DDE and o,p’-DDT.

**Figure 3:**
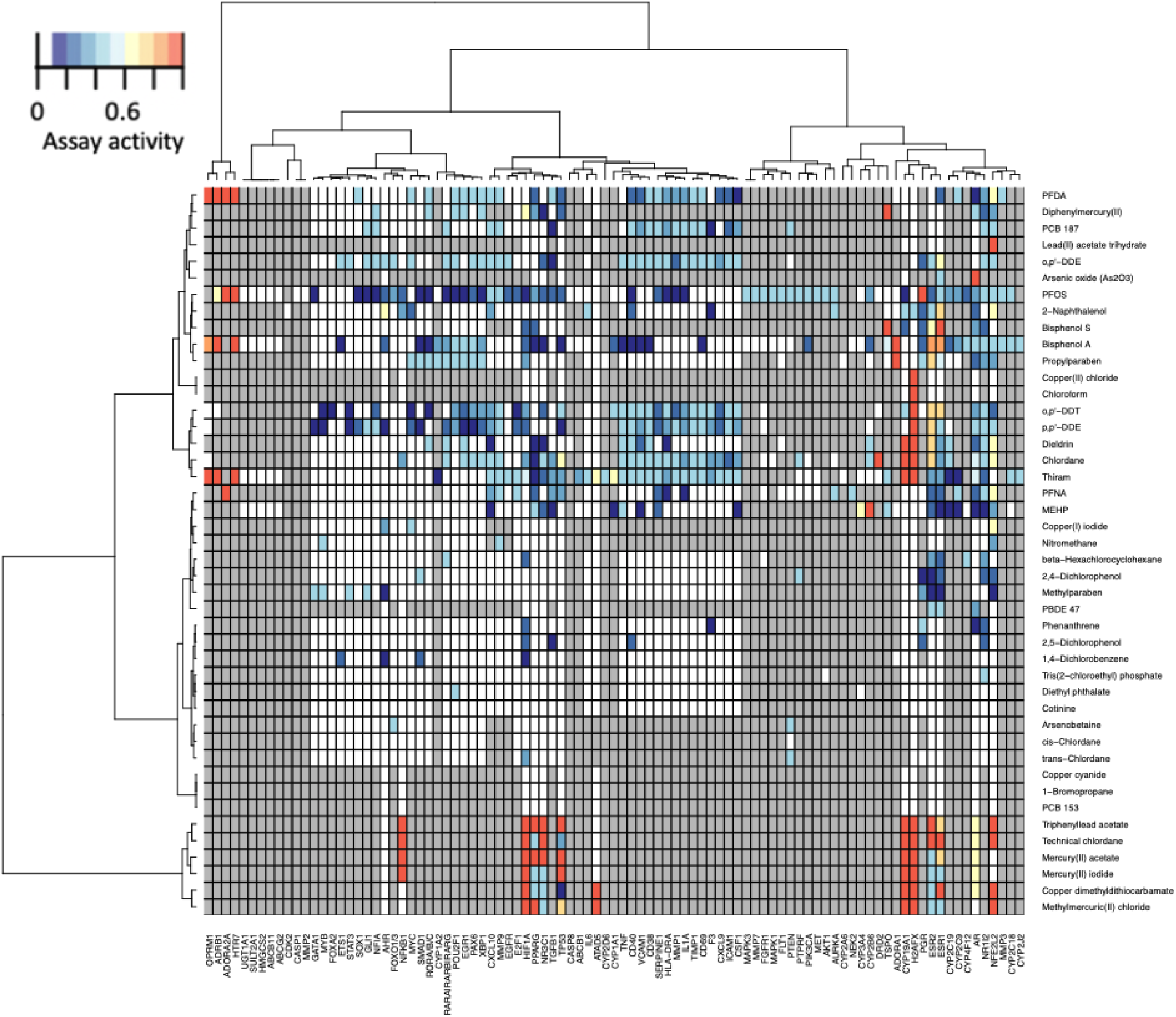
Biological Activity of Assays with Breast Cancer Gene Targets. Unbiased hierarchical clustering of selected BC genes and chemicals based on ratio of active (hitcall=1) to total assays run. Breast Cancer genes were identified through a comprehensive literature review. White indicates no assays are run for that chemical with that specific gene target. Gray indicates assays are run but none are active.

### Comparison of biomarker exposure in Black Women with bioactive levels

After characterizing the assay data available from ToxCast relevant to chemicals with disparities, we compare the biologically active chemical concentrations in ToxCast to the biomarker concentrations detected in NHANES (**Figure 3**). We display for each of the 43 selected chemicals, with metals on a separate plot, the observed concentrations of chemical biomarkers in the NHANES population (blue), the distribution of ACCs for active ToxCast assays (red) and the distribution of concentrations across active ToxCast assays assessing BC gene targets (green) (**Figure 4**). Overlap indicates that the concentration of the relevant chemical biomarkers measured in non-Hispanic Black women in NHANES occurs within the assay ACCs for the same chemical. Overlapping non-metal chemicals include: Methylparaben, p,p’-DDE, Propylparaben, 2,5-Dichlorophenol, 2-Naphthalenol, Dieldrin, beta-Hexachlorobiphenyl, Diethyl phthalate, 2,4-Dichlorophenol, MEHP, thiram, phenanthrene, 1-bromopropane, 1,4-dichlorobenzene, and cotinine **(Figure 4A).** Overlapping metals include lead, arsenic, copper and Arsenobetaine **(Figure 4B)**. **Supplementary Table 5** quantifies the overlap in concentrations of our three studied datasets: NHANES, ToxCast, and ToxCast BC. As indicated in our table, methylparaben has an overlap value of 2.9 log10 uM, this means that the concentration of NHANES, ToxCast, and BC overlapped by 10^2.9 = 794.33 uM.

**Figure 4:**
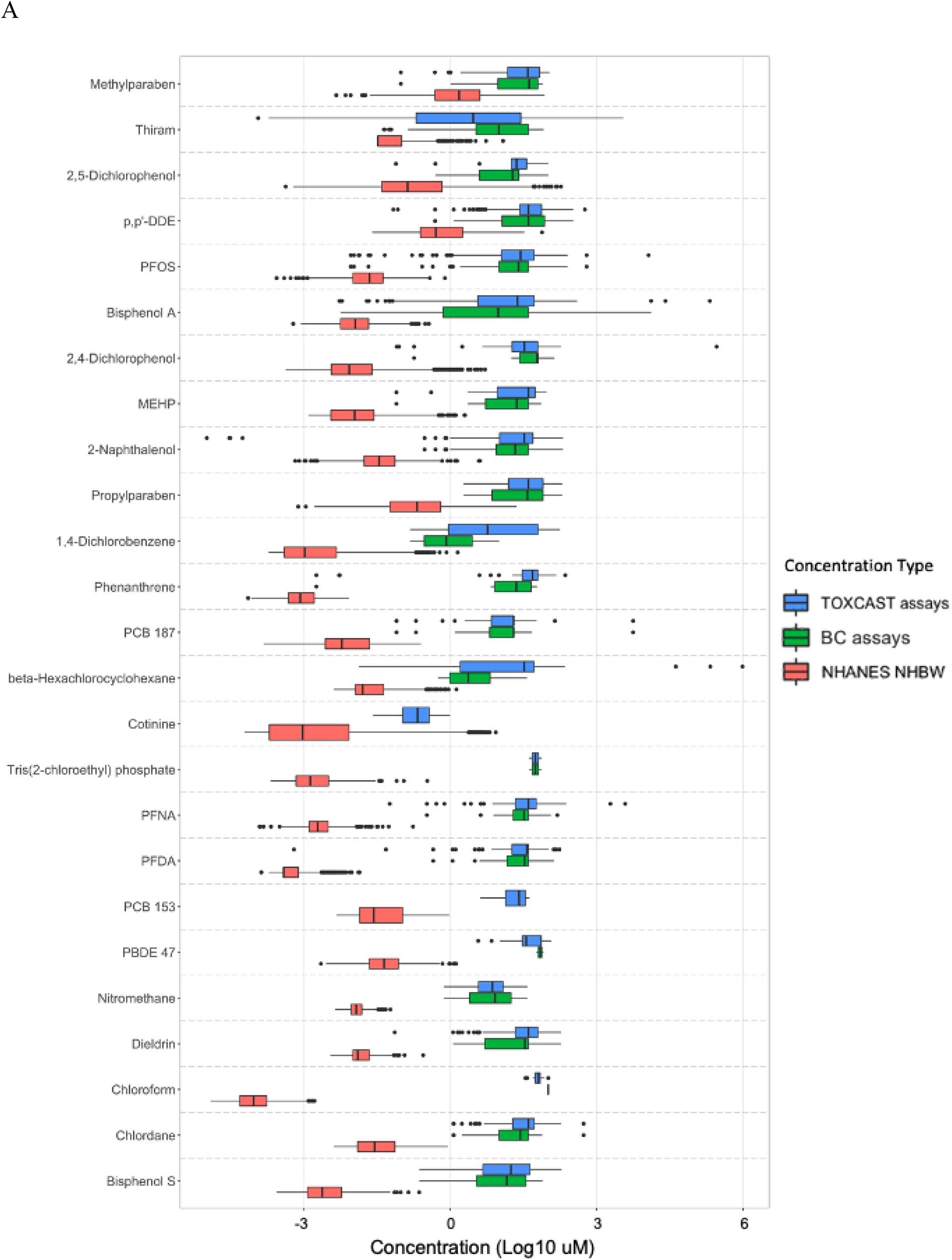
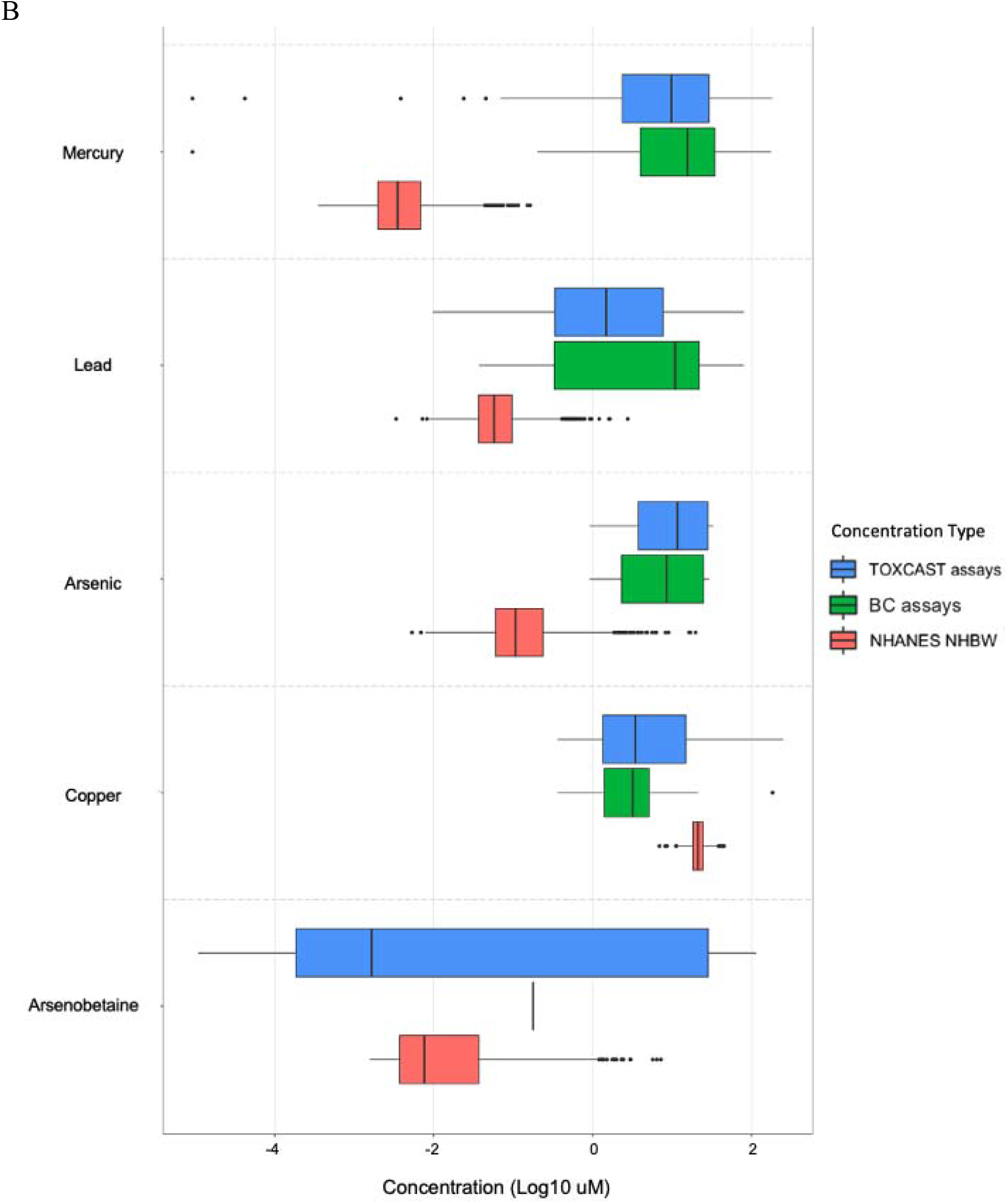
Comparing Chemical Concentrations in ToxCast and NHANES. Log10 ACC concentrations across all active assays from ToxCast (blue) and active BC gene ToxCast assays (green) are compared to the concentrations of the chemical biomarkers measured in NHB women in NAHNES (red). ACC is the concentration at which the dose-response fitted model reaches the cutoff considered “active”. Overlapping distributions indicate that chemical concentrations detectible in NHB women produce a response in ToxCast assays. Chemicals are ordered from highest to lowest overlapping distributions. A) All chemicals excluding metals. B) All metals.

### Breast Cancer Gene Assay Activity and Efficacy for Selected Chemicals

We focused on those chemicals with the most significant overlap between the active assay concentrations and measured biological concentrations in non-Hispanic Black women and the most active BC assays. Graphs in **Figure 5** show the TP and ACC of active assays testing BC genes for methylparaben, 2,5 dichlorophenol, and thiram. For methylparaben, assays with high TP are those testing *ESR2*, *AHR* and *NFE2L2* (**Figure 5A**). In **Figure 4,** the median biomarker concentrations of methylparaben for NHBW is about 1.53 μM. **Figure 5A** indicates that most of the assays are showing a gain or loss in function at this or slightly higher concentrations. The *ESR2* assays for methylparaben measure for activity in the “up” direction, indicating that methylparabens may increase *ESR2* levels. Assays associated with 2,5 dichlorophenol are mostly gain of function for *ESR1*, and those assays with a high TP and ACC are testing for *AR* (**Figure 5B**). In **Figure 4**, the median biomarker concentration of 2,5 dichlorophenol for NHBW is about 0.13 μM, a concentration at which we observe an increase of function in *NR1I2*, *HIF1A*, *and PGR* in **Figure 5B**. Additionally, there is a decrease of function in *TGFB1* and *CD40. TGFB1* and *CD40* are within the range of the median and 75^th^ percentile (0.135 μM and 0.687 μM respectively) observed biomarker concentrations. Assays for thiram with high TP are those testing *ESR1*, *AR*, *HIF1A*, *PPARG*, and *TP53*. This indicated that Thiram is active in many immune assays including *CXCL10*, *CXCL9*, and *IL1A*. These assays indicate a loss in function for these immune genes (**Figure 5C**). According to our concentration box plots, these assays seem to be active at all concentrations observed concentrations in NHBW. Plots for the TP and ACC for the other 40 chemicals are shown in **Supplemental Figure 2**.

**Figure 5:**
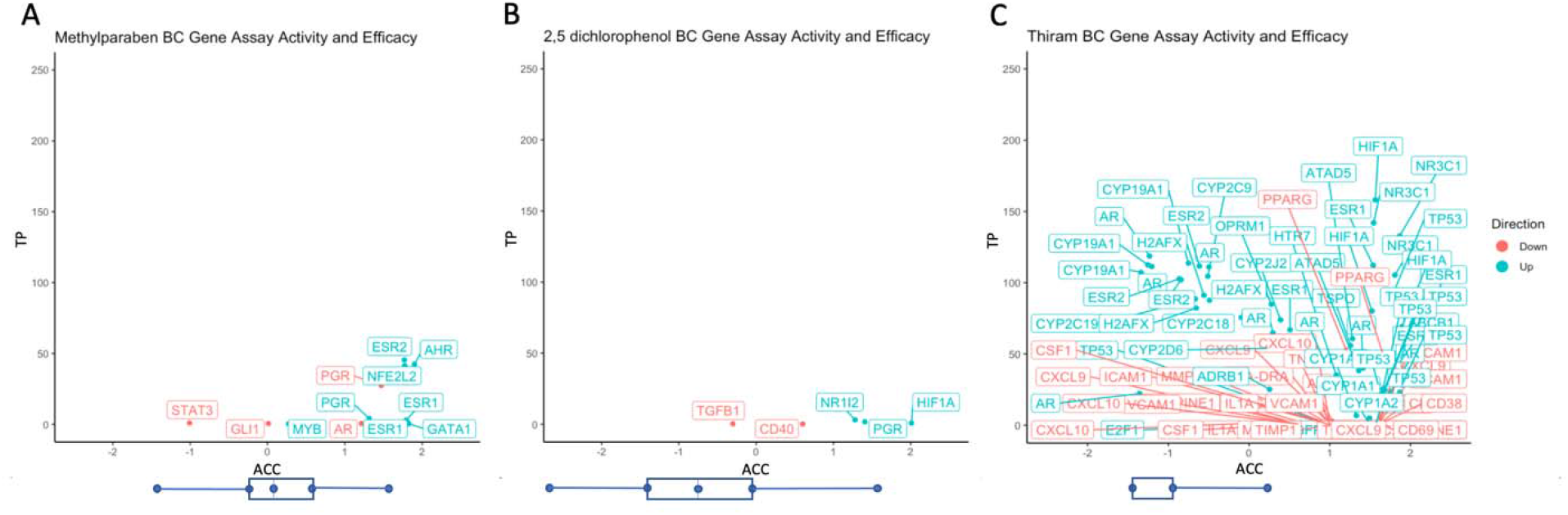
Breast Cancer Gene Assay Activity and Efficacy for Selected Chemicals. These graphs show the TP and ACC of active assays testing BC genes for certain chemicals. Methylparaben (A), 2,5-dichlorophenol (B), and thiram (C) had significant overlap between the active assay concentrations and measured biological concentrations in non-Hispanic Black women and the most active Breast Cancer assays. A higher TP indicates a greater response/efficacy. A higher ACC means a higher concentration at which the dose-response fitted model reaches the cutoff considered “active”. Direction of the assay is reported in red (loss of function) or blue (gain of function). The boxplot underneath the figure indicates the range of concentrations for each chemical in NHANES for NHBW.

## Discussion

Advancing methods to screen candidate chemicals for associations with specific disease outcomes is critical for prioritizing chemicals for further experimental and epidemiological investigation, as well as to design interventions in highly exposed populations. In this study, we aimed to identify chemical exposures which may play an etiologic role in breast cancer racial disparities and prioritize chemicals for further experimental investigation. By performing an analysis which integrates human biomonitoring data from NHANES and biological activity data from the EPA’s ToxCast program, we identified chemicals with significant exposure disparities by race and with biological activity relevant to breast cancer. We found that methylparaben, propylparaben, 2,5-dichorophenol, DDT and DDT metabolites (p,p’-DDE), and thiram are chemicals of particular interest for further experimental investigation based on our observations of breast cancer associated biological activity at concentrations relevant to human exposures.

In this study, we used lipid adjusted concentrations measured in serum lipids to best model a lipophilic chemicals’ concentrations in adipose rich tissue like the breast. We believe this measurement would be the most accurate. However, we do acknowledge that further enhancements to our method could be made as physiologically based toxicokinetic models of toxicant distribution to the human mammary gland are developed for these lipophilic chemicals.

The prioritization method of determining which assays are run for specific chemicals in ToxCast is based on a hazard prediction, where chemicals with low assay activity are categorized as “low priority” and not further tested (Dix *et al*., 2007). Chemicals with intermediate assay activity undergo further testing to re-evaluate prioritization. Finally, chemicals with high assay activity are prioritized as hazardous and recommended for further screening and testing outside of ToxCast (Dix et al. 2007). The high activity of assays that are related to breast cancer genes at relevant concentrations to non-Hispanic Black women provides evidence that these chemicals may be associated with breast cancer disparities. A possible limitation of ToxCast prioritization method is the limited data collected on some of the most harmful chemicals. Due to the high ratio of activity for a dangerous chemical, such as metals lead and mercury, they are classified as hazardous and removed from rotation through the assays to undergo further testing outside of ToxCast. This then leads to a limited amount of data for the hazardous chemicals, making it difficult to study the chemicals mechanism of action using this methodology.

Several other studies have used ToxCast for chemical prioritization including environmental chemicals for obesity outcomes, evaluation of food-relevant chemicals, assessment of human indoor exposome of chemicals in dust, and the evaluation of novel flame retardants (Auerbach S. *et al*., 2016; Karmaus *et al*., 2016; Dong *et al*., 2019; Bajard *et al*., 2019). Most of these studies maintained a similar methodology to our current study, where assay information from ToxCast is analyzed for a predefined set of chemicals. However, the study conducted by (Auerbach S. *et al*., 2016) used a different approach where experts were used to determine assays testing for biological processes that play important roles in the mechanism of diabetes and obesity. After selecting assay targets relevant in the biological process for diabetes and obesity, they were able to identify chemicals which perturbed these targets and pathways. Differing approaches to prioritize chemicals and assess biological activity and relevant doses demonstrates the broad utility of ToxCast.

TNBCs and aggressive breast cancers with poor prognosis have been shown to have a stem cell-like biological phenotype (Thong *et al*., 2020; Malta *et al*., 2018), and developmental pathways are commonly dysregulated by toxicant exposures (Thong *et al*., 2019). African American women are disproportionately likely to be diagnosed with TNBC and are more likely to die of breast cancer across all subtypes. Here, we also found that stemness related genes, including *AHR*, *SOX1*, *GLI1*, and *HIF-1A*, are targets of chemicals with characterized exposure disparities in African American women. *AHR* has been shown to regulate immunity and maintain cell differentiation (Hao and Whitelaw, 2013; Kawajiri and Fujii-Kuriyama, 2017), deregulation of this protein may contribute to tumor initiation (Gasiewicz *et al*., 2008). Over expression of HIF1A has been shown to promote tumor growth in breast cancer (Schwab *et al*., 2012). *HIF-1A* activation in cancer cells regulates expression of stemness transcription factors, including OCT4, NANOG, SOX2, and KLF4 (Mathieu *et al*., 2011). A recent study found that chemotherapy induced *HIF-1A* controlled pathway led to epigenetic activation of pluripotency factor transcription (Lu *et al*., 2020), suggesting that this pathway can be activated by stressor exposures. We previously showed that *HIF-1A* is required for stem cell proliferation from normal human mammary gland *in vitro*, including both spheroid formation and organoid growth (Rocco *et al*., 2018). Determining which chemicals can push cells into a stemness phenotype can help in testing prioritization as well as identify novel mechanisms by which chemical exposures can promote the development of aggressive breast cancers.

Through an unbiased approach, we observed activation patterns in multiple breast cancer relevant biological pathways for different chemicals with characterized exposure disparities. A significant advantage of ToxCast data is the wide range of biological targets assessed for each chemical. For example, immune system-associated genes, including *IL1A*, *CXCL10*, *VCAM1*, and *STAT3* are commonly targeted by many of these chemicals, including chlordane, PFOS, DDT, PFNA, and PFDA. The immune system plays an important role in cancer development and prognosis through numerous mechanisms, such as cancer associated inflammation and immune tolerance. Studies have correlated tumors with high immune cell infiltration with more aggressive subtypes of breast cancer such as basal-like or TNBC (O’Meara *et al*., 2019; Acerbi *et al*., 2015). Estrogen receptor (*ESR1*) and androgen receptor (*AR*) are also common targets of chemicals with exposure disparities. For example, p,p’-DDE, propylparaben, and thiram exposure all increased AR and ESR1 function, although at very different doses, with thiram activating these receptors at a much lower dose (**Figure 5**). AR expression has been shown to stimulate cellular proliferation through MYC regulation and induce invasiveness through PI3-K signaling in TNBC development. Additionally, studies indicate that AR may be a key factor in chemoresistance (Giovannelli *et al*., 2018, 2019). Nearly 70% of breast tumors are shown to express ER, where high expression results in increased proliferation and increased expression of phosphorylated AKT (Gonzalez *et al*., 2019).

We encountered methodological challenges in performing this integrated NHANES and ToxCast analysis which provides our study with several limitations. First, we are limited by the assays run per chemical, which varied widely. Since there are several chemicals that did not have assays run for specific gene targets, deriving pathway analysis results from these data using Gene Ontology or Gene Set Enrichment Analysis was not successful. Additionally, we recognize that multiple chemicals may lead to the same metabolite. In our data, this would result in an overrepresentation of that metabolite. Second, due to the possibility of false positive and negative hit calls, Toxcast added a processing step to assign warnings known as “flags.” Flags are recommended to be considered with discretion due to how ToxCast’s flag automation process can lead to errors in an assay’s flag assignment (Judson *et al*., 2016 and EPA, 2018). Hence, we did not exclude data based on flags. Third, scientists using ToxCast data have identified a phenomenon known as cytotoxic burst (CTB) where many of the assays begin to respond at cytotoxic concentrations of the chemicals (Fay *et al*., 2018b). The bounds of the CTB were determined to identify those assays that elicit cytotoxic responses at concentrations lower than the CTB, therefore prioritizing those assays with high specificity but low responsivity. In the current study, we did not take CTB into account, although we did determine those chemicals with responsive assays within the biological concentrations observed in NHANES (**Figure 3**).

Another limitation of our study is that we are using concentrations of chemicals in NHANES participants using urine and blood measures, rather than concentrations of chemicals measured directly in breast tissue. Persistent lipophilic chemicals are preferentially stored in the mother’s breast adipose tissue (Mead, 2008). The female body can mobilize lifetime fat stores to produce milk for her infant, therefore transmitting the persistent environmental contaminants to her newborn during breastfeeding (Lehmann *et al*., 2014). Chemicals such as BPA, methylparaben (MP), and propylparaben (PP), dichloro-diphenyl-trichloroethane (DDT) and metabolite dichlorodiphenyldichloroethylene (DDE) have all been detected in breast milk with concentrations ranging from 0.0005μM – 0.0033μM (PP), 0.0033μM – 0.015 μM (MP), 0.0013μM – 0.0455μM (BPA), 0.069μM – 50.31μM (DDE) to 0.0031μM – 19.46μM (DDT). (Hines *et al*., 2015; Mendonca *et al*., 2014; Agency for Toxic Substances and Disease Registry, 2008). The development of physiologically based toxicokinetic models designed to predict breast tissue concentrations of chemicals based on biomarkers commonly assessed in blood and urine would significantly improve our ability to estimate the impacts of these exposures on breast biology in diverse human populations.

In conclusion, our results demonstrate that African American women are exposed to chemicals with breast cancer associated biological activity at doses relevant to human exposure, for multiple chemicals for which they have higher biomarker levels than white women. Future studies should aim to analyze pathways and genes identified as active at biologically relevant concentrations as more ToxCast assay data becomes available. Additionally, ongoing work in our laboratory is using *in vitro* assessment of chemicals in primary human mammary cells collected from non-Hispanic Black and non-Hispanic White women to validate the biological activity which we identified here. These experiments will help to inform whether integration of exposure data from NHANES with biological activity data from Toxcast is a relevant methodology to identify hazardous chemicals that may be involved in the development and prognosis of breast cancer.

## Notes

### Competing Interest Statement

The authors have declared no competing interest.

