## Supplemental Figure 1 for "Identifying the Link Between Chemical Exposures and Breast Cancer in African American Women via ToxCast High Throughput Screening Data"

### Slide 1
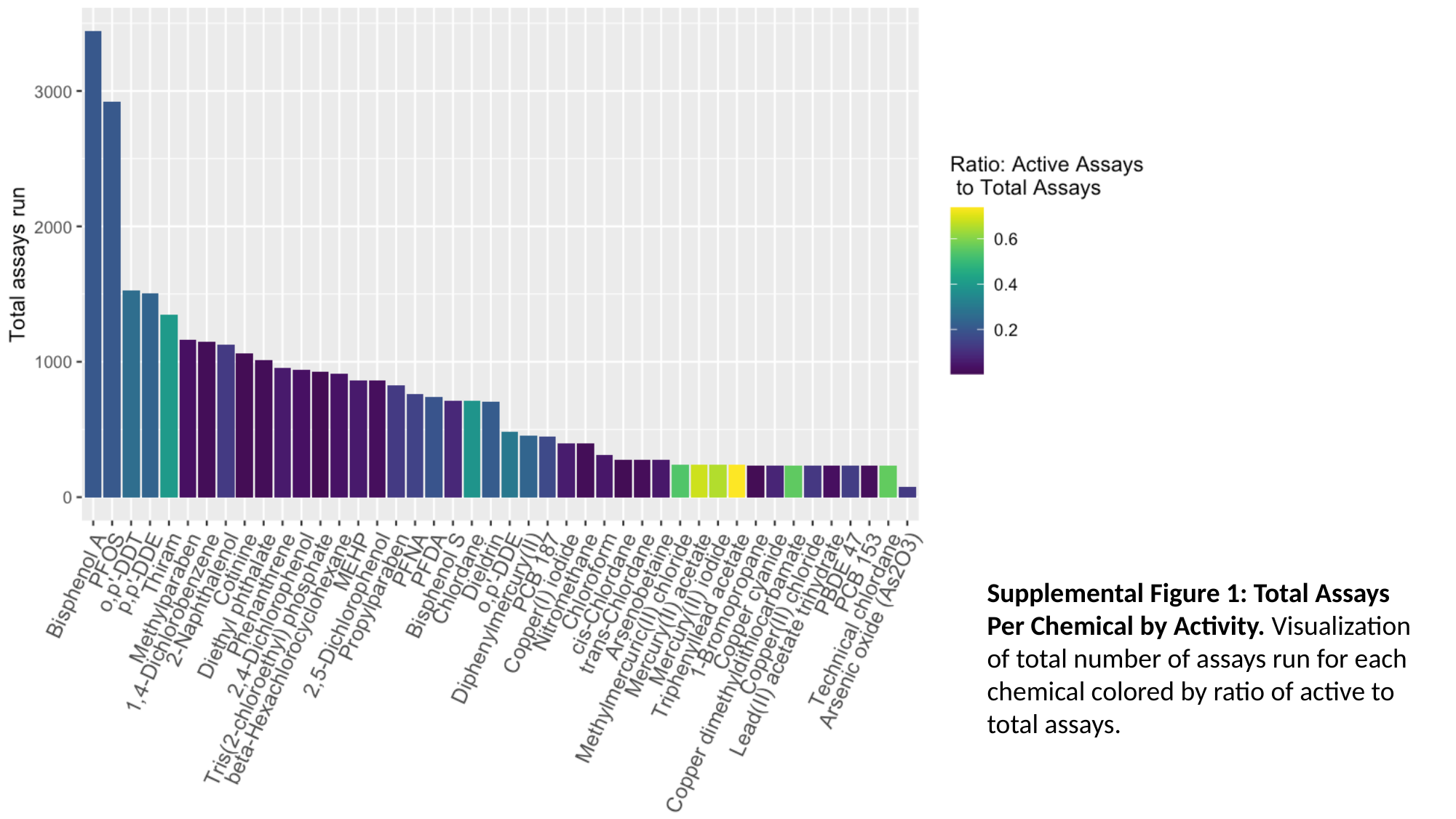

Supplemental Figure 1: Total Assays Per Chemical by Activity. Visualization of total number of assays run for each chemical colored by ratio of active to total assays.
