## Supplemental Figure 2 for "Identifying the Link Between Chemical Exposures and Breast Cancer in African American Women via ToxCast High Throughput Screening Data"

### Slide 1
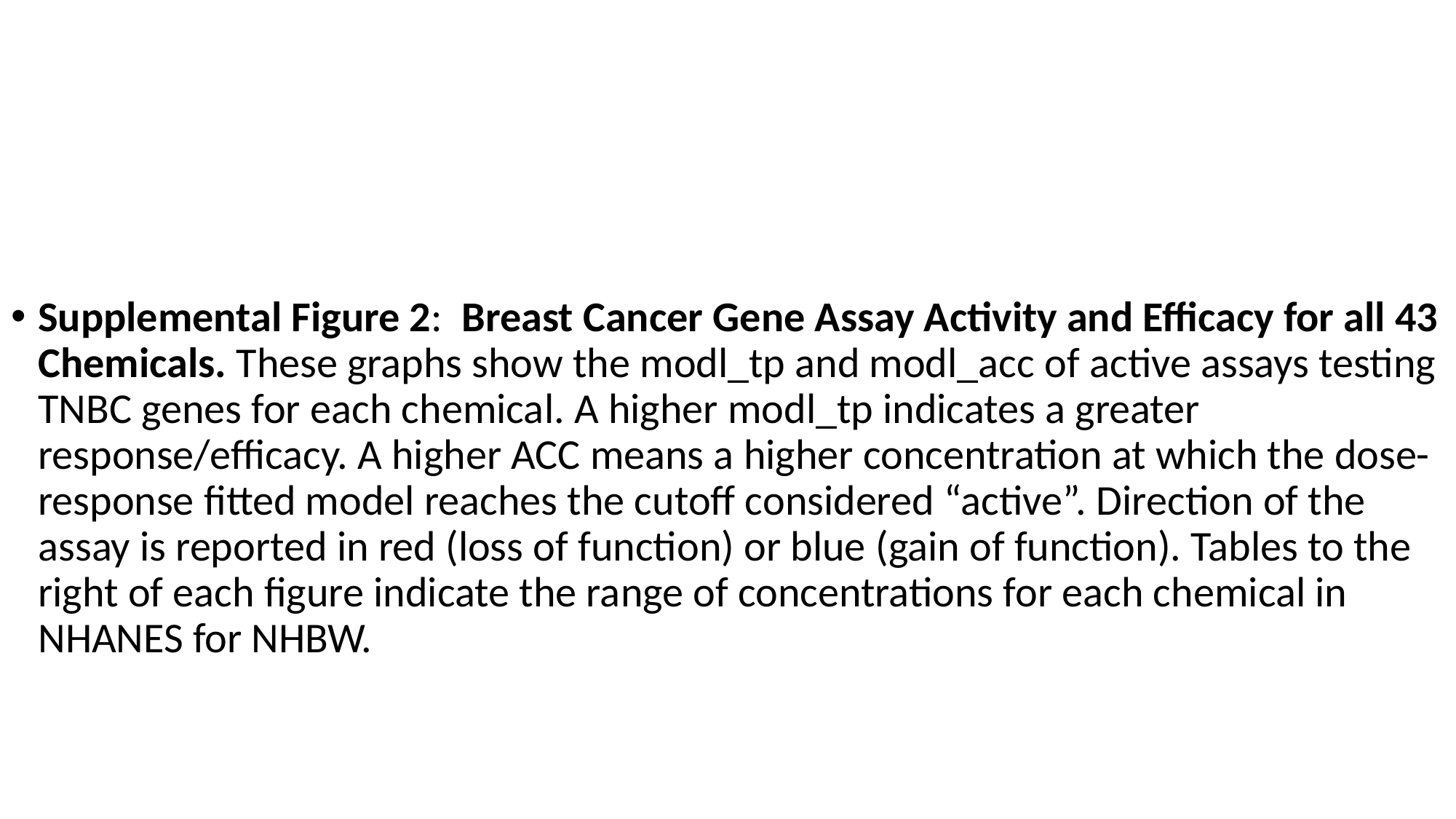

Supplemental Figure 2: Breast Cancer Gene Assay Activity and Efficacy for all 43 Chemicals. These graphs show the modl_tp and modl_acc of active assays testing TNBC genes for each chemical. A higher modl_tp indicates a greater response/efficacy. A higher ACC means a higher concentration at which the dose-response fitted model reaches the cutoff considered “active”. Direction of the assay is reported in red (loss of function) or blue (gain of function). Tables to the right of each figure indicate the range of concentrations for each chemical in NHANES for NHBW.

### Slide 2
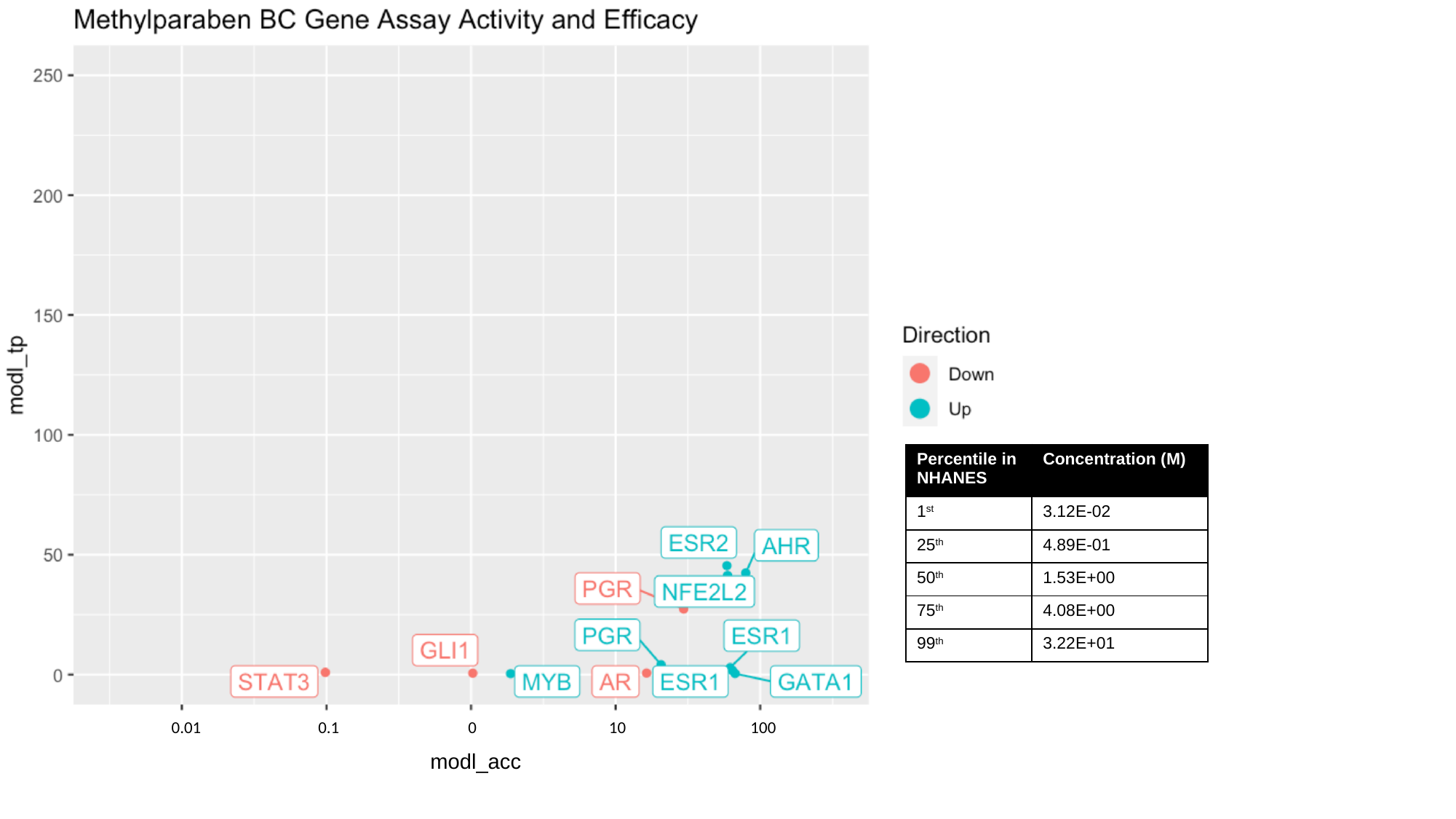

100
0.01
0.1
0
10
modl_acc

### Slide 3
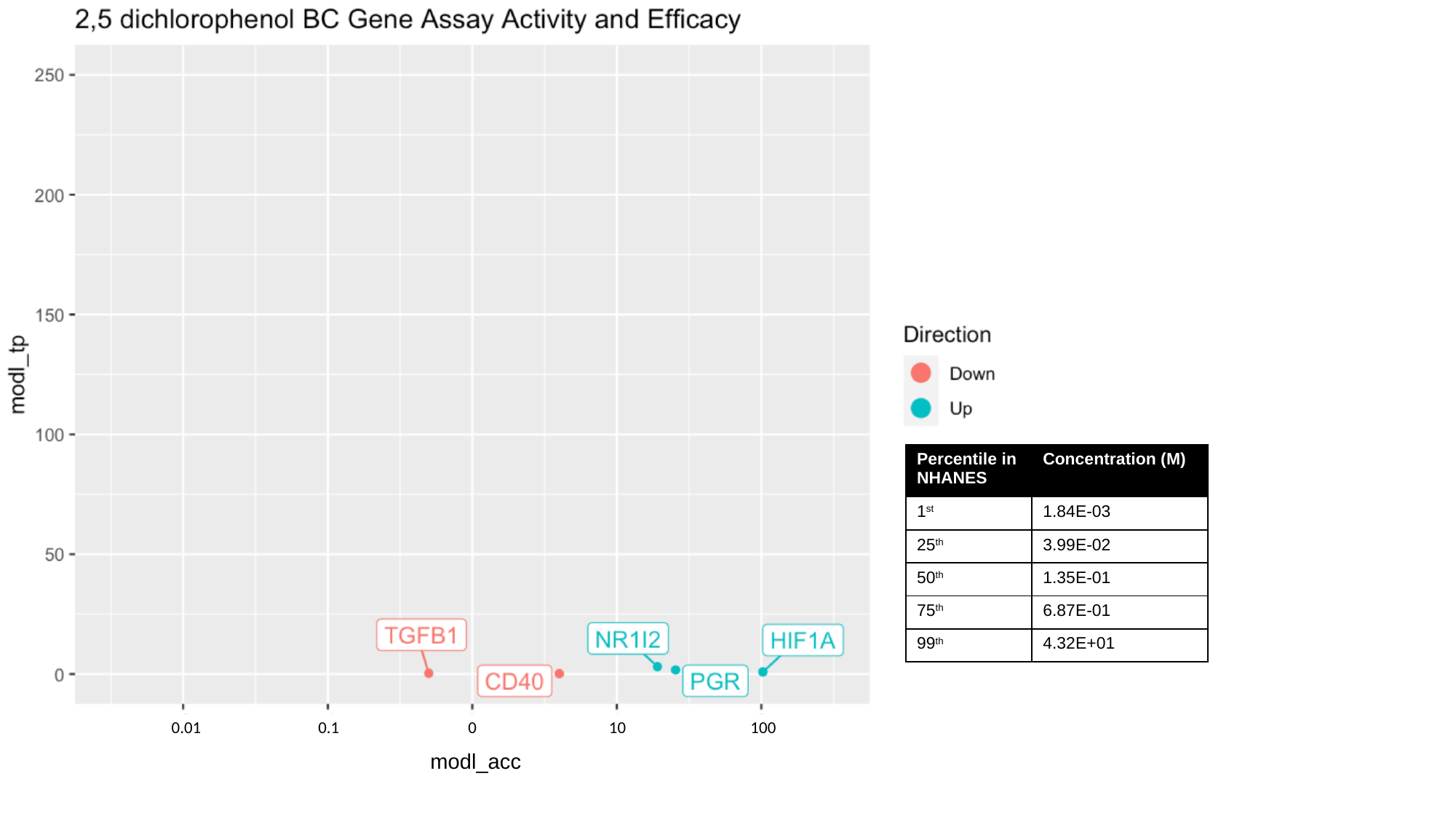

100
0.01
0.1
0
10
modl_acc

### Slide 4
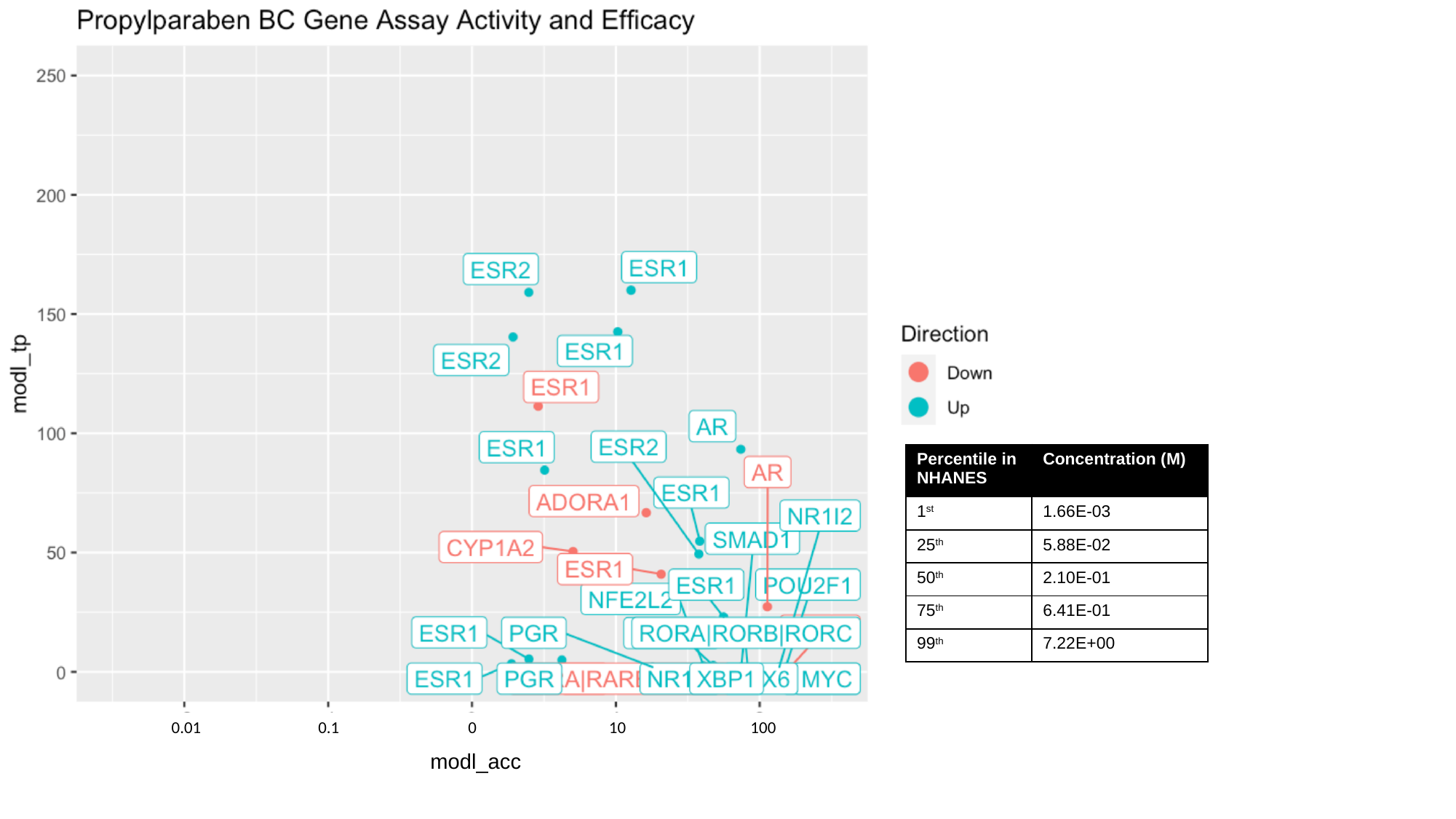

100
0.01
0.1
0
10
modl_acc

### Slide 5
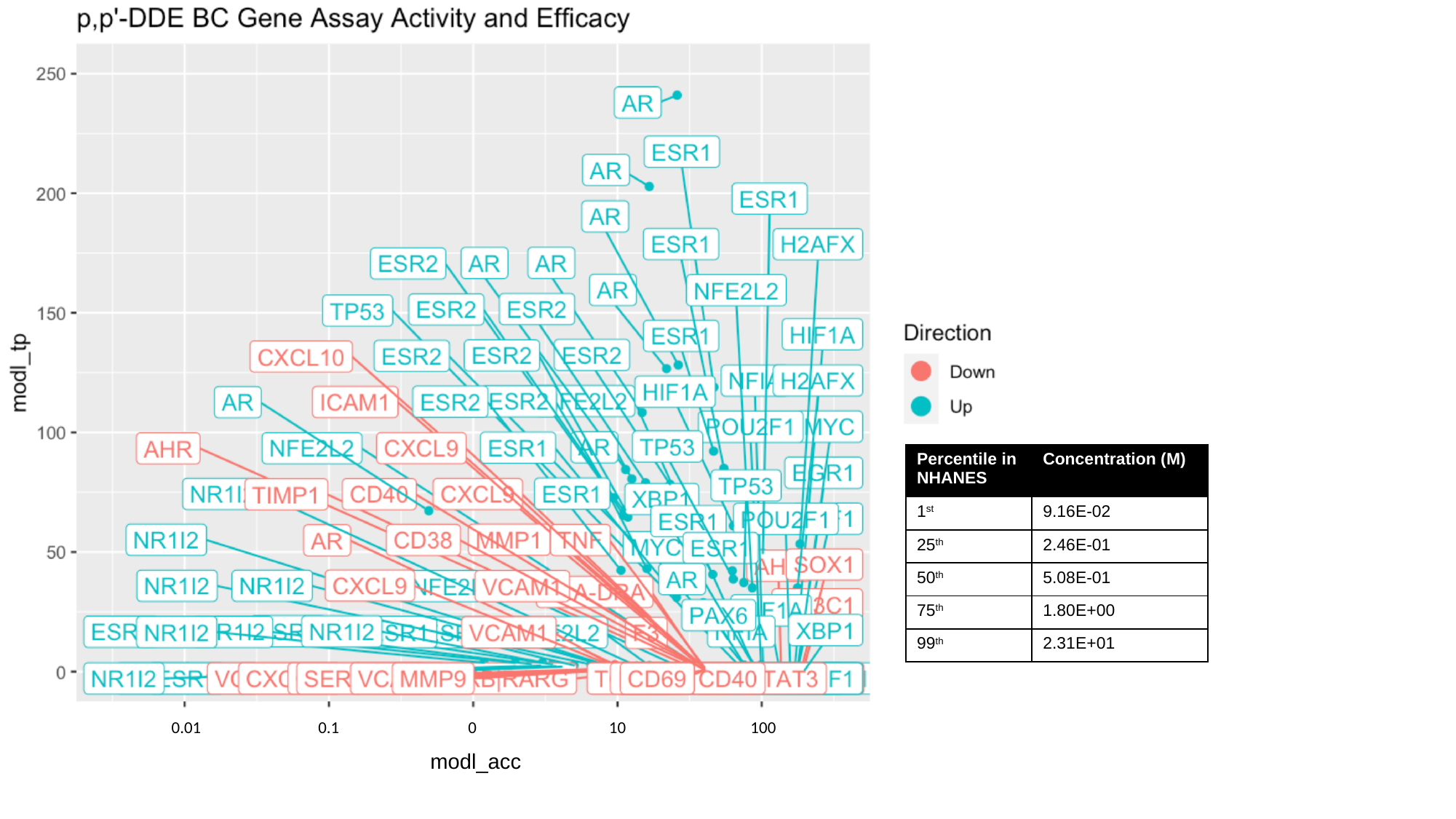

100
0.01
0.1
0
10
modl_acc

### Slide 6
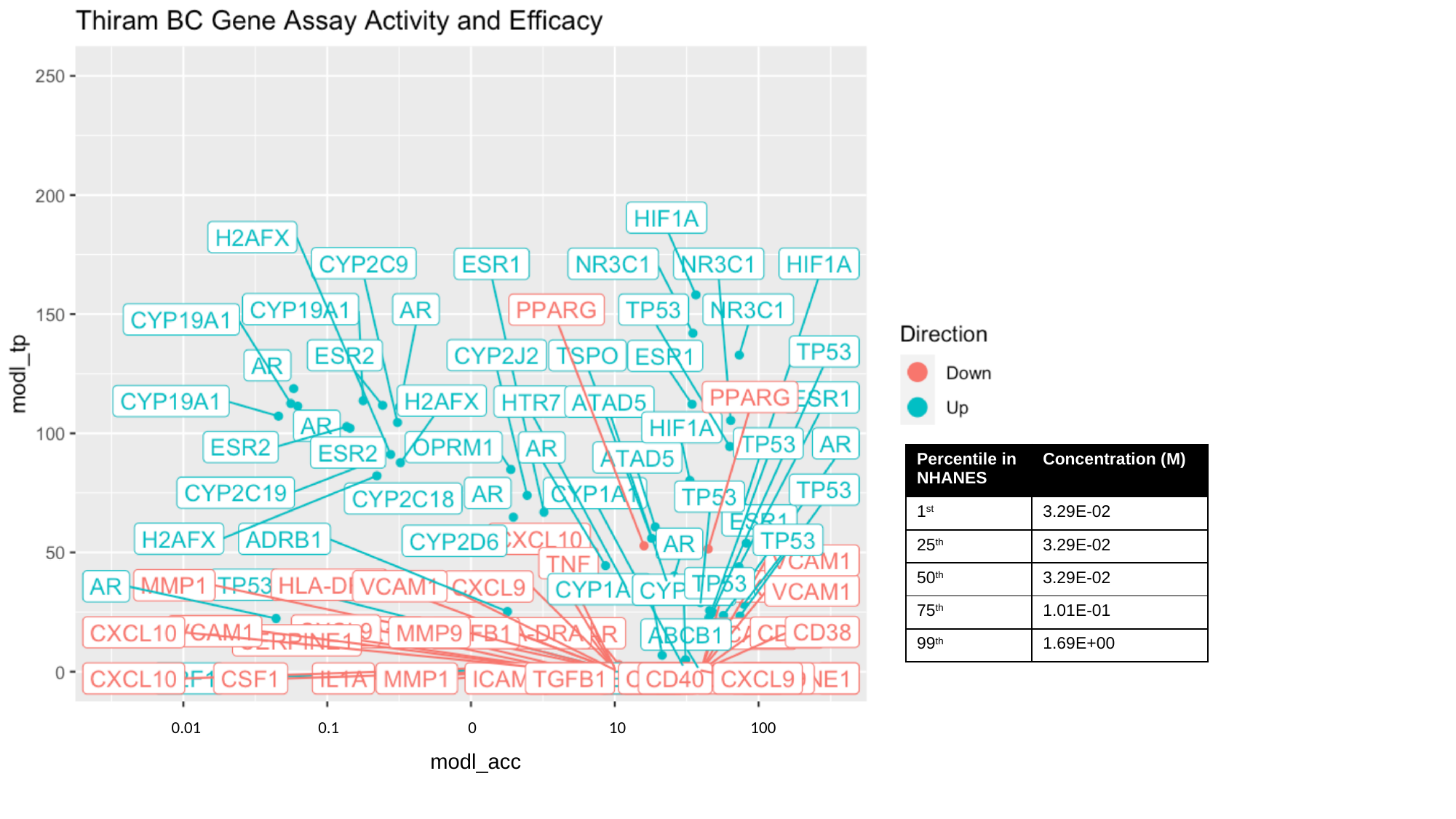

100
0.01
0.1
0
10
modl_acc

### Slide 7
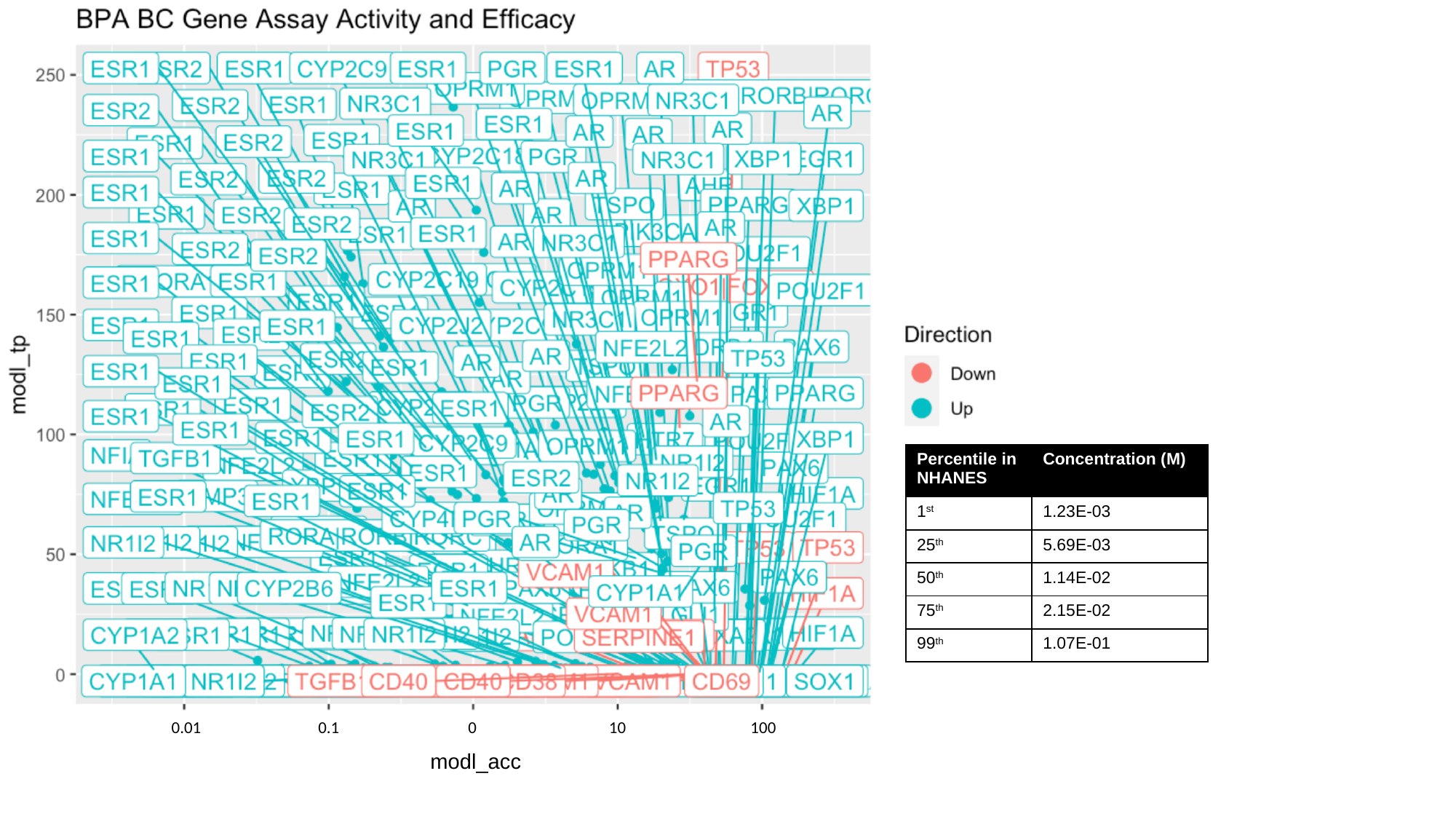

100
0.01
0.1
0
10
modl_acc

### Slide 8
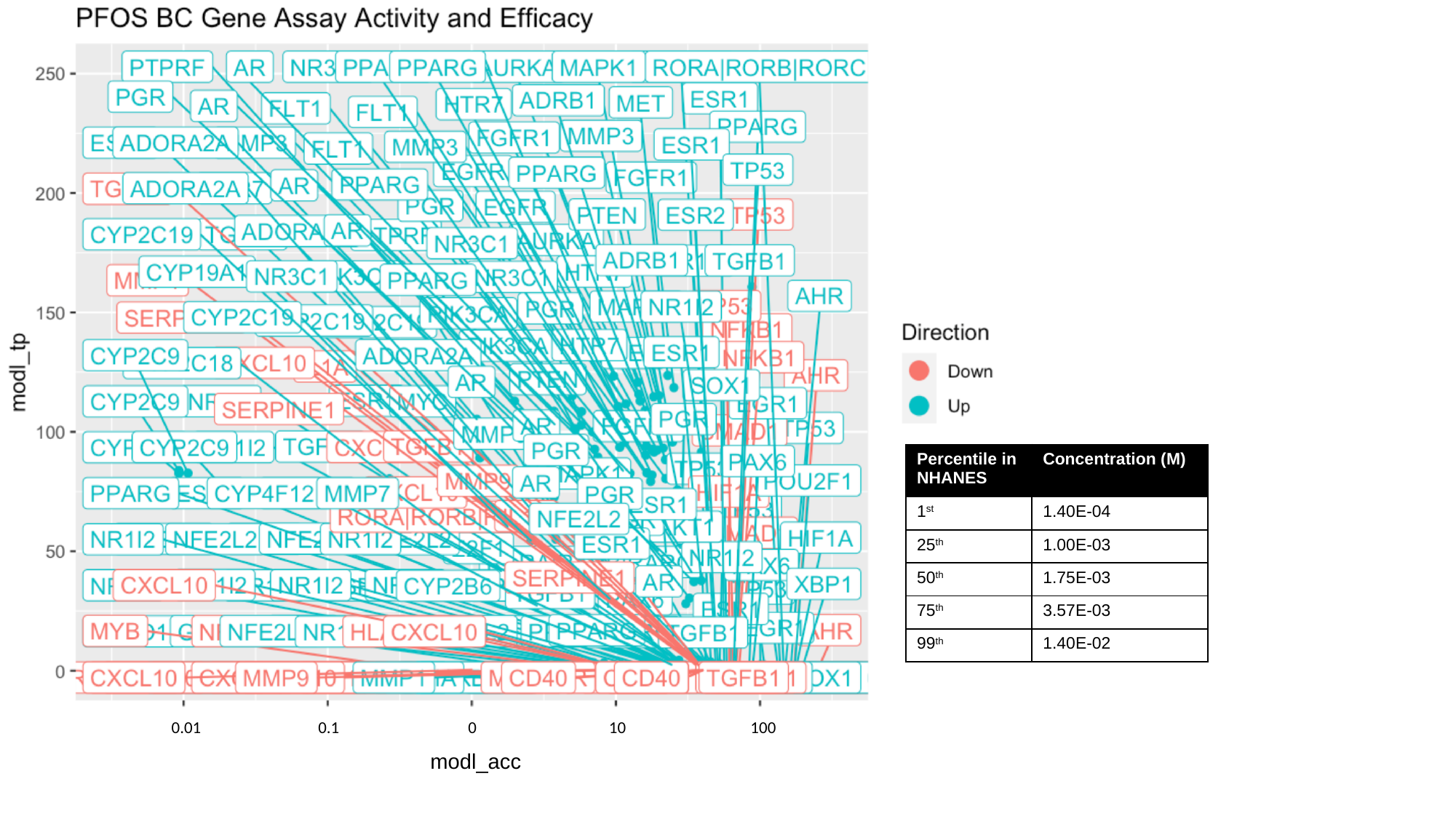

100
0.01
0.1
0
10
modl_acc

### Slide 9
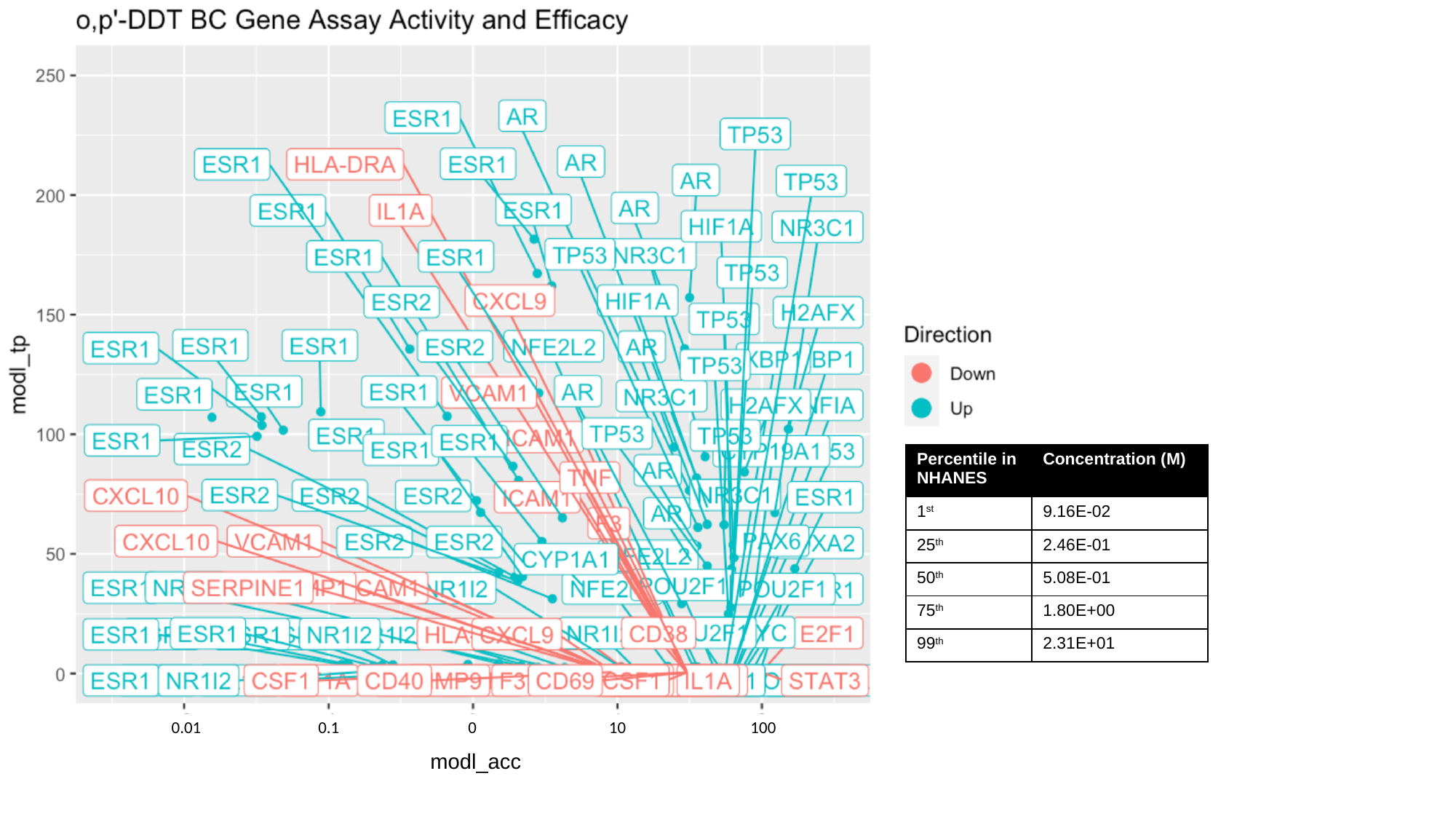

100
0.01
0.1
0
10
modl_acc

### Slide 10
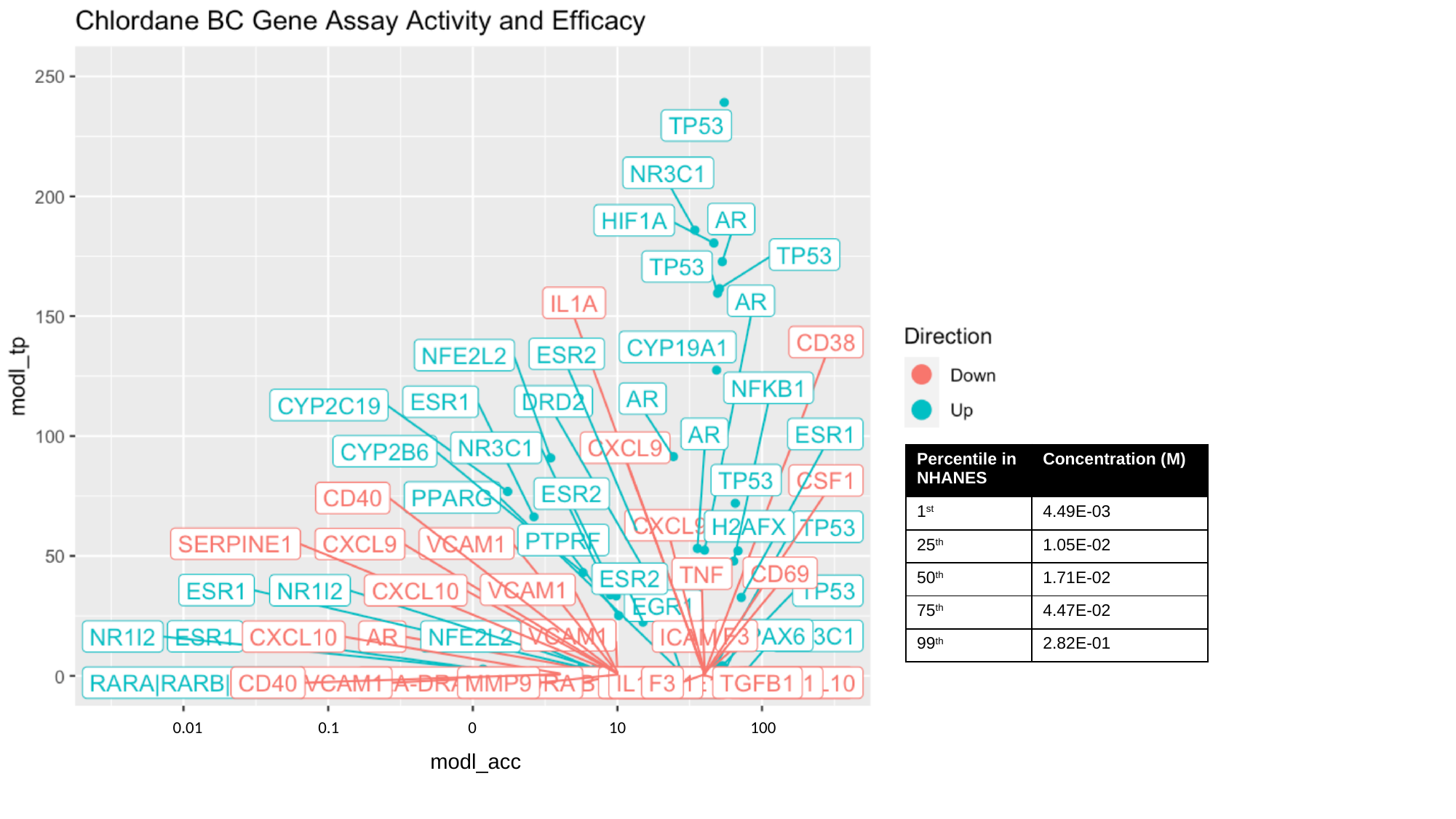

100
0.01
0.1
0
10
modl_acc

### Slide 11
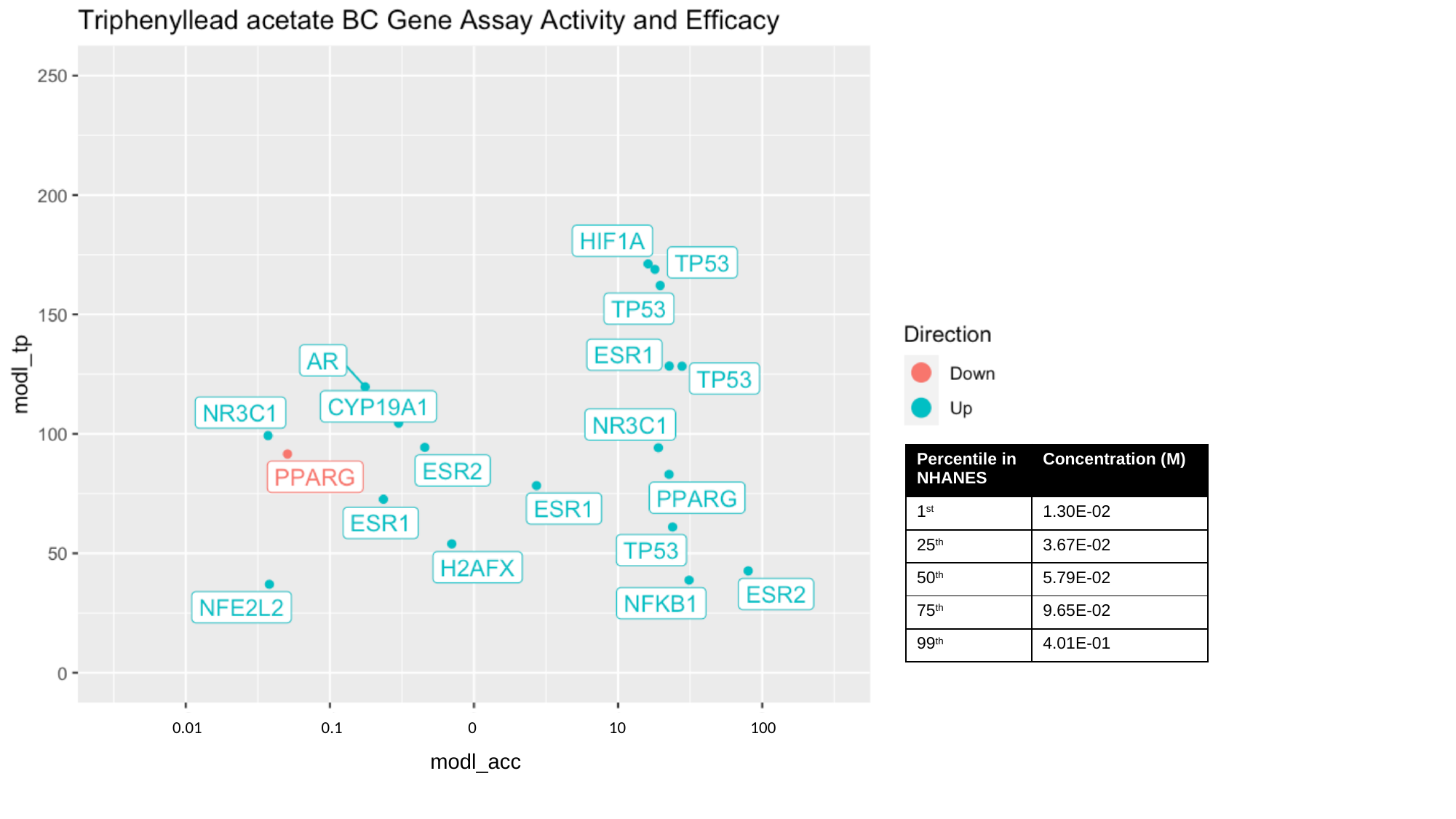

0.01
100
0.1
0
10
modl_acc

### Slide 12
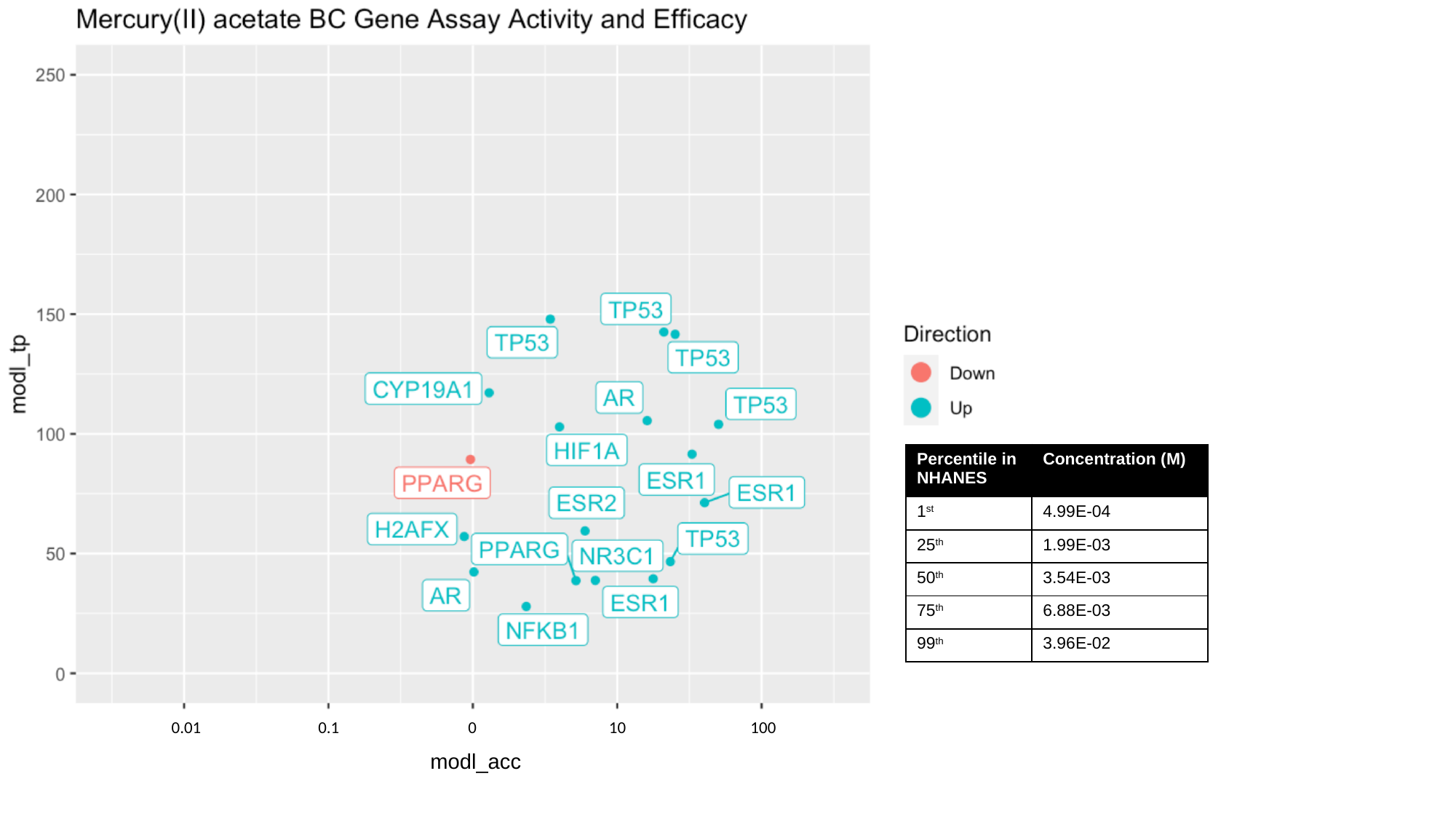

100
0.01
0.1
0
10
modl_acc

### Slide 13
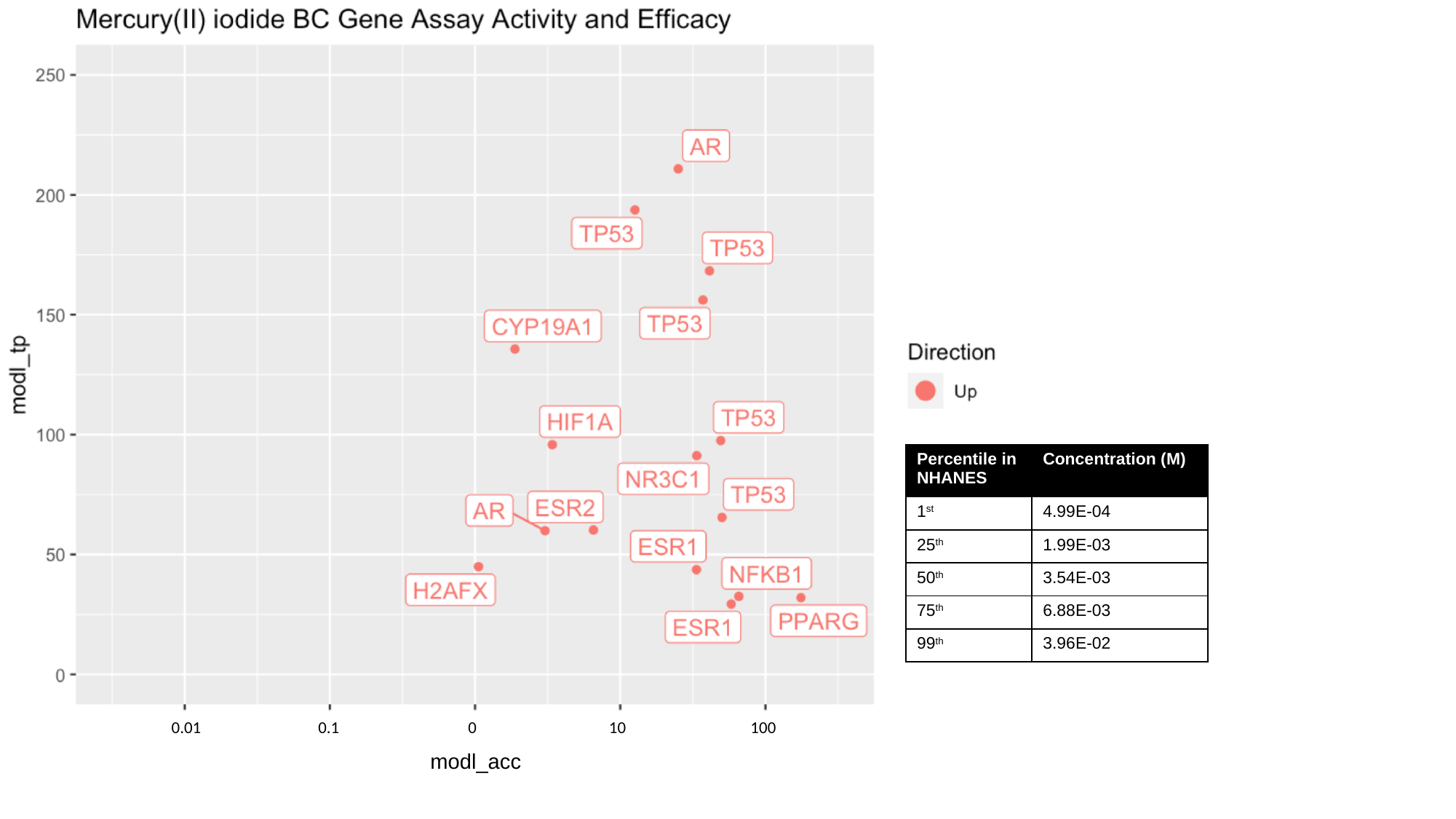

100
0.01
0.1
0
10
modl_acc

### Slide 14
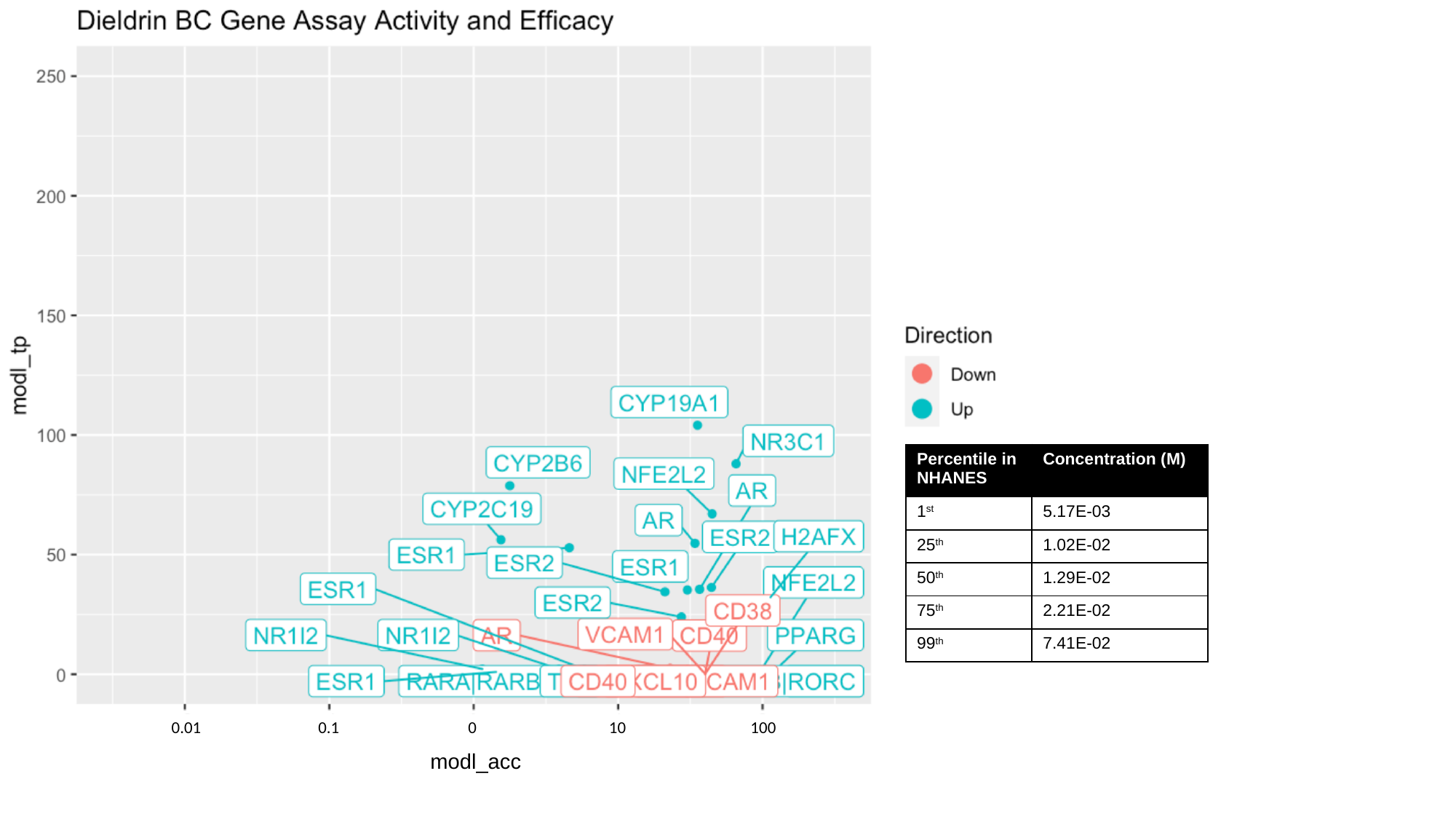

100
0.01
0.1
0
10
modl_acc

### Slide 15
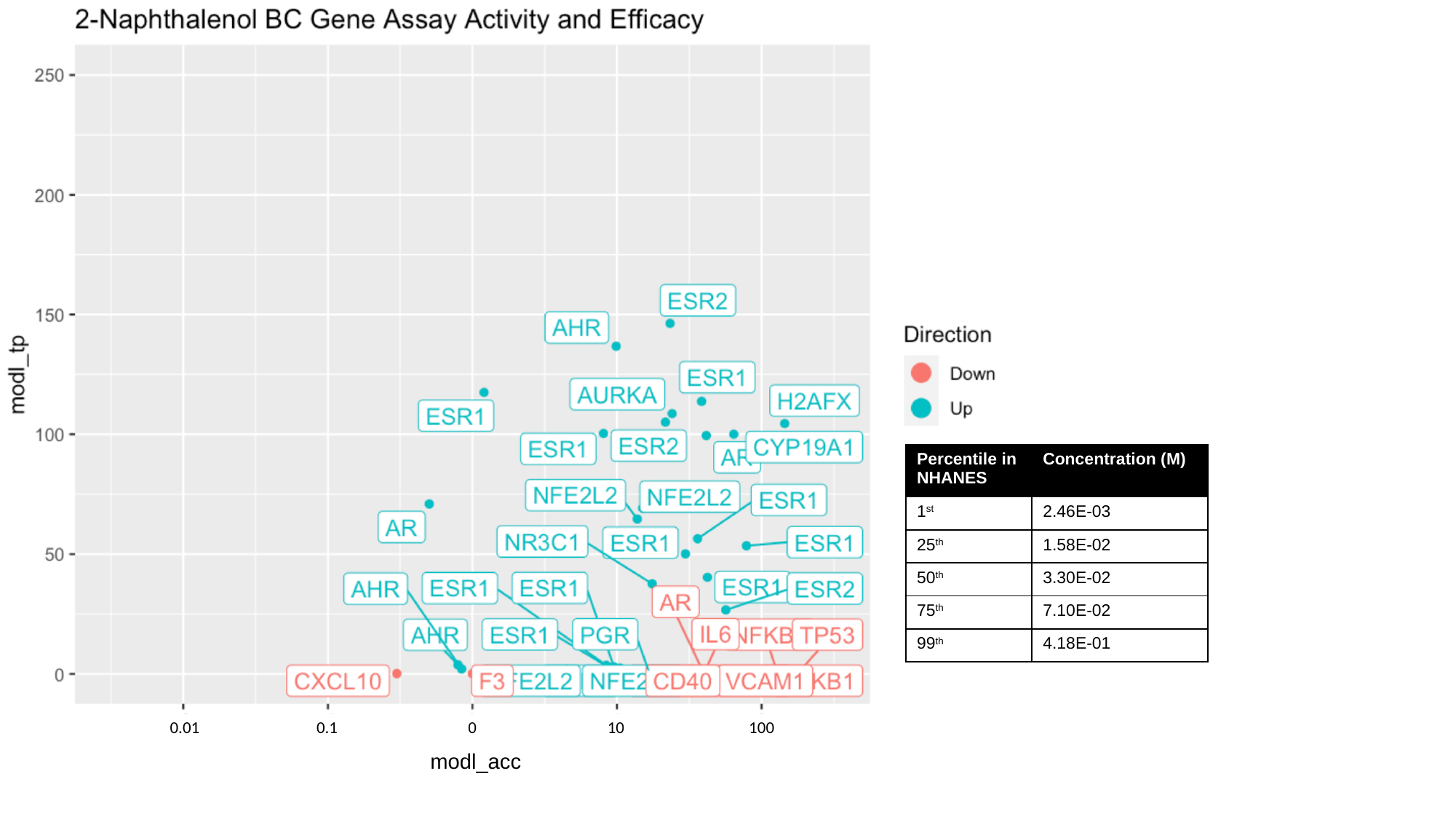

100
0.01
0.1
0
10
modl_acc

### Slide 16
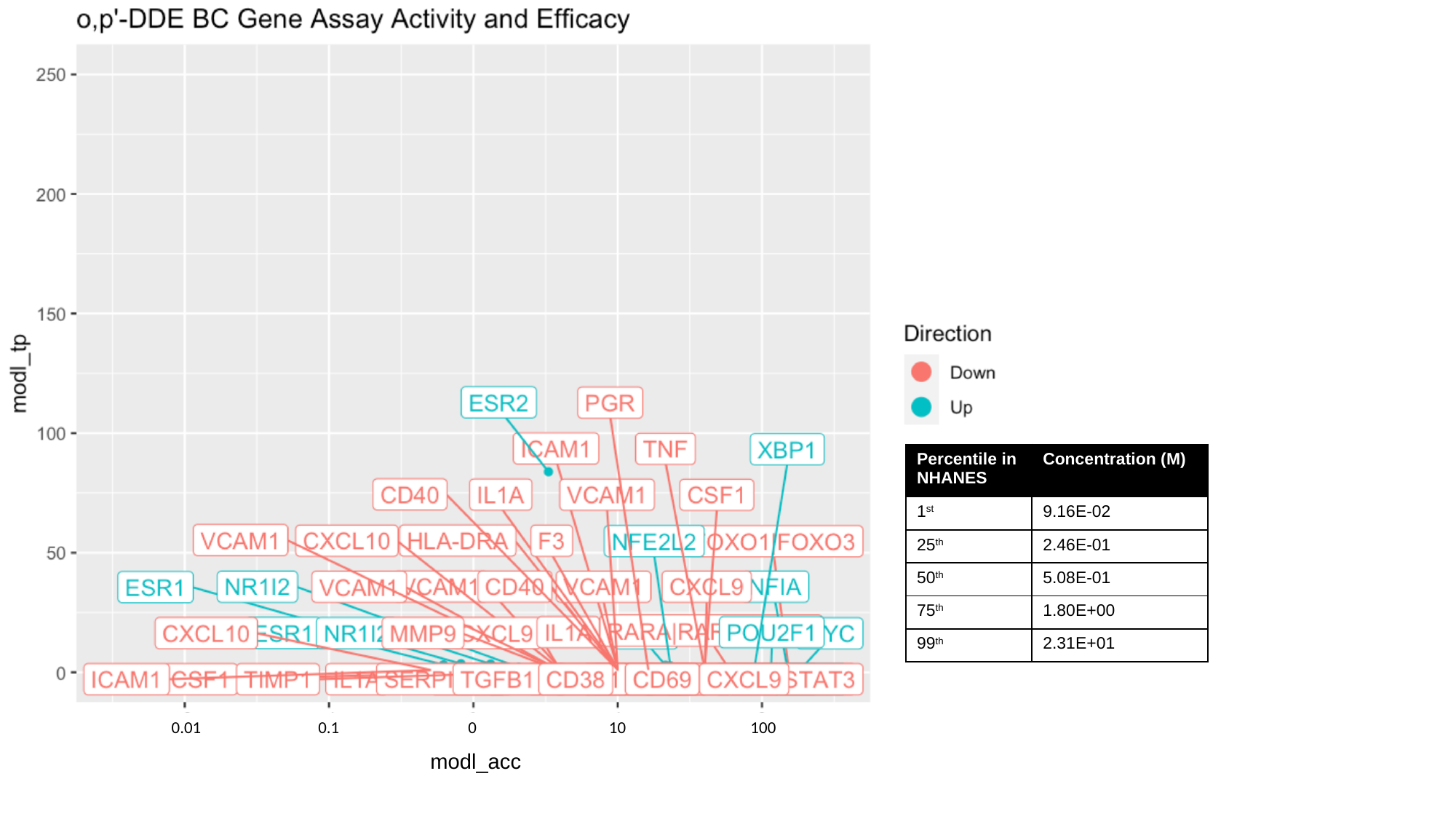

100
0.01
0.1
0
10
modl_acc

### Slide 17
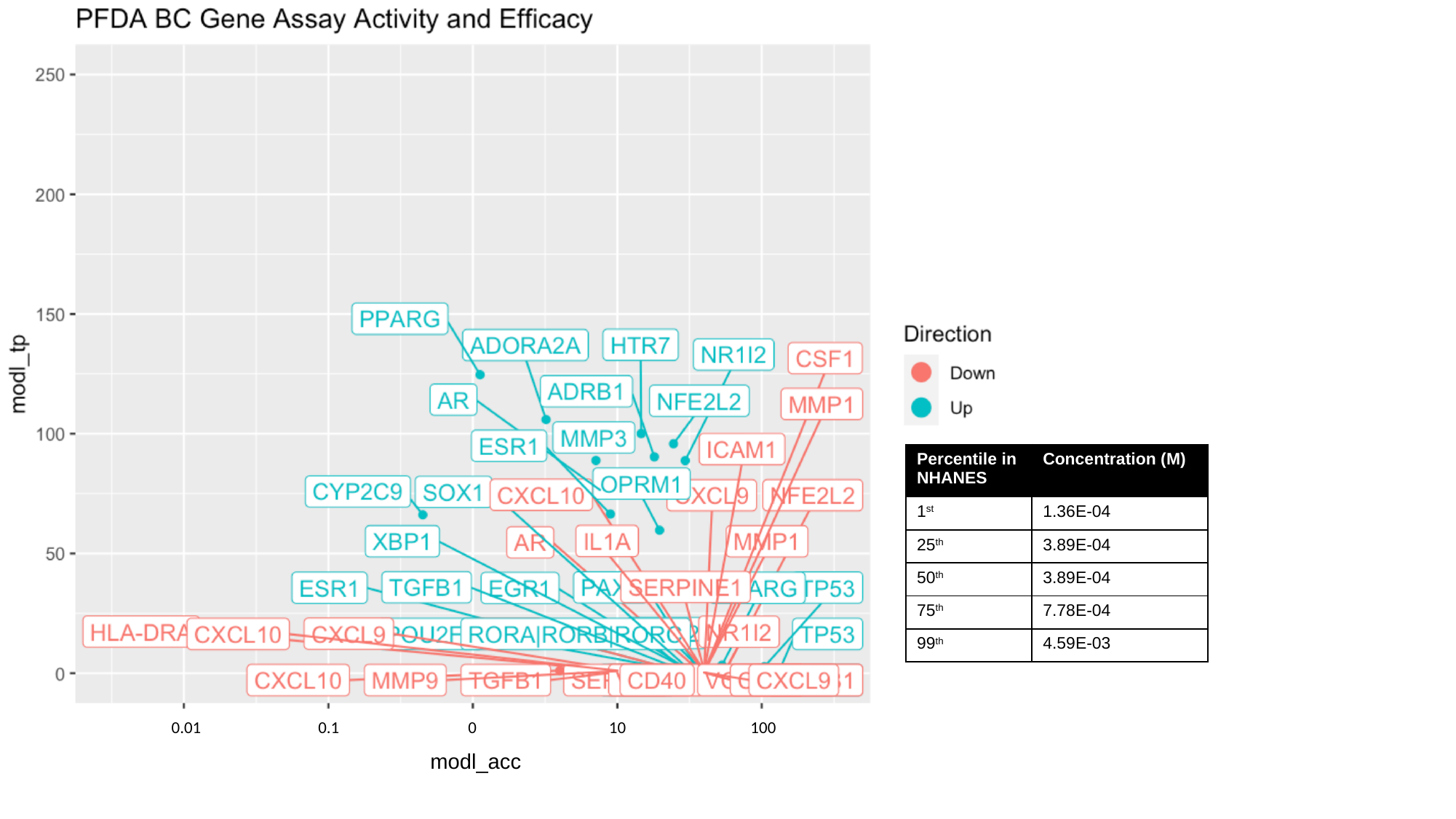

100
0.01
0.1
0
10
modl_acc

### Slide 18
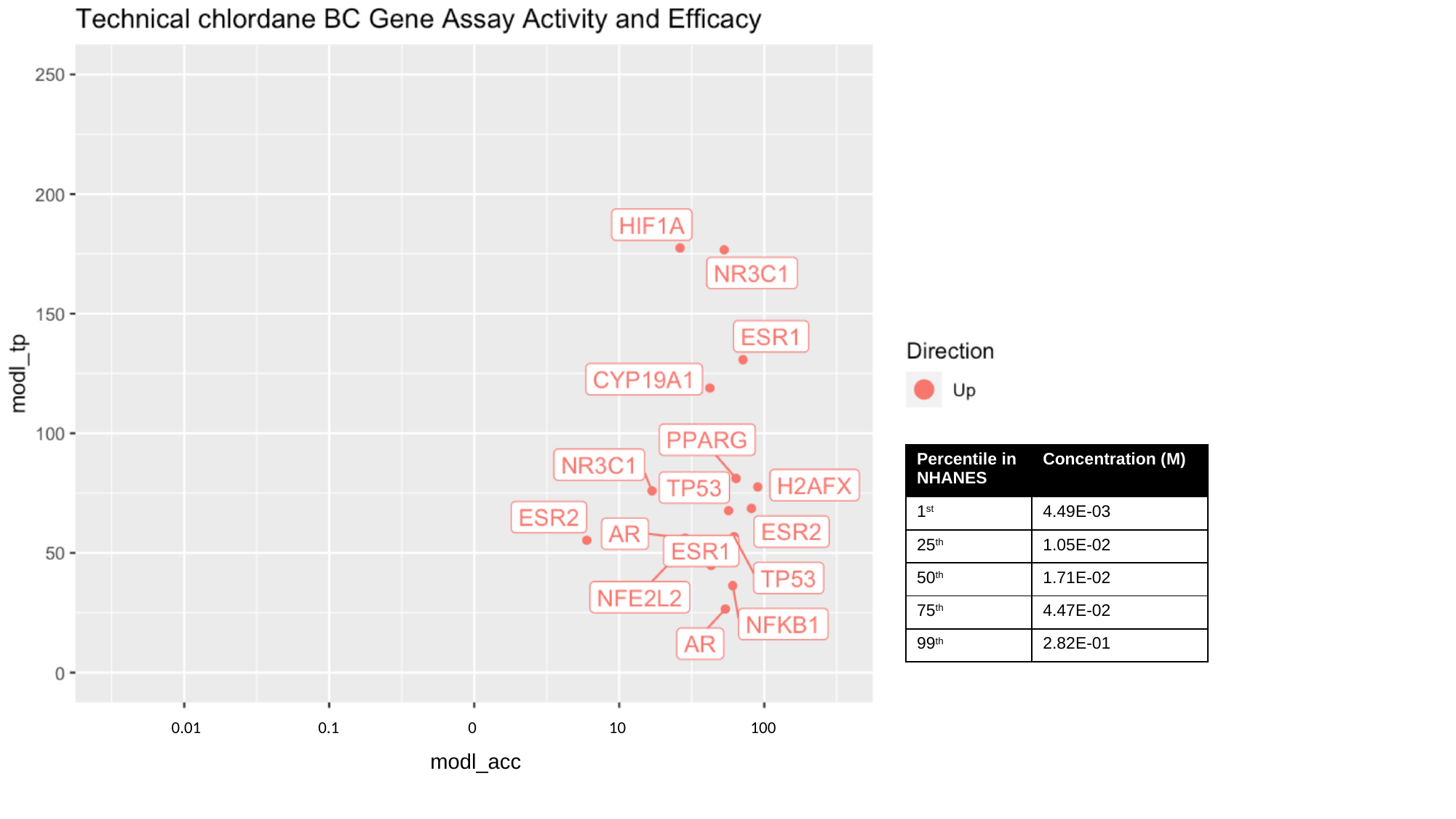

100
0.01
0.1
0
10
modl_acc

### Slide 19
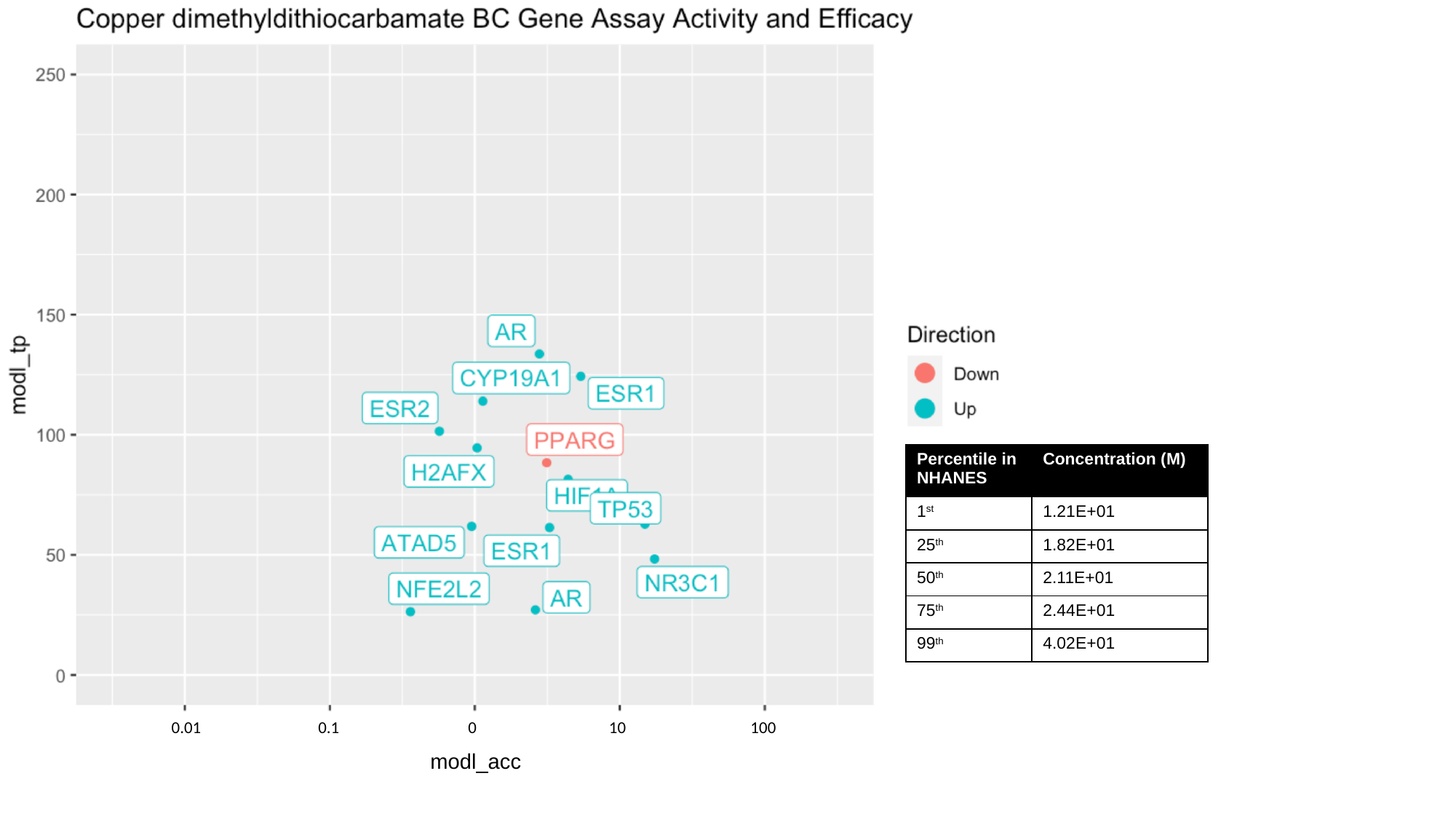

100
0.01
0.1
0
10
modl_acc

### Slide 20
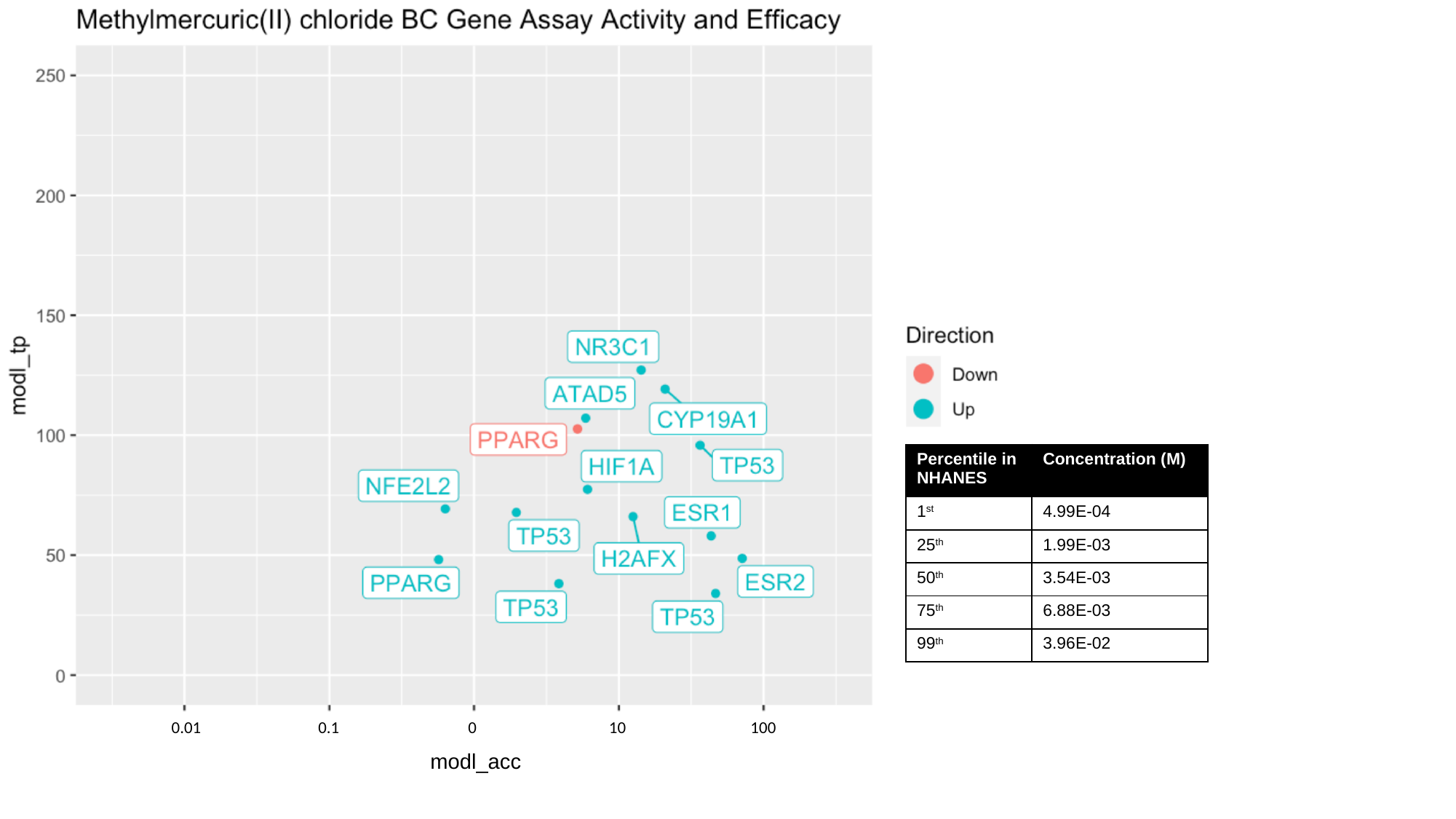

100
0.01
0.1
0
10
modl_acc

### Slide 21
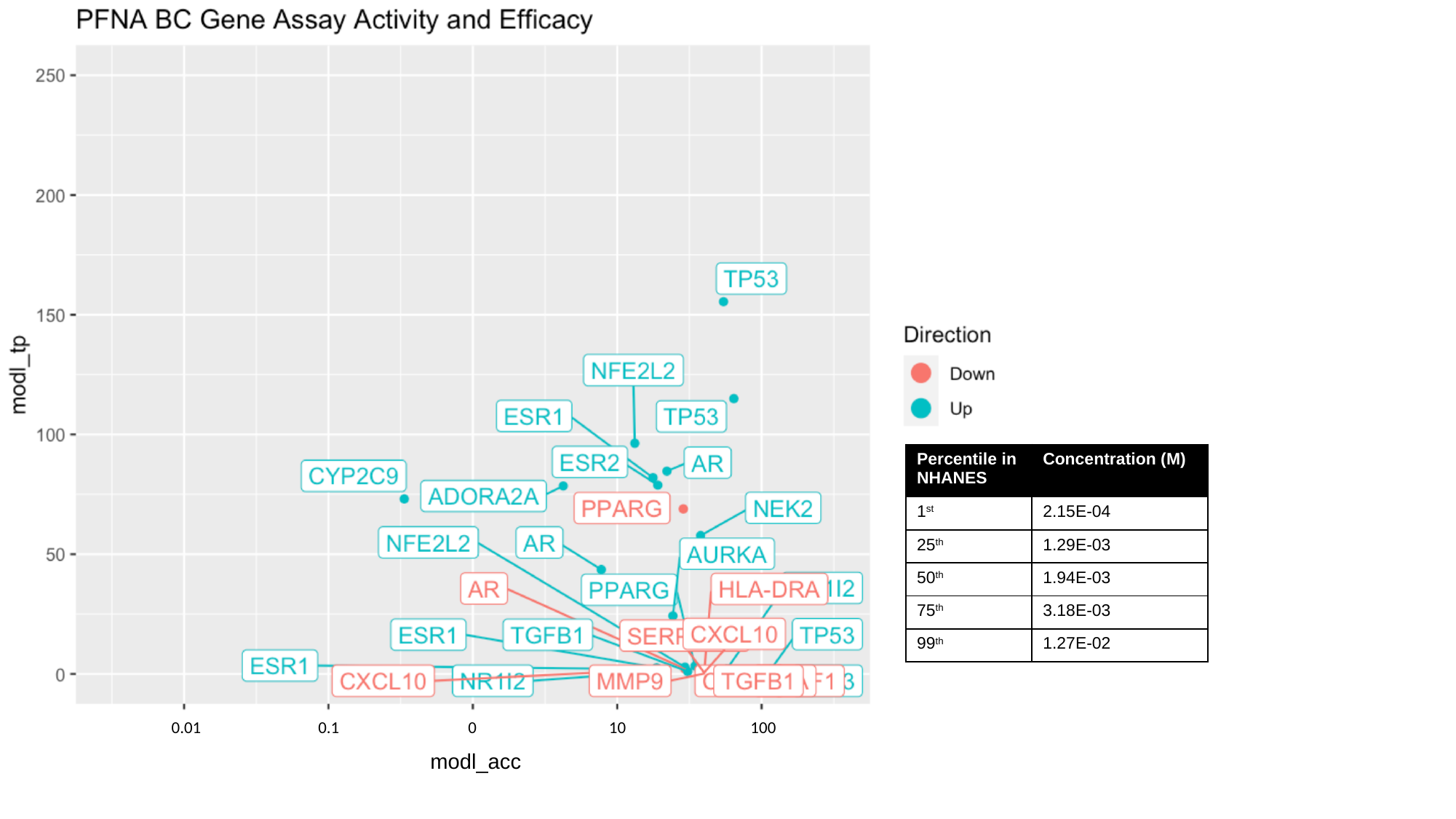

100
0.01
0.1
0
10
modl_acc

### Slide 22
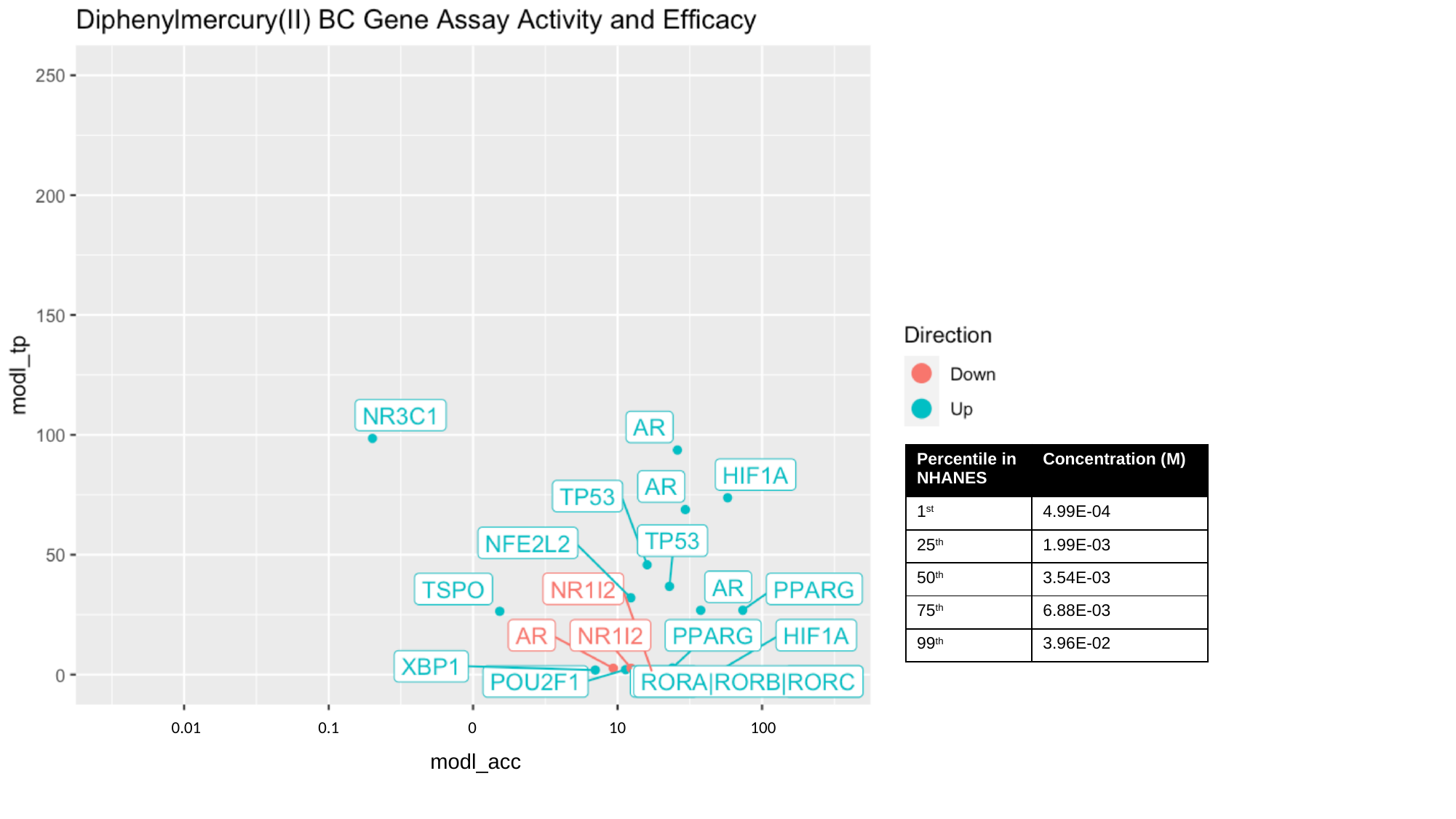

100
0.01
0.1
0
10
modl_acc

### Slide 23
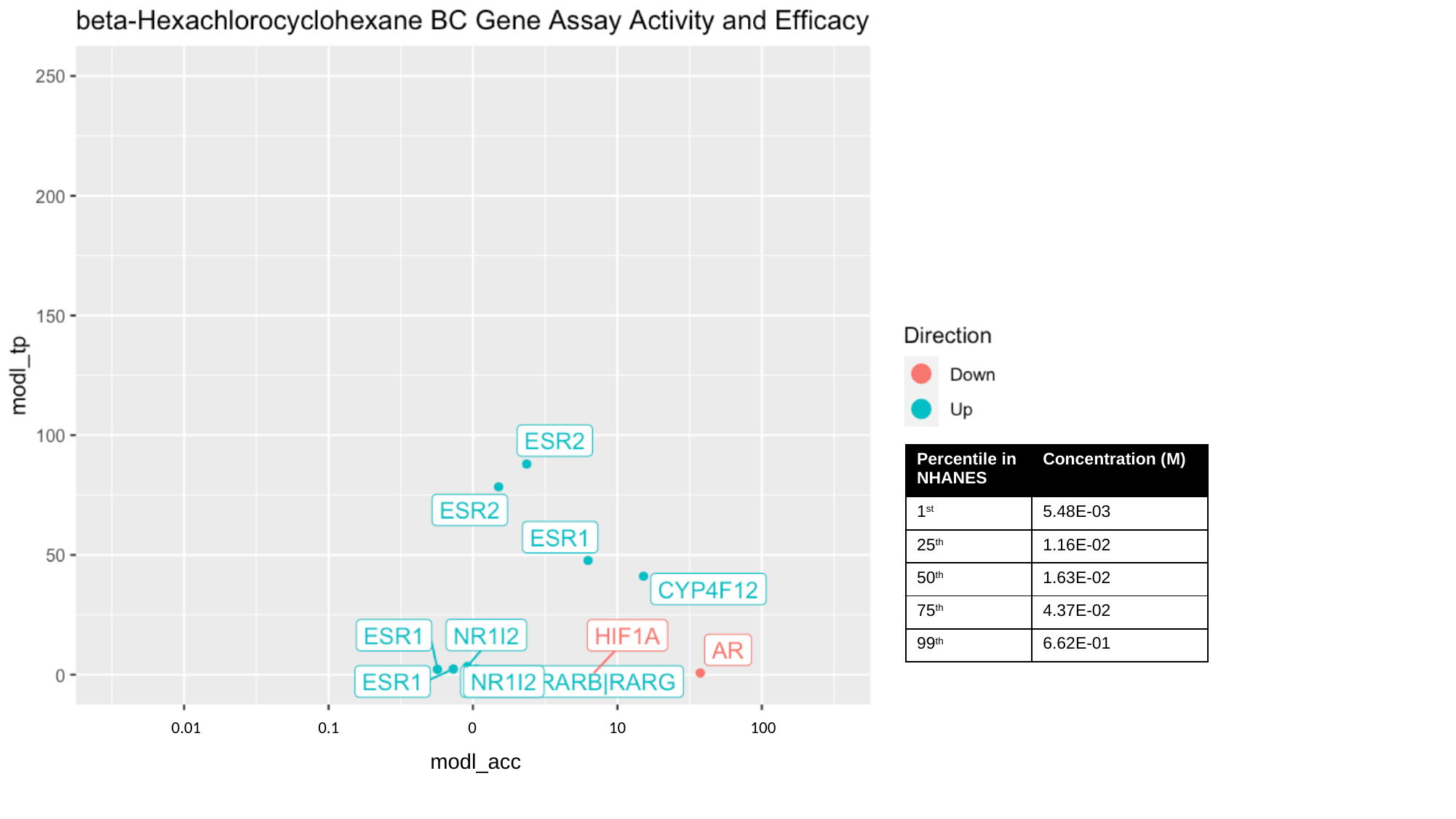

100
0.01
0.1
0
10
modl_acc

### Slide 24
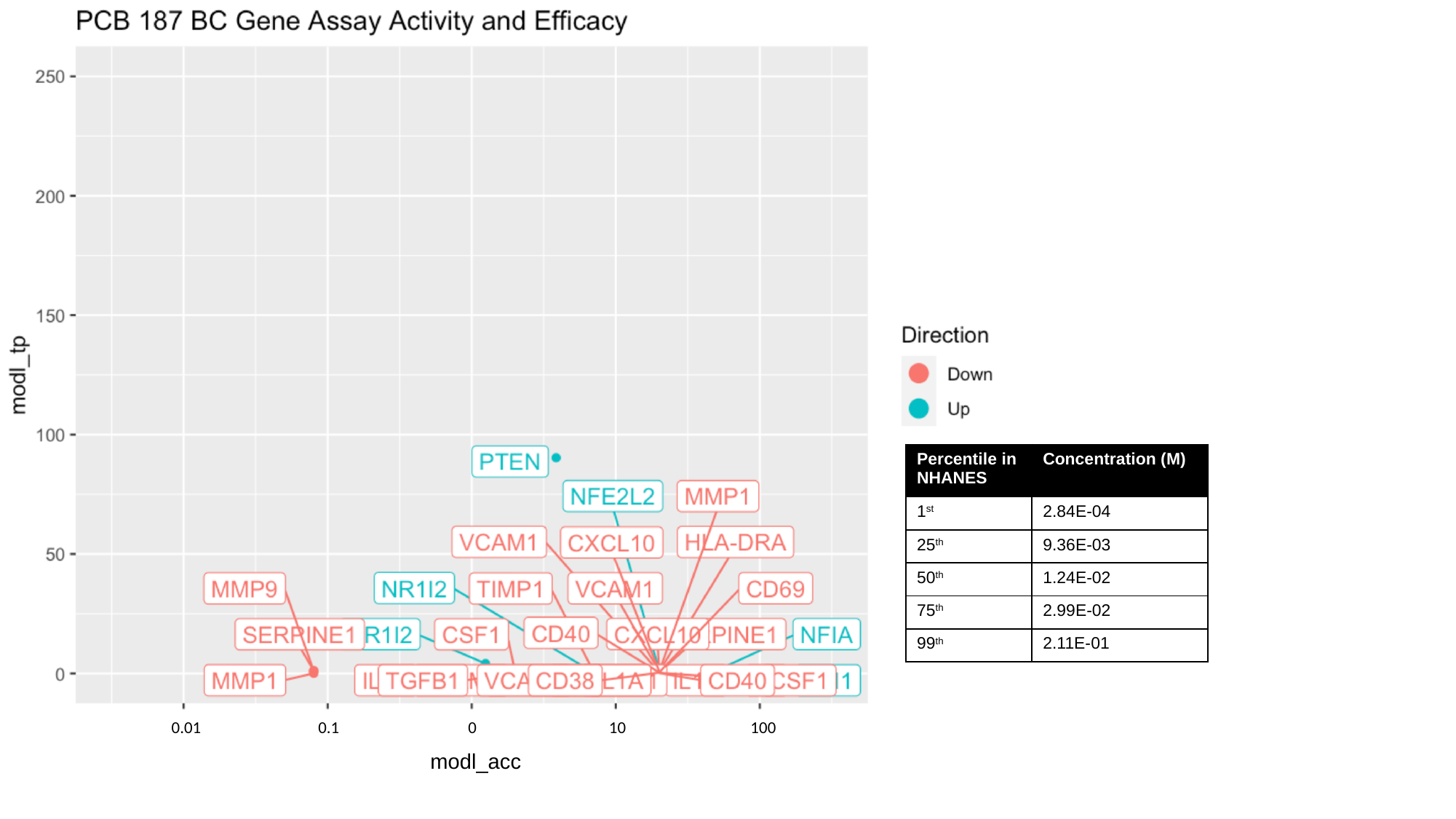

100
0.01
0.1
0
10
modl_acc

### Slide 25
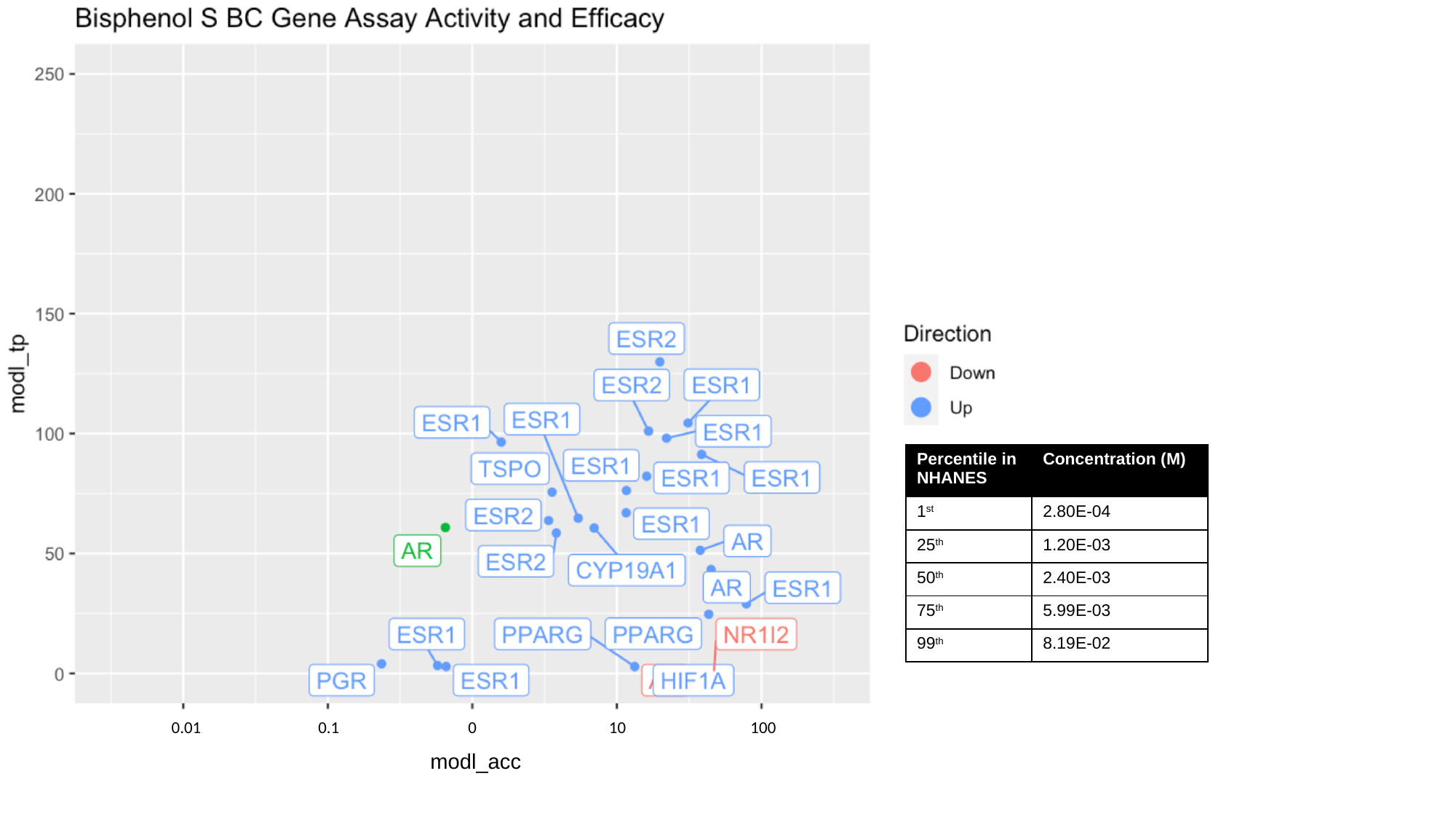

100
0.01
0.1
0
10
modl_acc

### Slide 26
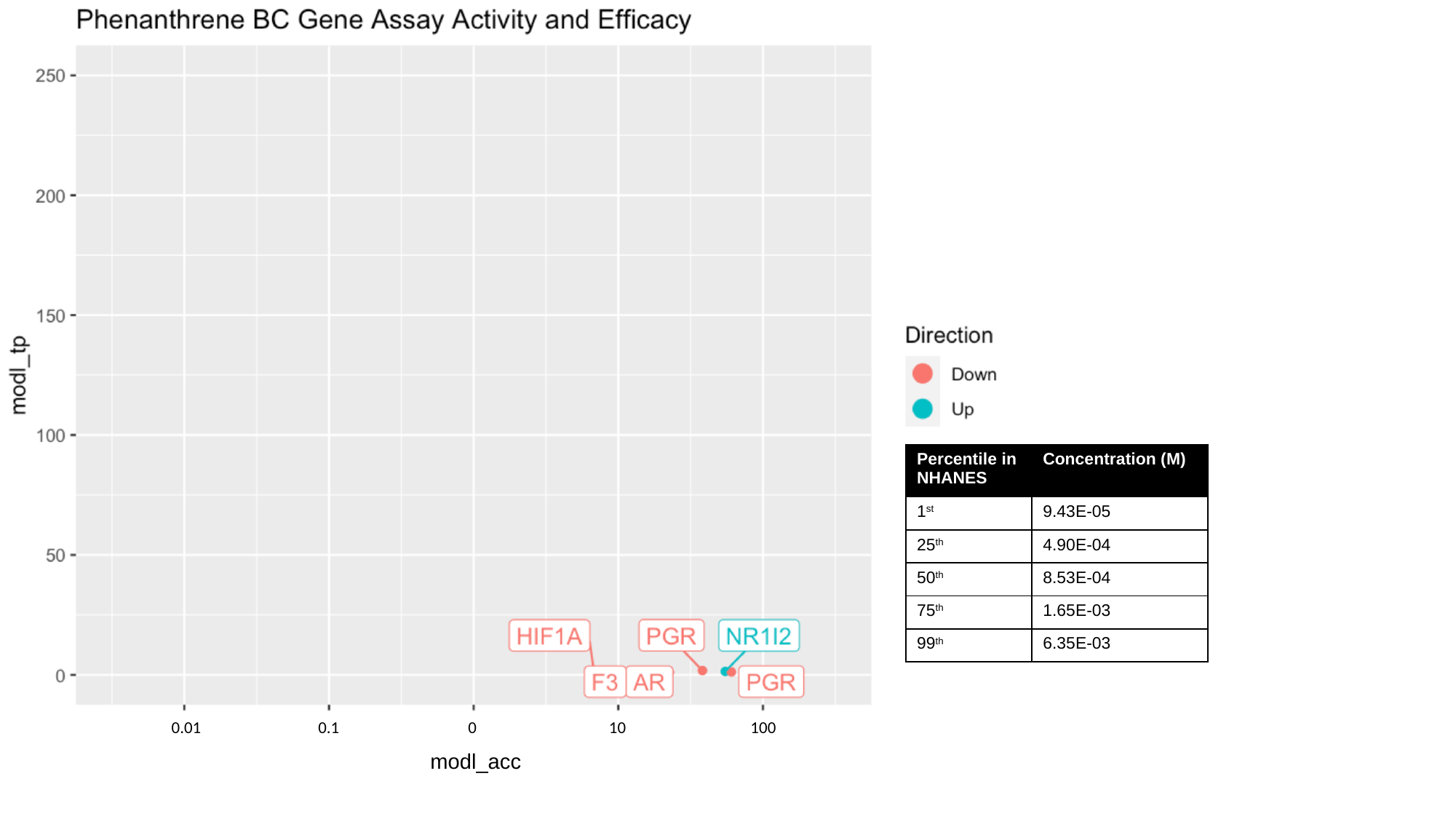

100
0.01
0.1
0
10
modl_acc

### Slide 27
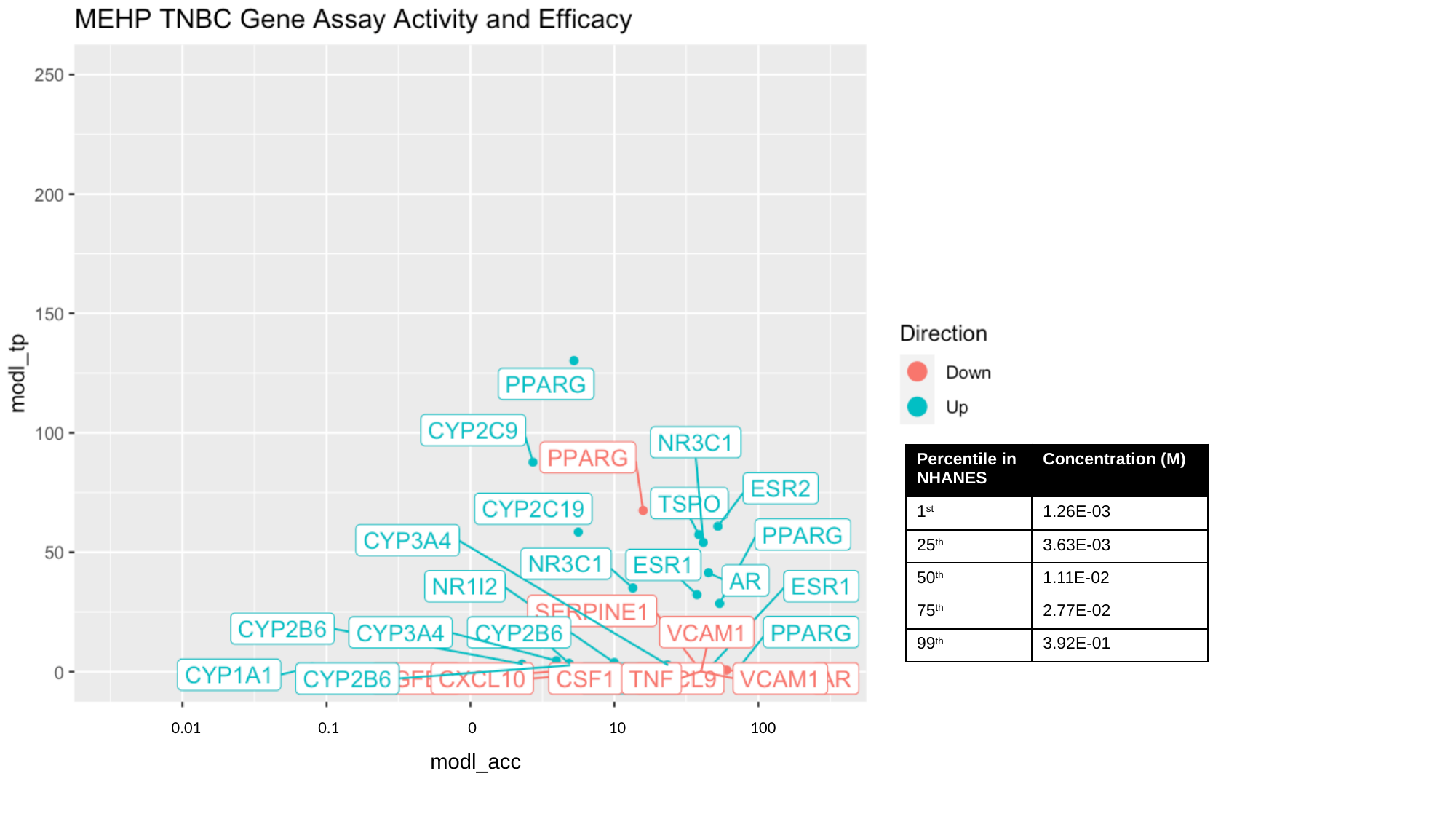

100
0.01
0.1
0
10
modl_acc

### Slide 28
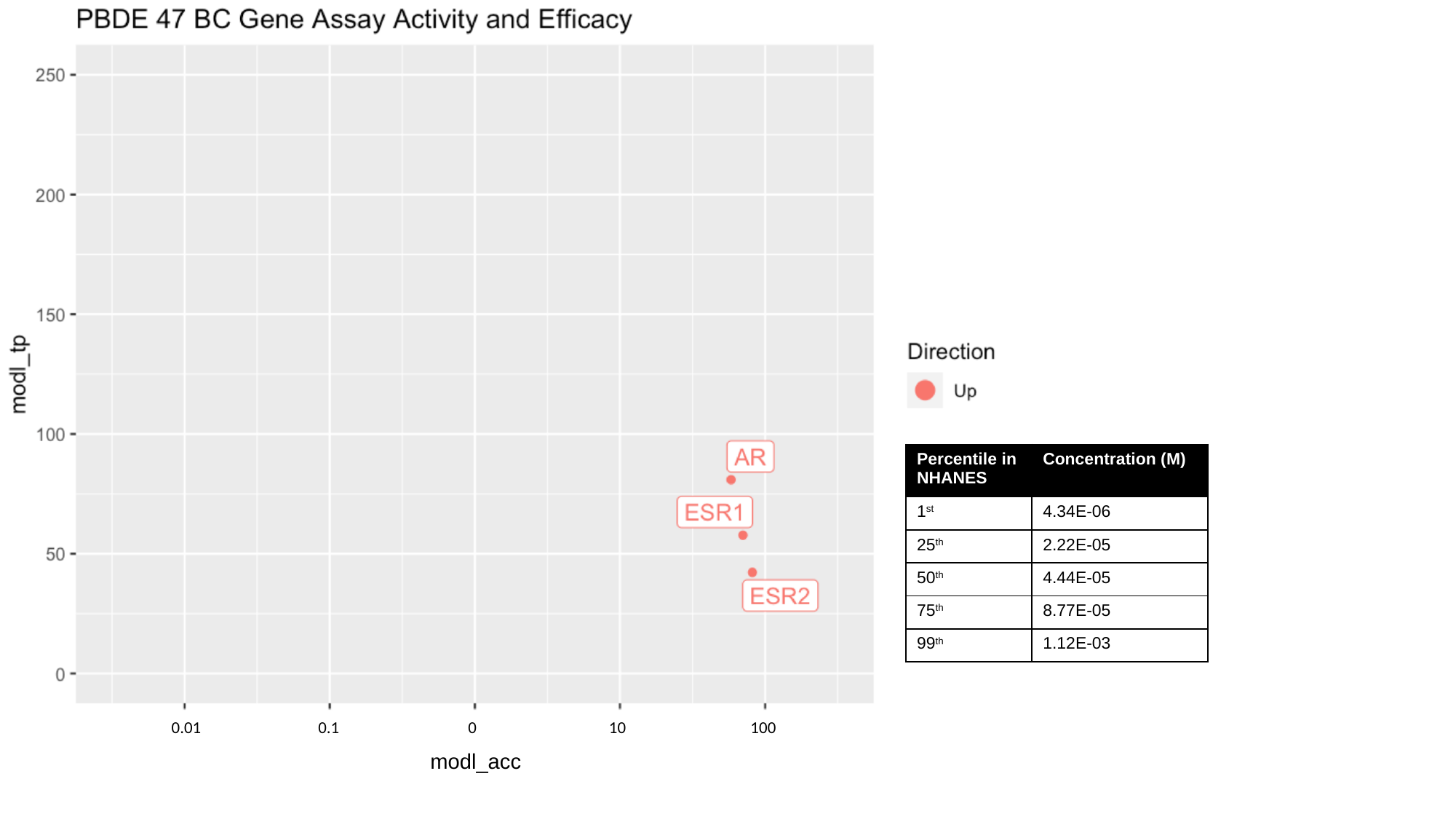

100
0.01
0.1
0
10
modl_acc

### Slide 29
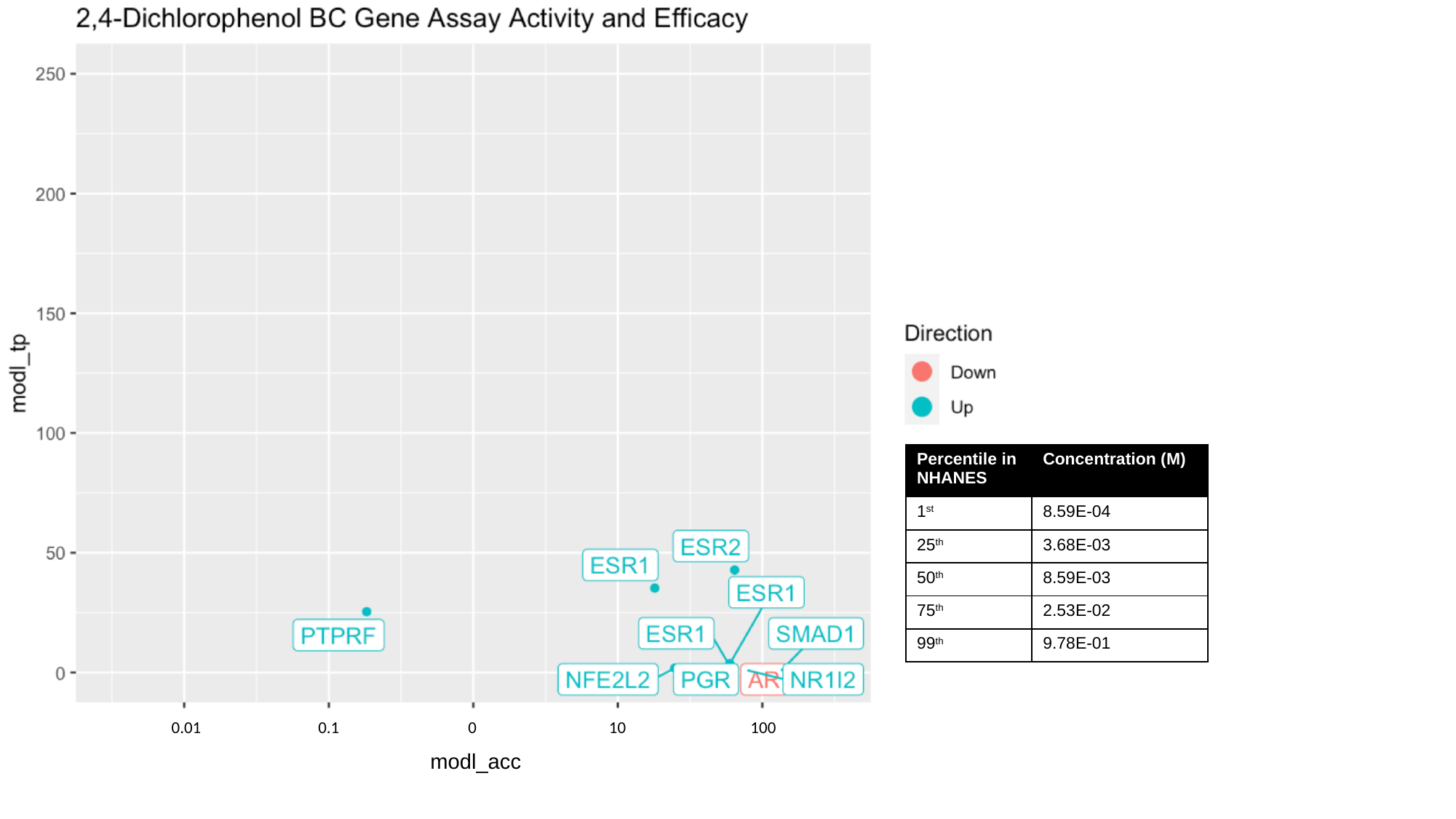

100
0.01
0.1
0
10
modl_acc

### Slide 30
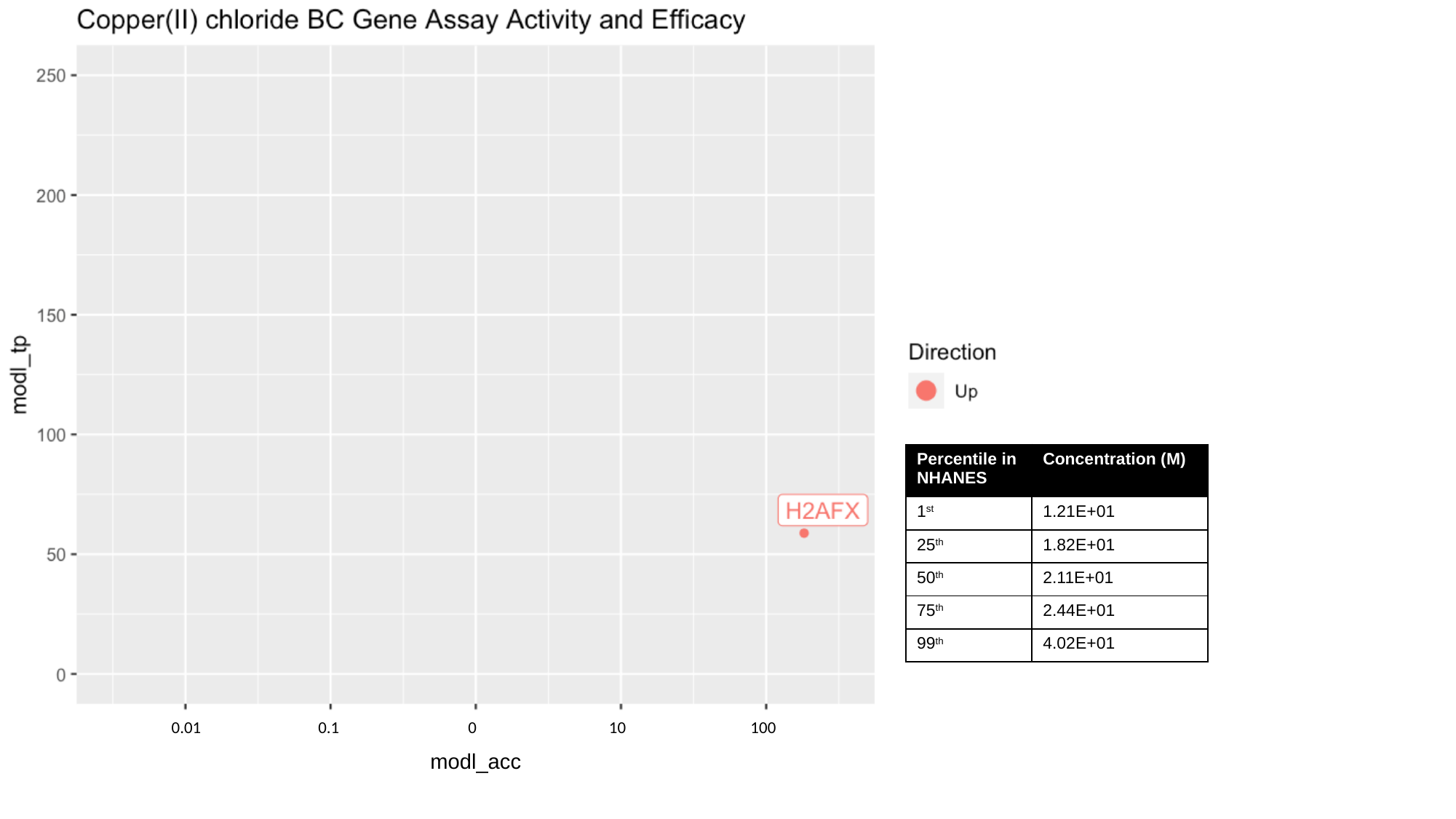

100
0.01
0.1
0
10
modl_acc

### Slide 31
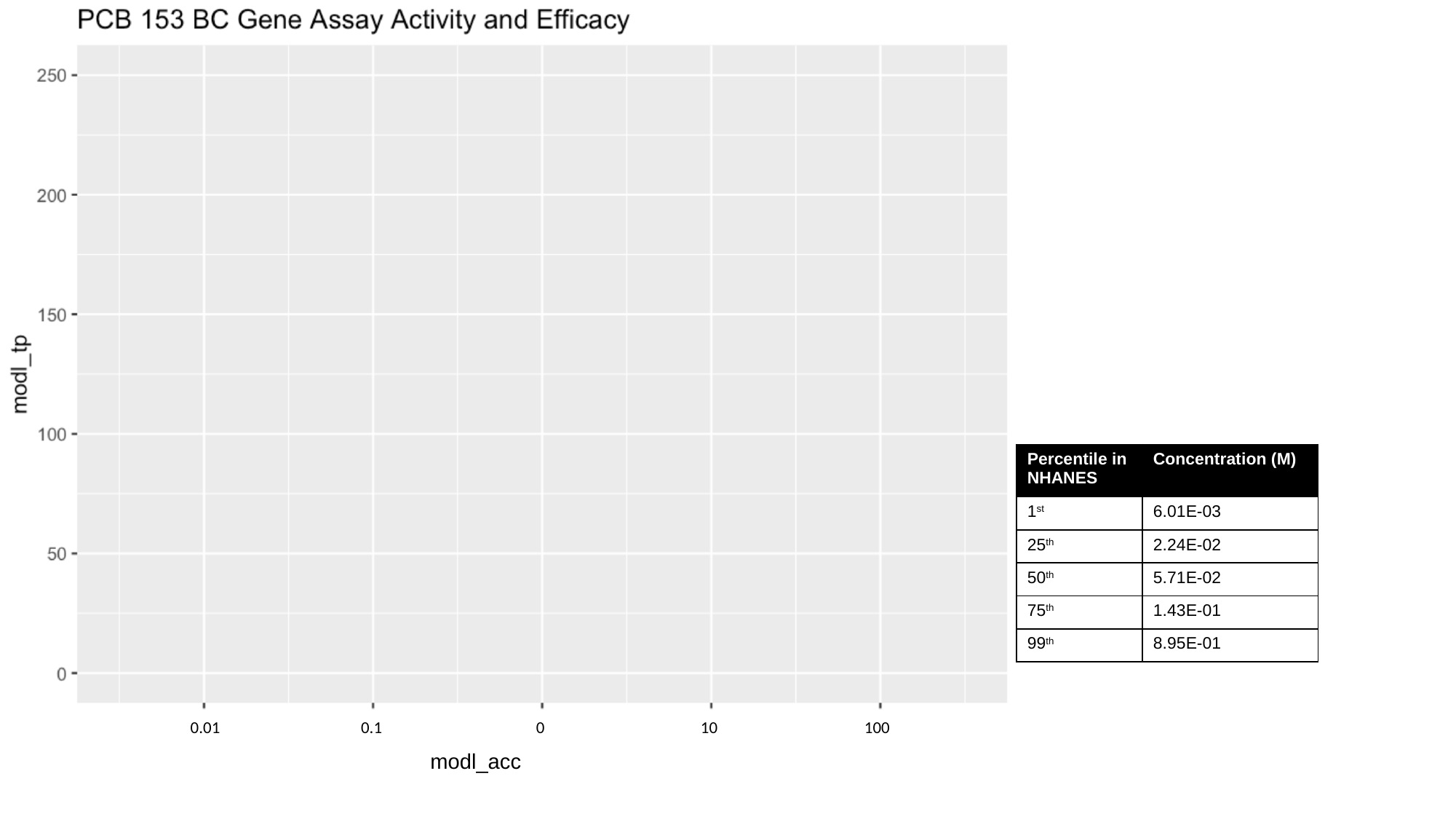

100
0.01
0.1
0
10
modl_acc

### Slide 32
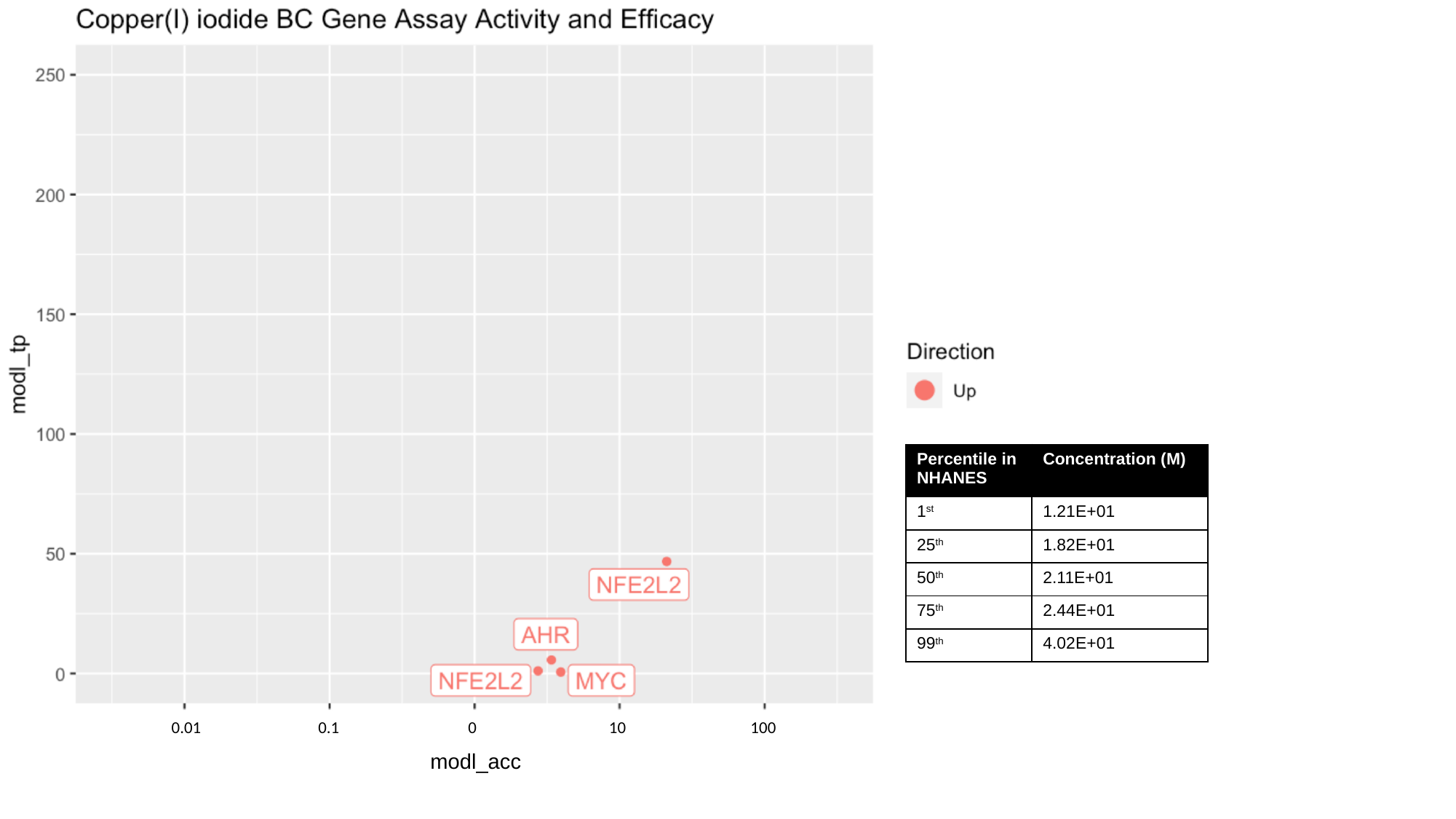

100
0.01
0.1
0
10
modl_acc

### Slide 33
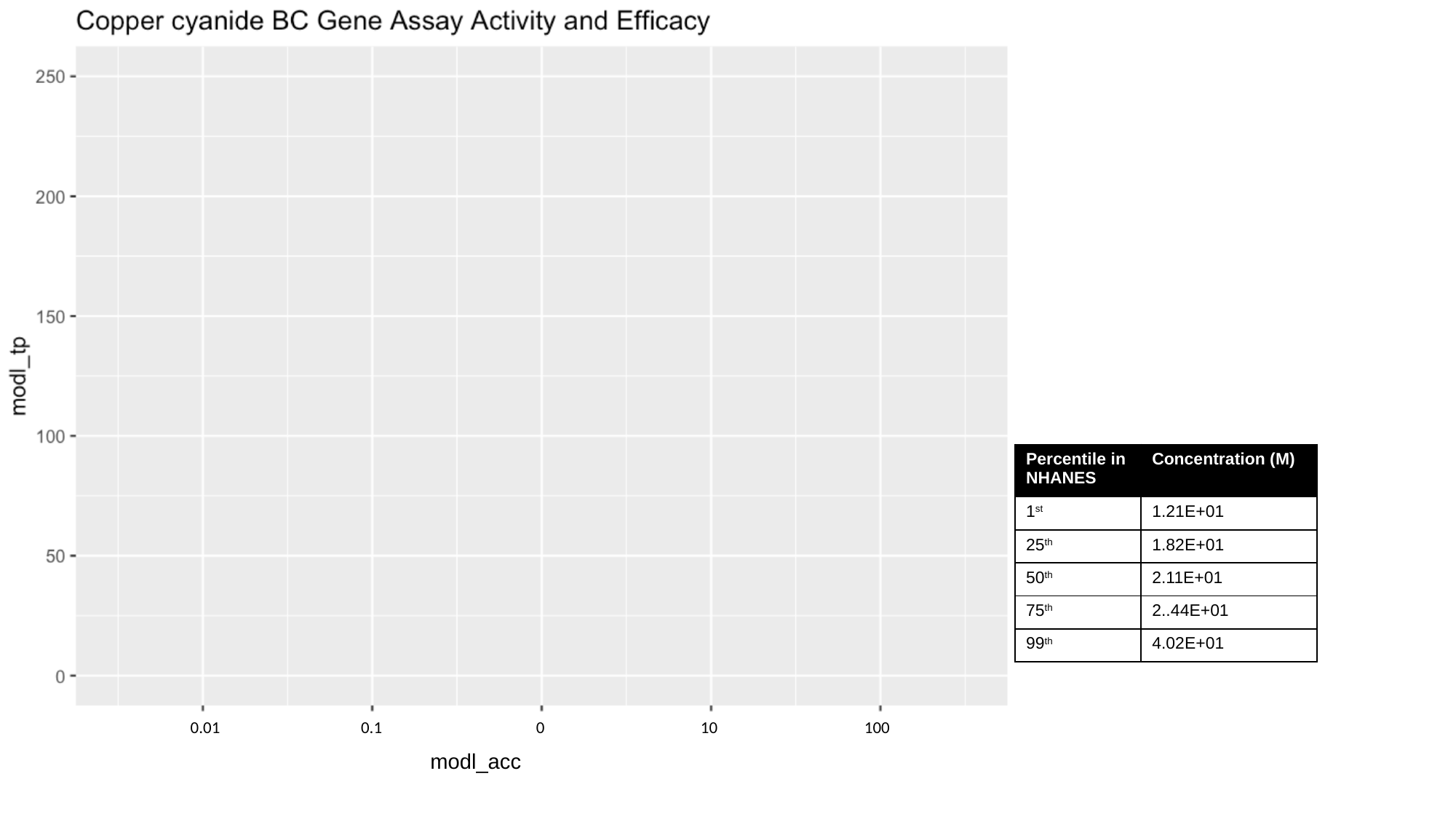

100
0.01
0.1
0
10
modl_acc

### Slide 34
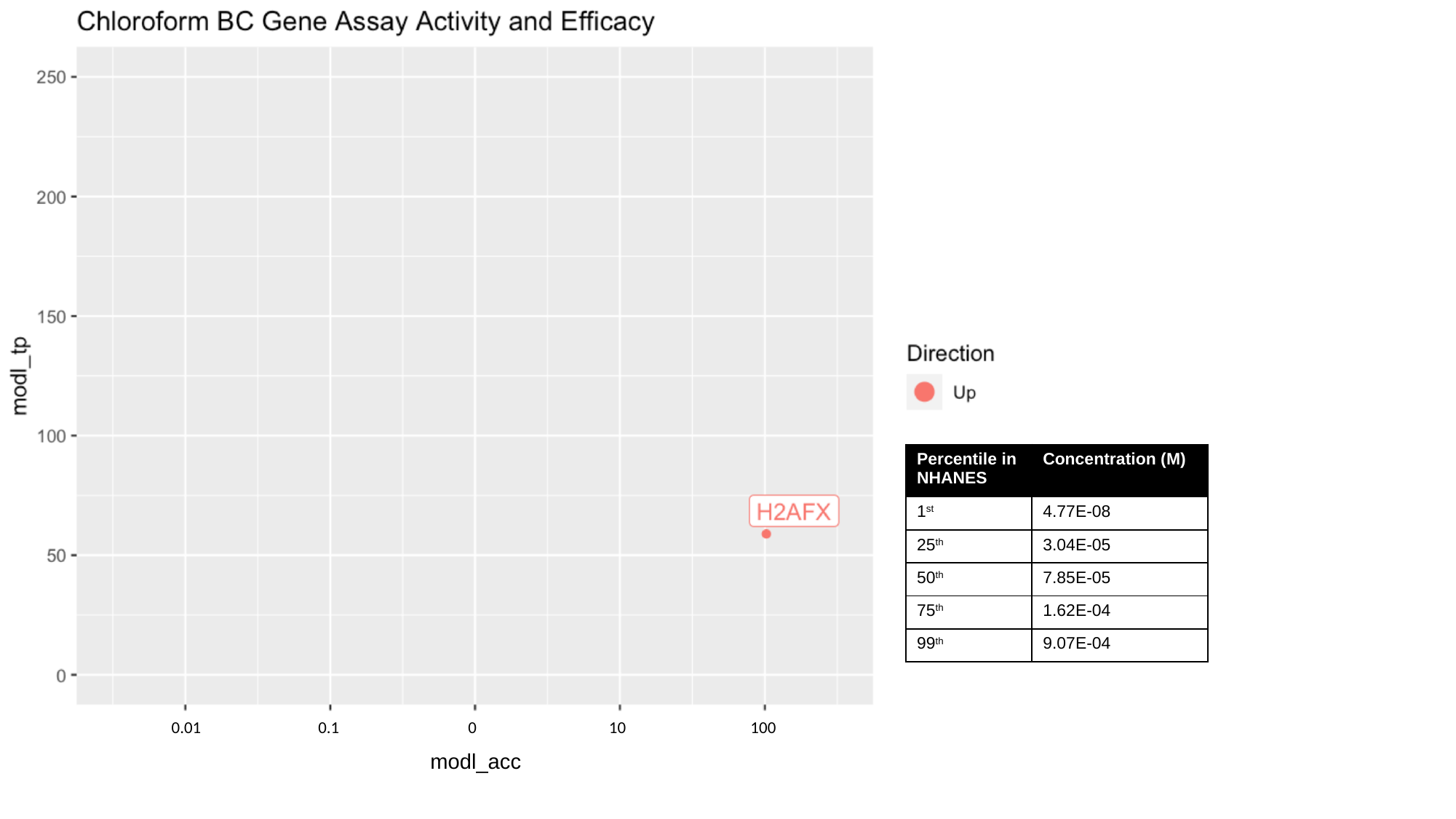

100
0.01
0.1
0
10
modl_acc

### Slide 35
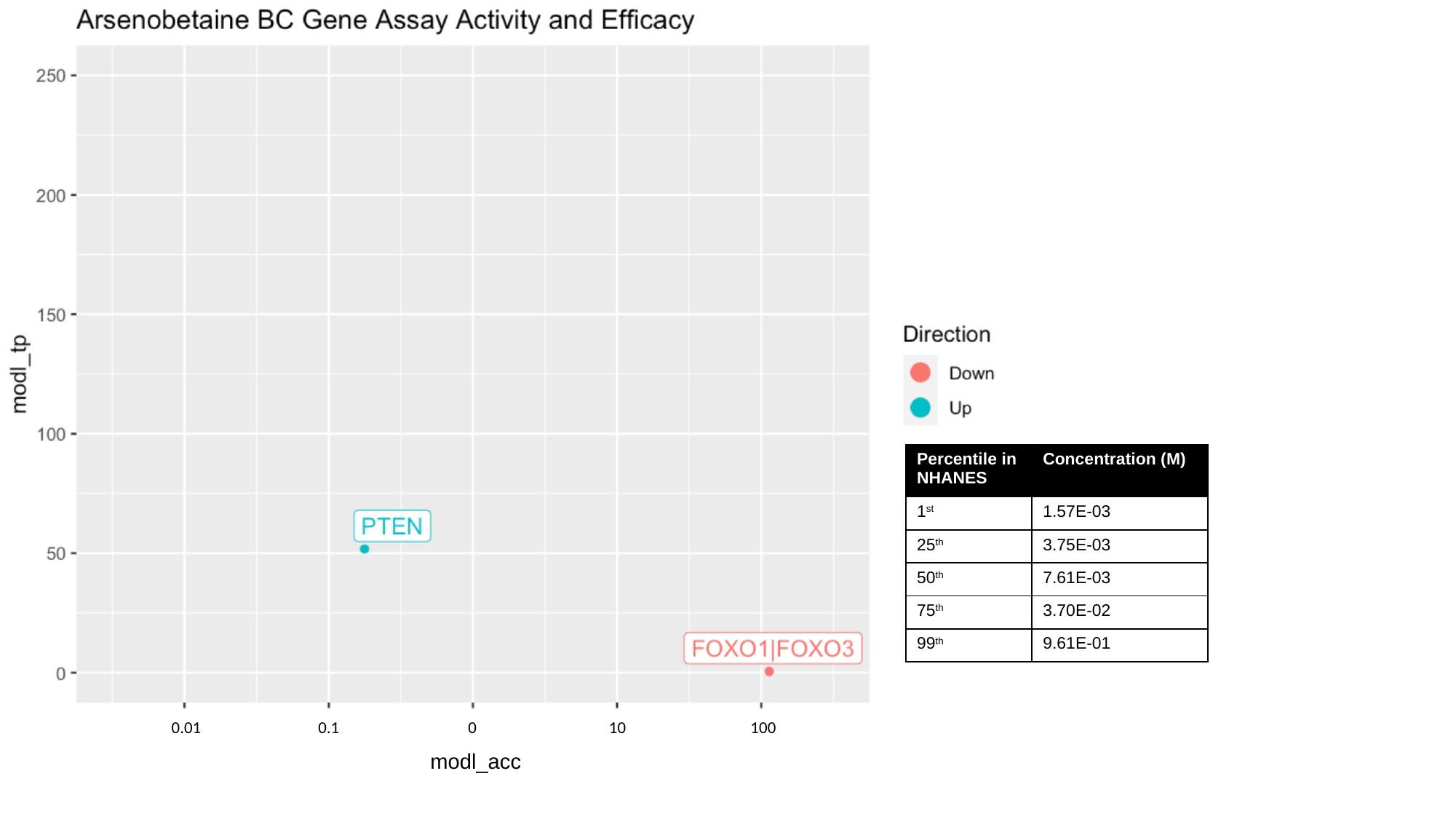

100
0.01
0.1
0
10
modl_acc

### Slide 36
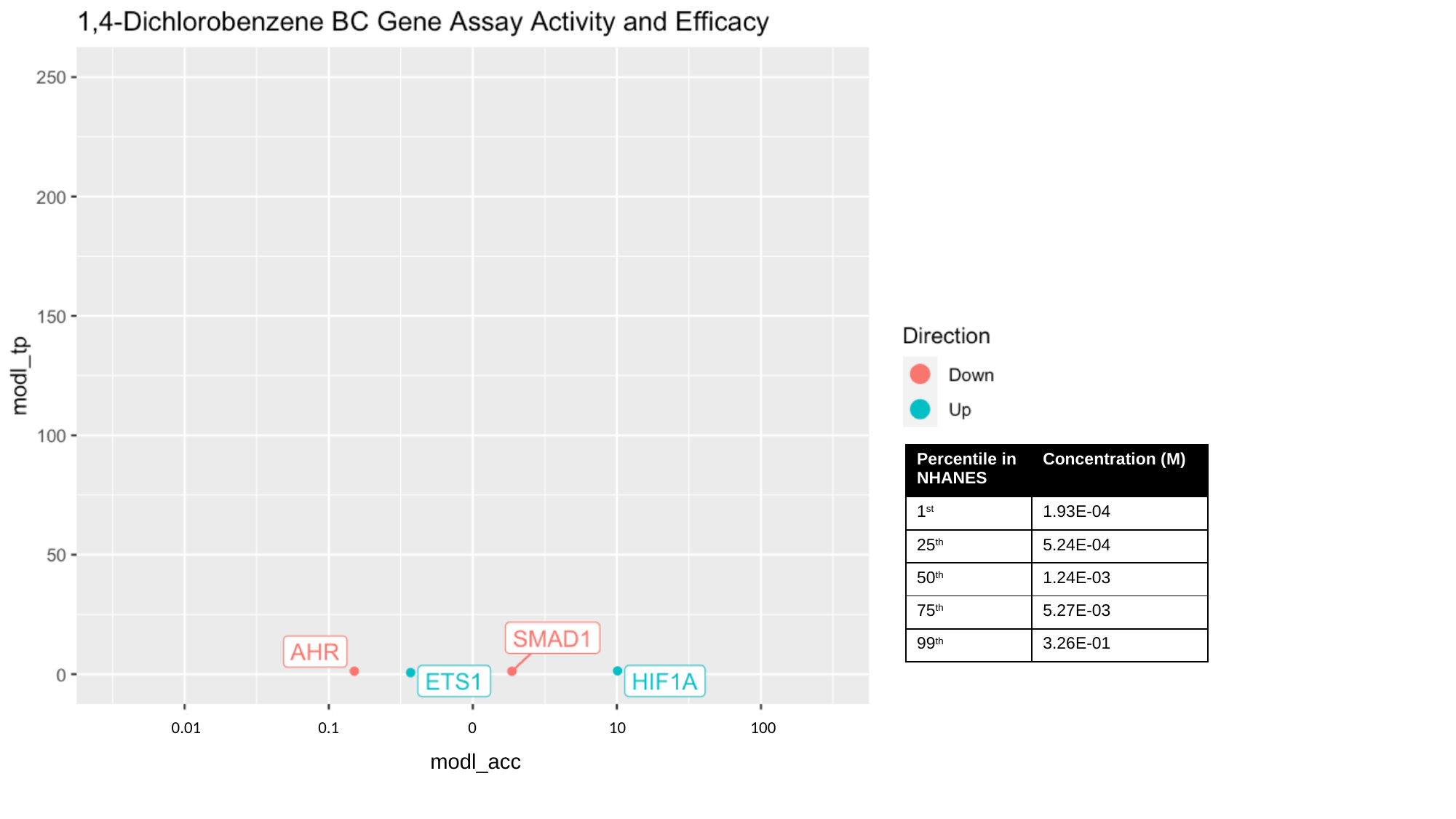

100
0.01
0.1
0
10
modl_acc

### Slide 37
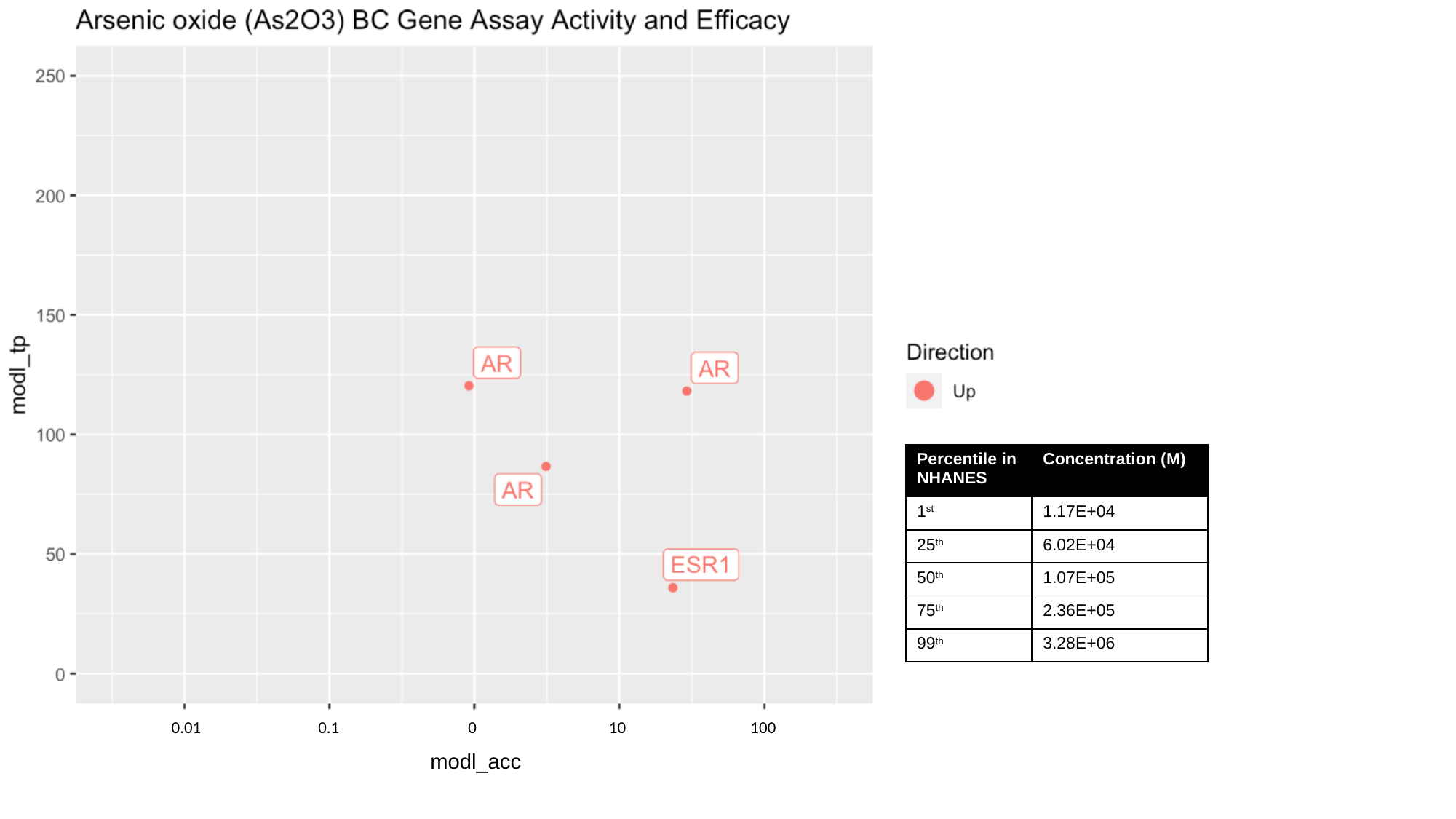

100
0.01
0.1
0
10
modl_acc

### Slide 38
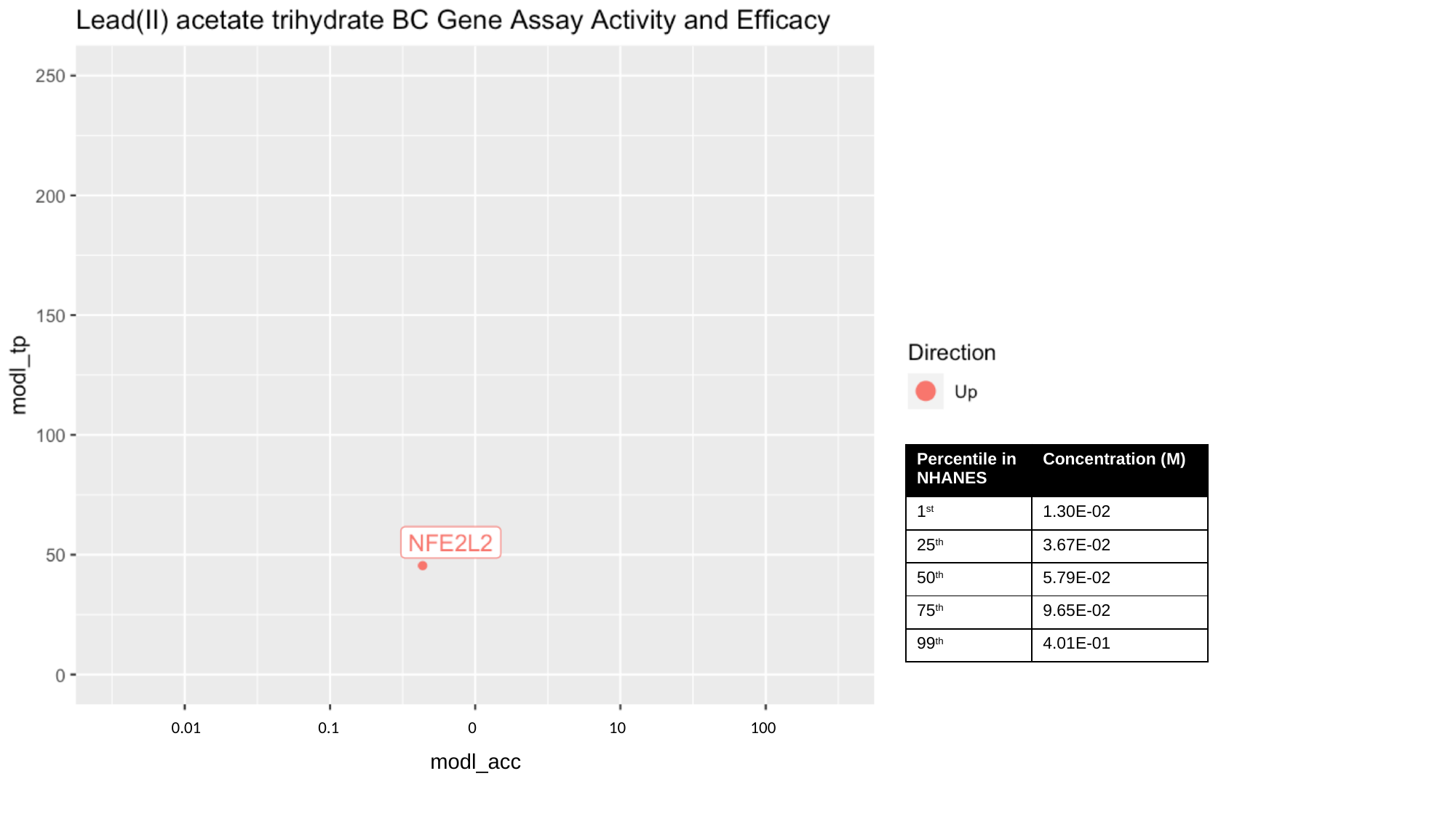

100
0.01
0.1
0
10
modl_acc

### Slide 39
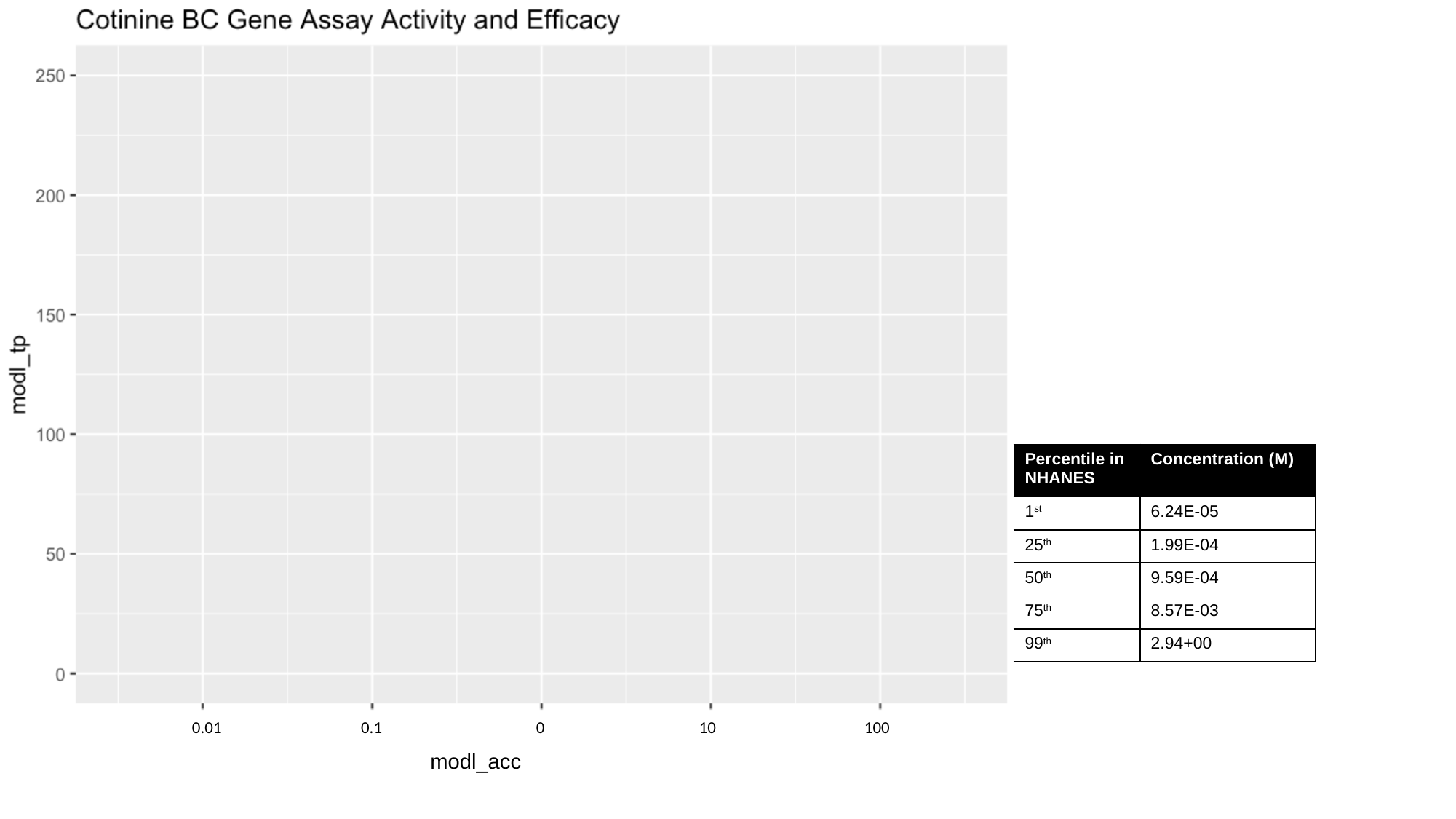

100
0.01
0.1
0
10
modl_acc

### Slide 40
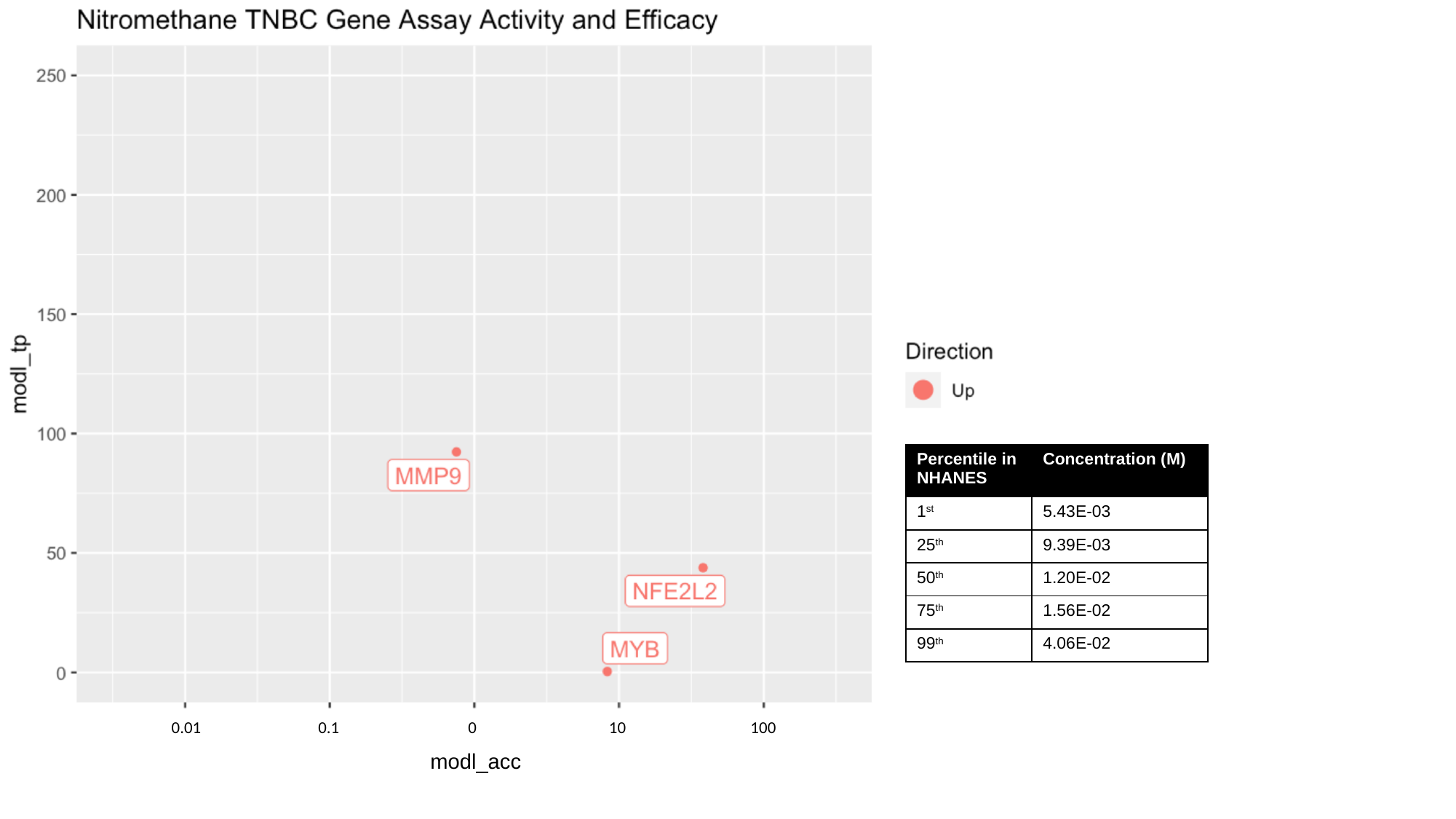

100
0.01
0.1
0
10
modl_acc

### Slide 41
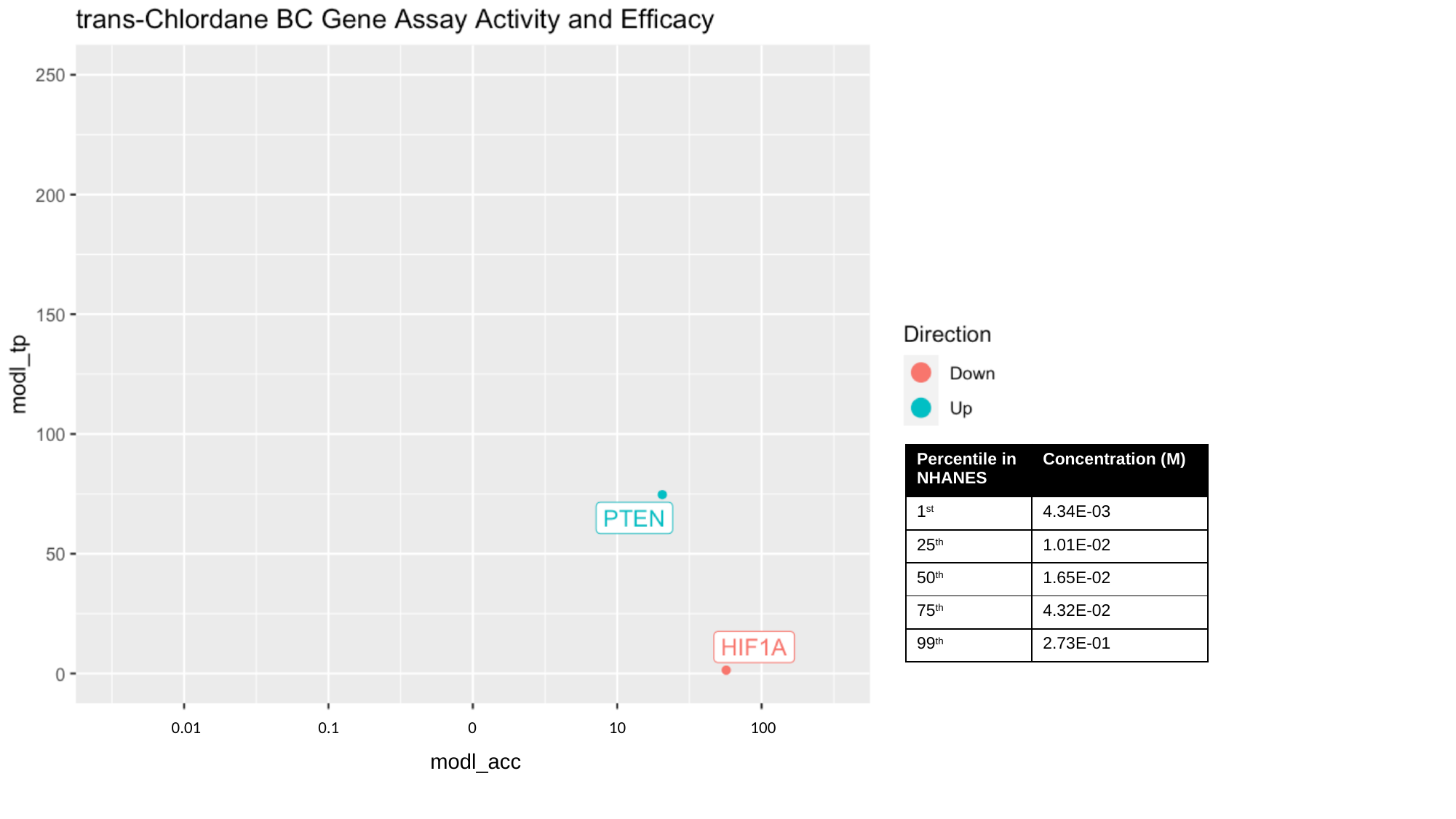

100
0.01
0.1
0
10
modl_acc

### Slide 42
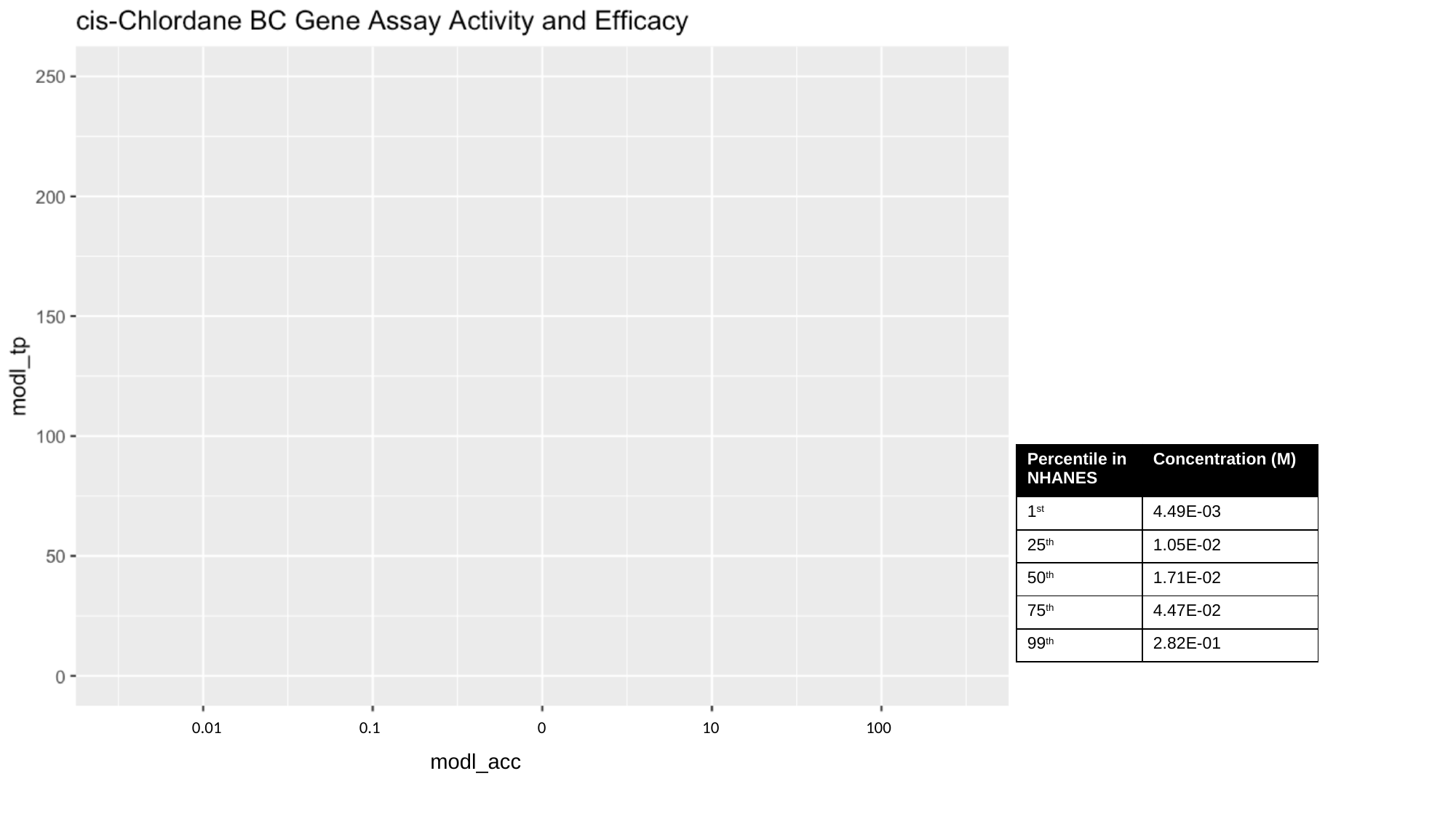

100
0.01
0.1
0
10
modl_acc

### Slide 43
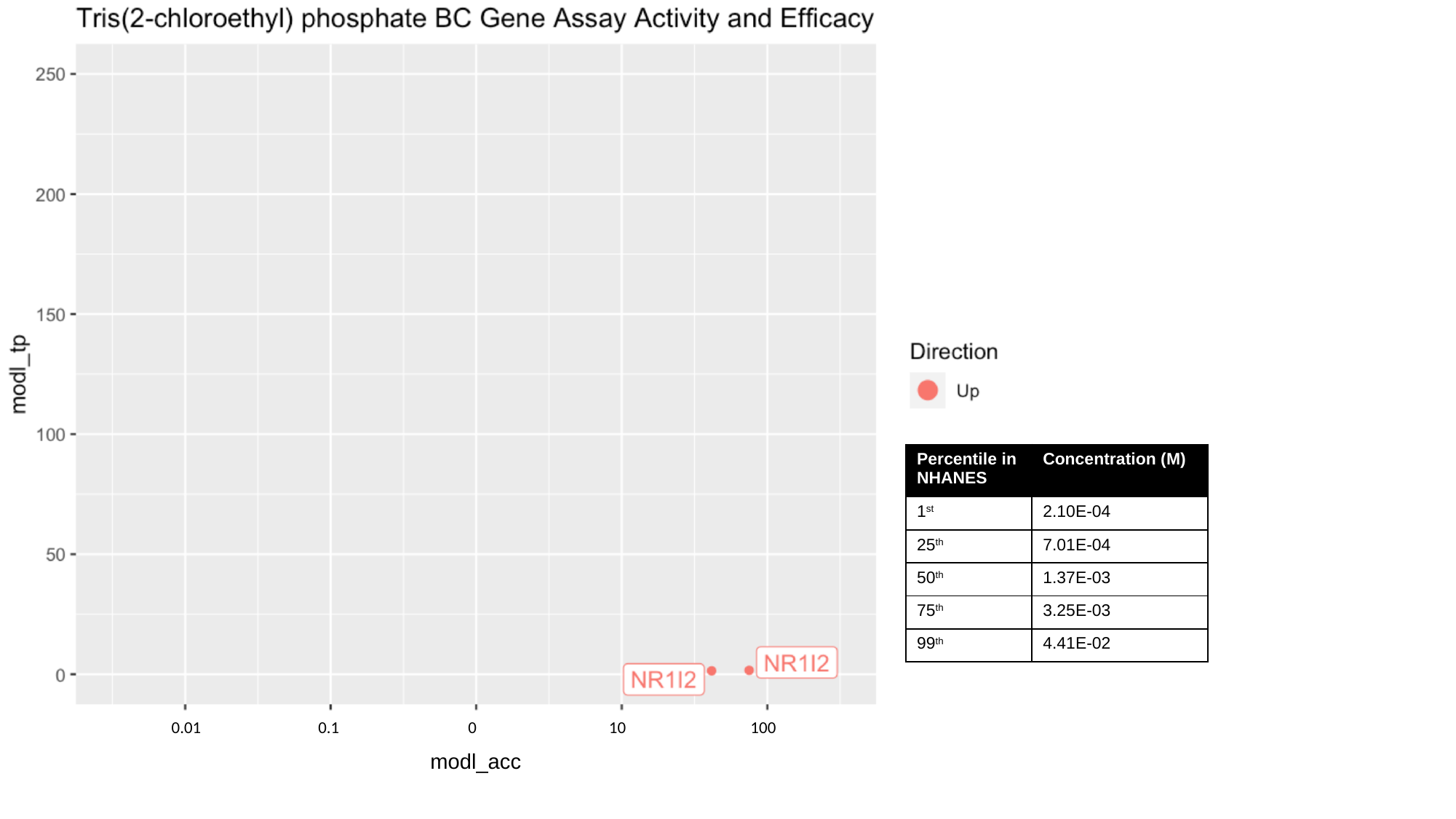

100
0.01
0.1
0
10
modl_acc

### Slide 44
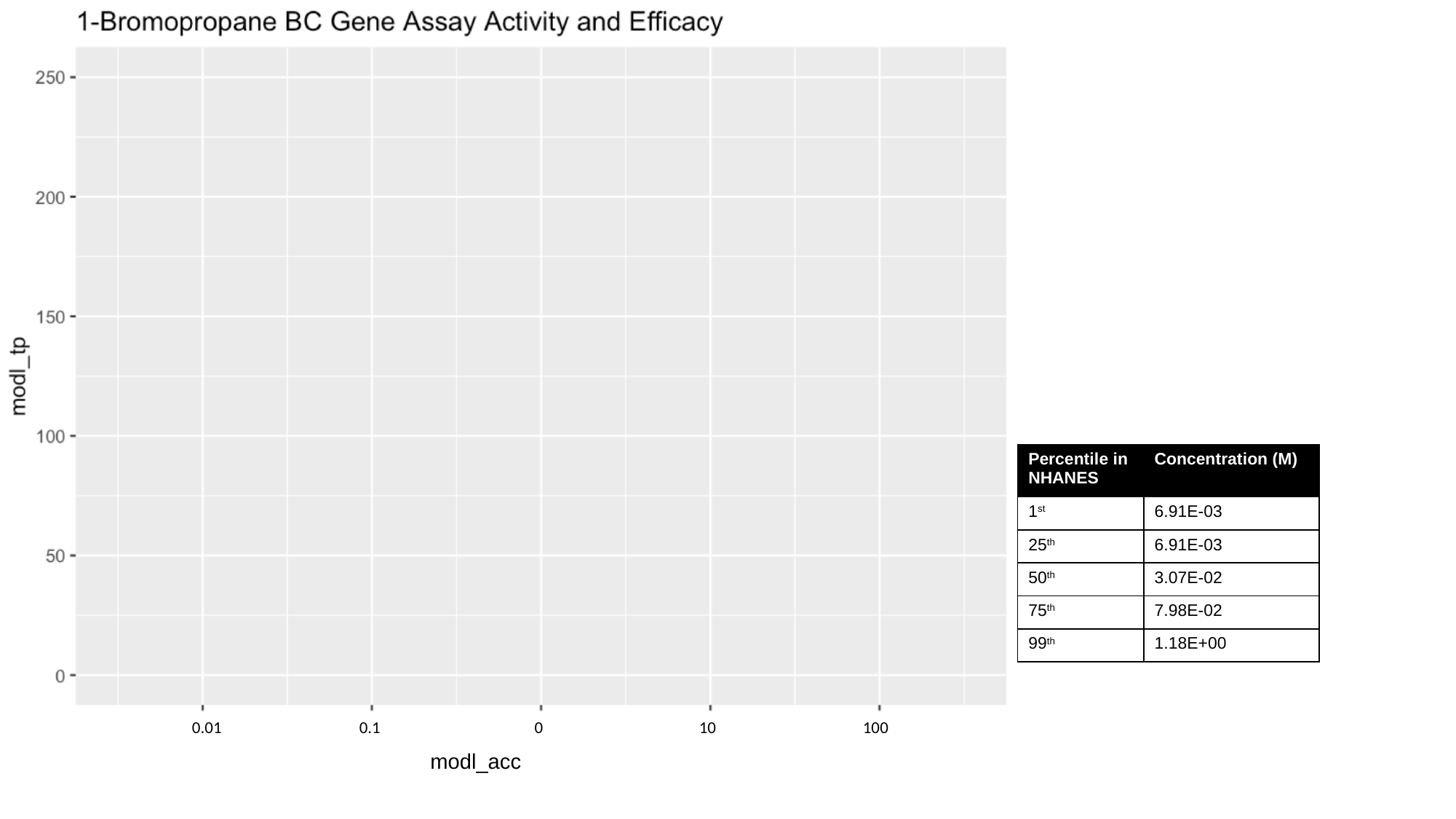

100
0.01
0.1
0
10
modl_acc
